# Stable Species Boundaries Despite Ten Million Years of Hybridization in Tropical Eels

**DOI:** 10.1101/635631

**Authors:** Julia M. I. Barth, Chrysoula Gubili, Michael Matschiner, Ole K. Tørresen, Shun Watanabe, Bernd Egger, Yu-San Han, Eric Feunteun, Ruben Sommaruga, Robert Jehle, Robert Schabetsberger

## Abstract

Genomic evidence is increasingly underpinning that hybridization between taxa is commonplace, challenging our views on the mechanisms that maintain their boundaries. Here, we focus on seven catadromous eel species (genus *Anguilla*), and use genome-wide sequence data from more than 450 individuals sampled across the tropical Indo-Pacific, morphological information, and three newly assembled draft genomes to compare contemporary patterns of hybridization with signatures of past gene flow across a time-calibrated phylogeny. We show that the seven species have remained distinct entities for up to 10 million years, despite a dynamic scenario of incomplete isolation whereby the current frequencies of hybridization across species pairs (over 5% of all individuals were either F1 hybrids or backcrosses) contrast remarkably with patterns of past introgression. Based on near-complete asymmetry in the directionality of hybridization and decreasing frequencies of later-generation hybrids, we identify cytonuclear incompatibilities and hybrid breakdown as two powerful mechanisms that can support species cohesion even when hybridization has been pervasive throughout the evolutionary history of entire clades.

## Introduction

The turn of the century has witnessed a paradigm shift in how we view the role of hybridization for building up biological diversity. While hybridization was previously assumed to be spatially restricted and confined to a small number of taxa, it became gradually recognized that incomplete isolation of genomes is widespread across eukaryotes, with varied effects on adaptation and speciation (Mallet, 2005, 2007; Abbott et al., 2013; Taylor and Larson, 2019). More recently, this view has been further fuelled by technical and analytical advances which enable the quantification of past introgression, the genetic exchange through hybridization, across entire clades, revealing that it is often the most rapidly radiating clades that experienced high frequencies of gene exchange (Meier et al., 2017; Lamichhaney et al., 2018; Kozak et al., 2018; Edelman et al., 2018). This seemingly paradoxical association between introgression and rapid species proliferation underlies a key question in evolutionary biology: How can species in diversifying clades be accessible for introgression but nevertheless solidify their species boundaries? To answer this question, insights are required into the mechanisms that gradually reduce the degree to which hybridization generates introgression; however, these mechanisms are still poorly understood because contemporary hybridization and past introgression have so far not been jointly studied and compared across multiple pairs of animal species with different divergence times within a single clade.

Teleost fish provide well-established model systems to reveal processes of diversification, including the impact of hybridization on speciation (e.g., Malinsky et al., 2018a; Hench et al. 2019). A particularly promising system for hybridization research are catadromous freshwater eels of the genus *Anguilla*, one of the most species-rich genera of eels with high economic value (Nelson et al., 2016). These fishes are renowned for their unique population biology, whereby all individuals of a given species reproduce panmictically in one or only few oceanic spawning areas (Jacobsen et al., 2014; Pujolar and Maes, 2016). Moreover, spawning is temporally and spatially overlapping between multiple species, which therefore are expected to have great potential for interspecies mating (Avise et al. 1990; Schabetsberger et al. 2015). Frequent occurrence of hybridization has in fact been demonstrated with genomic data for the two Atlantic *Anguilla* species (*A. anguilla* and *A. rostrata*), with a particularly high proportion of hybrids in Iceland (Albert et al. 2006; Gagnaire et al., 2012; Wielgoss et al., 2014; Pujolar and Maes, 2016). However, while these Atlantic species have so far received most of the scientific attention, the greatest concentration of *Anguilla* species is present in the tropical Indo-Pacific, where 11 species occur and may partially spawn at the same locations (Kuroki et al., 2012; Arai, 2016). A locally high frequency of hybrids between one of the species pairs in this region, *A. marmorata* and *A. megastoma*, has been suggested by microsatellites and small datasets of species-diagnostic single-nucleotide polymorphisms (SNPs) (Schabetsberger et al. 2015); however, the pervasiveness of hybridization across all tropical eel species, the degree to which hybridization leads to introgression in these species, and the mechanisms maintaining species boundaries have so far remained poorly known.

In the present paper, we use high-throughput sequencing and morphological analyses for seven species of tropical eels sampled across the Indo-Pacific to (i) infer their age and diversification history, (ii) determine the frequencies of contemporary hybridization between the species, and (iii) quantify signatures of past introgression among them. Our unique combination of approaches allows us to compare hybridization and introgression across multiple pairs of animal species with different ages and demonstrates how cytonuclear incompatibilities and hybrid breakdown can strengthen species boundaries in the face of frequent hybridization.

## Results

### Extensive sampling

Collected in 13 field expeditions over the course of 14 years, our dataset included 456 individuals from 14 localities covering the distribution of anguillid eels in the tropical Indo-Pacific (Fig. 1a, Supplementary Table 1). Whenever possible, eels were tentatively identified morphologically in the field. Restriction-site-associated DNA (RAD) sequencing for all 456 individuals resulted in a comprehensive dataset of 704,480 RAD loci with a mean of 253.4 bp per locus and up to 1,518,299 SNPs, depending on quality-filtering options (Supplementary Figure 1). RAD sequences mapping to the mitochondrial genome unambiguously assigned all individuals to one of the seven tropical eel species *A. marmorata, A. megastoma, A. obscura, A. luzonensis, A. bicolor, A. interioris*, and *A. mossambica*, in agreement with our morphological assessment that indicated that the remaining four Indo-Pacific *Anguilla* species *A. celebesensis*, *A. bengalensis*, *A. borneensis*, and *A. reinhardtii* were not included in our dataset (Supplementary Figure 2). For those individuals for which sufficient morphological information was available (*n* = 161, restricted to *A. marmorata, A. megastoma, A. obscura*, and *A. interioris*), the measures predorsal length without head length (PDH) and distance between the origin of the dorsal fin and the anus (AD), size-standardized by total length (TL; Watanabe et al., 2009), revealed clear species-specific clusters but also intermediate individuals (Figure 1b, Supplementary Figure 3). This diagnosis was further supported by principal-component analysis (PCA) of seven morphological characters (Supplementary Figure 3). After excluding individuals with low-quality sequence data, the sample set used for genomic analyses contained 430 individuals of the seven species, including 325 *A. marmorata*, 41 *A. megastoma*, 36 *A. obscura*, 20 *A. luzonensis*, 4 *A. bicolor*, 3 *A. interioris*, and 1 *A. mossambica* (Supplementary Tables 2,3). The large number of individuals available for *A. marmorata, A. megastoma*, and *A. obscura*, sampled at multiple sites throughout their geographic distribution (Fig. 1a; Supplementary Table 1), permitted detailed analyses of genomic variation within these species (Supplementary Note 1). These analyses distinguished four populations in the geographically widespread species *A. marmorata* (Ishikawa et al., 2004; Minegishi et al., 2008; Watanabe et al., 2008; Gagnaire et al., 2011) but detected no population structure in *A. megastoma* and *A. obscura* (Supplementary Figure 4), that are both presumed to have a single spawning area in the western South Pacific (Schabetsberger et al., 2015, 2016).

**Figure 1:**
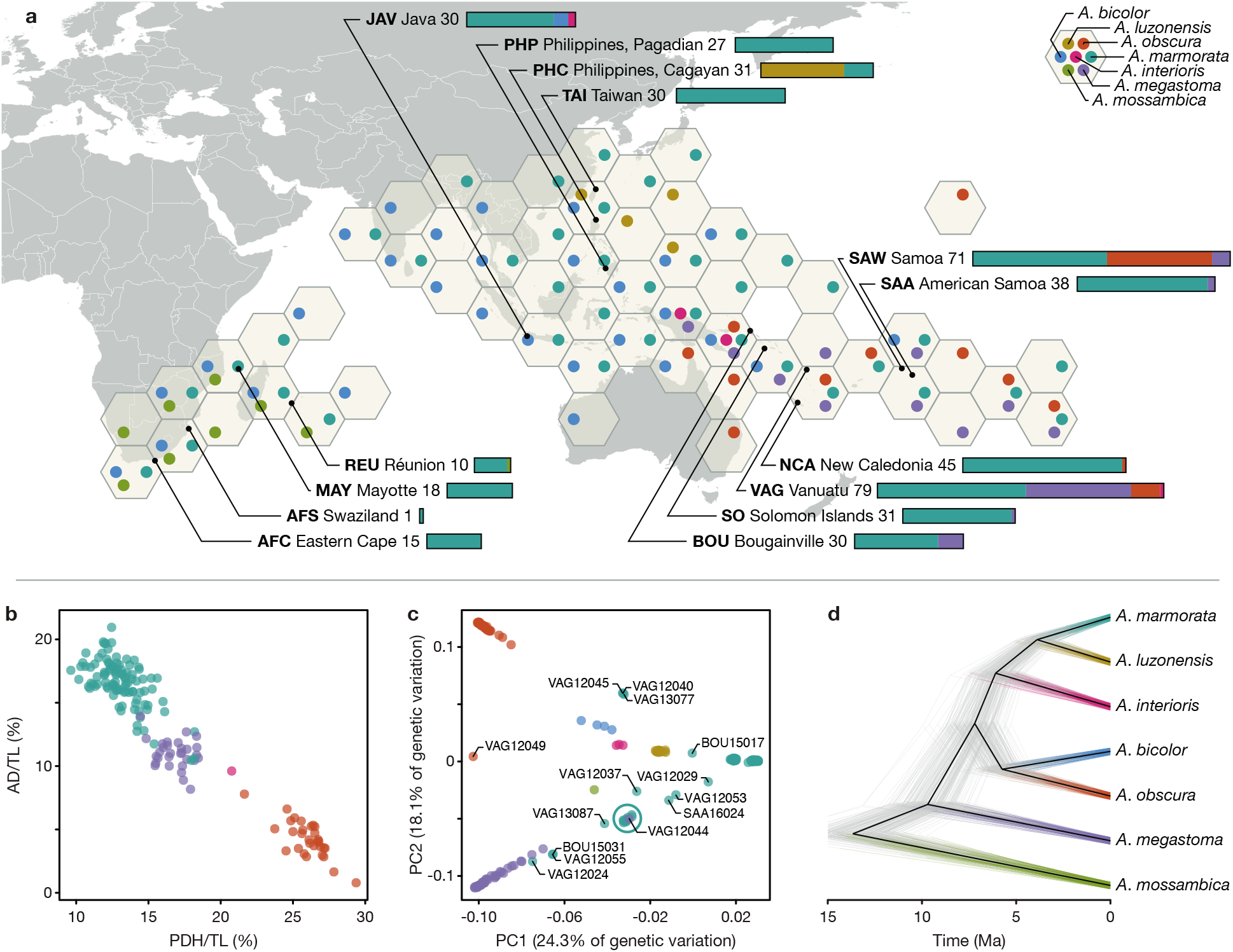
Genomic and morphological variation in tropical eels. **a)** Distribution of *Anguilla* species in the Indo-Pacific. The color and position of dots within hexagons indicate species presence within the region covered by the hexagon, according to the Global Biodiversity Information Facility database (GBIF.org 2019). Sampling locations are indicated with black dots. Numbers following location names specify the number of samples taken. Stacked bars indicate the species identities of individuals, according to mitochondrial and morphological species assignment. **b)** Morphological variation among the four species *A. marmorata* (*n* = 100), *A. megastoma* (*n* = 30), *A. obscura* (*n* = 30), and *A. interioris* (*n* = 1). Dots represent individuals and are colored according to mitochondrial species identity. **c)** Genomic PCA based on 155,896 variable sites. Specimen IDs are given for individuals with intermediate genotypes. The cyan circle indicates a cluster of 11 individuals mitochondrially assigned to *A. marmorata* (SAA16011, SAA16012, SAA16013, SAA16027, SAW17B27, SAW17B49, VAG12012, VAG12018, VAG12019, VAG13071, VAG13078), in addition to the highlighted VAG12044 which is mitochondrially assigned to *A. megastoma*. **d)** Time-calibrated phylogeny based on 5,000 transition sites. Each individual tree shown in gray represents a sample from the posterior tree distribution; a maximum-clade-credibility summary tree is shown in black. Color code in b), c), and d) is identical to a). PC: Principal component; AD: distance between the dorsal fin and the anus; PDH: predorsal length without head length; TL: total length.

### Deep divergences among tropical eel species

To analyze genomic variation among tropical eel species, we first performed PCA based on a dataset of 155,896 SNPs derived from RAD sequencing (Supplementary Figure 1). With few exceptions, the 430 individuals grouped according to species, and the seven species included in our dataset formed largely well-separated clusters (Fig. 1c, Supplementary Figure 5). Pairwise nuclear genetic distances between species ranged from 0.0053 to 0.0116 (uncorrected *p*-distance; excluding individuals with intermediate genotypes) and were largest for *A. mossambica* (0.0103-0.0116), followed by *A. megastoma* (0.0079-0.0090; excluding the comparison with *A. mossambica*, Supplementary Table 4). We further investigated the relationships among tropical eels species and their divergence times by applying Bayesian phylogenetic inference to genome-wide SNPs (Stange et al., 2018), using the multi-species coalescent model implemented in the software SNAPP (Bryant et al., 2012). As SNAPP does not account for rate variation among substitution types, we performed separate analyses with transitions and transversions, both of which supported the same species-tree topology. In agreement with the pairwise genetic distances, *A. mossambica* appeared as the sister to a clade formed by all other species, and *A. megastoma* was resolved within this clade as the sister to a group formed by the species pair *A. bicolor* and *A. obscura* and the species trio *A. marmorata, A. luzonensis*, and *A. interioris*, with *A. marmorata* and *A. luzonensis* being most closely related within this trio (Fig. 1d, Supplementary Figure 6). Each node of this species tree received full Bayesian support (Bayesian posterior probability, BPP, 1.0) regardless of whether transitions or transversions were used, and, except for the interrelationships of *A. marmorata, A. luzonensis*, and *A. interioris*, the tree agreed with previous phylogenies of mitochondrial sequences (Aoyama et al., 2001; Minegishi et al., 2005; Teng et al., 2009; Tseng, 2016, and references therein). Using a published age estimate for the divergence of *A. mossambica* (Jacobsen et al. 2014) to time calibrate the species tree, our analysis of transition SNPs with SNAPP showed that the clade combining all species except *A. mossambica* began to diverge around 9.7 Ma (divergence of *A. megastoma*; 95% HPD 11.7-7.7 Ma). This age estimate was robust to the use of transversions instead of transitions, alternative topologies enforced through constraints, and subsampling of taxa (Supplementary Figure 6).

To allow for the integration into other timelines of eel diversification based on multi-marker data (Rabosky et al., 2018; Musilova et al., 2019), we performed whole-genome sequencing (WGS) and generated new draft genome assemblies for *A. marmorata, A. megastoma*, and *A. obscura* (N50 between 54,849 bp and 64,770 bp; Supplementary Table 5), and extracted orthologs of the markers used in the studies of Musilova et al. (2019) and Rabosky et al. (2018). The use of these combined datasets together with age calibrations from the two studies also had little effect on age estimates, with the divergence of *A. megastoma* estimated around 6.5 Ma (95% HPD 7.2-5.8 Ma) or around 15.5 Ma (17.0-13.8 Ma), respectively (Supplementary Figure 6). Thus, all our analyses of divergence times point to an age of the clade formed by *A. marmorata, A. megastoma, A. obscura, A. luzonensis, A. bicolor*, and *A. interioris* roughly on the order of 10 Ma.

### High frequency of contemporary hybridization

Despite their divergence times up to around 10 Ma, our genomic dataset revealed ongoing hybridization in multiple pairs of tropical eel species. Analyses of genomic variation with PCA revealed a number of individuals with genotypes intermediate to the main clusters formed by the seven species (Fig. 1c, Supplementary Figure 5). The same individuals also appeared admixed in maximum-likelihood ancestry inference with the software ADMIXTURE (Alexander et al., 2009; Supplementary Figure 7) and had high levels of coancestry with two other species in analyses of RAD haplotype similarity with the program fineRADstructure, indicative of hybrid origin (Supplementary Figure 8; Malinsky et al., 2018b). For each of those putative hybrid individuals, we produced ancestry paintings (Runemark et al., 2018) based on sites that are fixed for different alleles in the parental species. In these ancestry paintings, the genotypes of the putative hybrids are assessed for those sites fixed between parents, with the expectation that first-generation (F1) hybrids should be heterozygous at almost all of these sites, and backcrossed hybrids of the second generation should be heterozygous at about half of them. All of the putative hybrids were confirmed by the ancestry paintings, showing that our dataset includes 20 hybrids between *A. marmorata* and *A. megastoma*, 3 hybrids between *A. marmorata* and *A. obscura*, 1 hybrid between *A. megastoma* and *A. obscura*, and 1 hybrid between *A. marmorata* and *A. interioris* (Fig. 2a-d, Supplementary Figures 9-12, Supplementary Tables 7). The frequency of hybrids in our dataset is thus 5.8% overall and up to 22.5% at the hybridization hotspot of Gaua, Vanuatu (Schabetsberger et al. 2015; Supplementary Figure 13, Supplementary Table 8). This high frequency is remarkable, given that most animal species produce hybrids at a frequency far below 1% (Mallet, 2005; Mallet et al., 2007). The heterozygosities of the hybrids are clearly bi-modal with a peak near 1 and another around 0.5 (Fig. 2i, Supplementary Table 7), supporting the presence of both first-generation hybrids as well as backcrossed second-generation hybrids. Using the mitochondrial genomes of hybrids as an indicator of their maternal species, we quantified the proportions of their nuclear genomes derived from the maternal species, *f*_m,genome_, based on their genotypes at the fixed sites used for ancestry painting. The distribution of these *f*_m,genome_ values has three peaks centered around 0.25 (4 individuals), 0.5 (18 individuals), and 0.75 (3 individuals), suggesting that backcrossing has occurred about equally often with both parental species (Fig. 2j). In agreement with the interpretation of seven individuals as backcrossed second-generation hybrids, scaffolds represented by multiple sites in the ancestry painting largely showed the same pattern at all of these sites, indicating that recombination breakpoints are rare (Supplementary Figure 14). However, chromosome-length assemblies would be required to exclude the presence of more than two recombination breakpoints on the same chromosome, which would indicate that hybridization occurred more than two generations ago.

In their size-standardized overall morphology, all hybrids for which morphological information was available (*n* = 15) were intermediate between the two parental species (Fig. 2e-h). Following Watanabe et al. (2009), we measured this overall morphology by the ratios AD/TL and PDH/TL, where AD is the distance between the dorsal fin and the anus, TL is the total length, and PDH is the predorsal length without the head. From these two ratios, we quantified the morphological similarity of hybrids to their maternal species relative to their paternal species, *f*_m,morphology_, as their position on an axis connecting the mean phenotypes of the two parental species. Similar to the distribution of *f*_m,genome_ values (Fig. 2j), the distribution of *f*_m,morphology_ values (Fig. 2k) also has three peaks centered close to 0.25, 0.5, and 0.75, even though these are less pronounced. In fact, the individuals with the lowest (BOU15031) and highest (VAG12029) *f*_m,morphology_ values also had the lowest and highest *f*_m,genome_ values, respectively (Supplementary Table 7), indicating that genomic similarity to parental species is correlated with morphological similarity (Fig. 2l).

**Figure 2:**
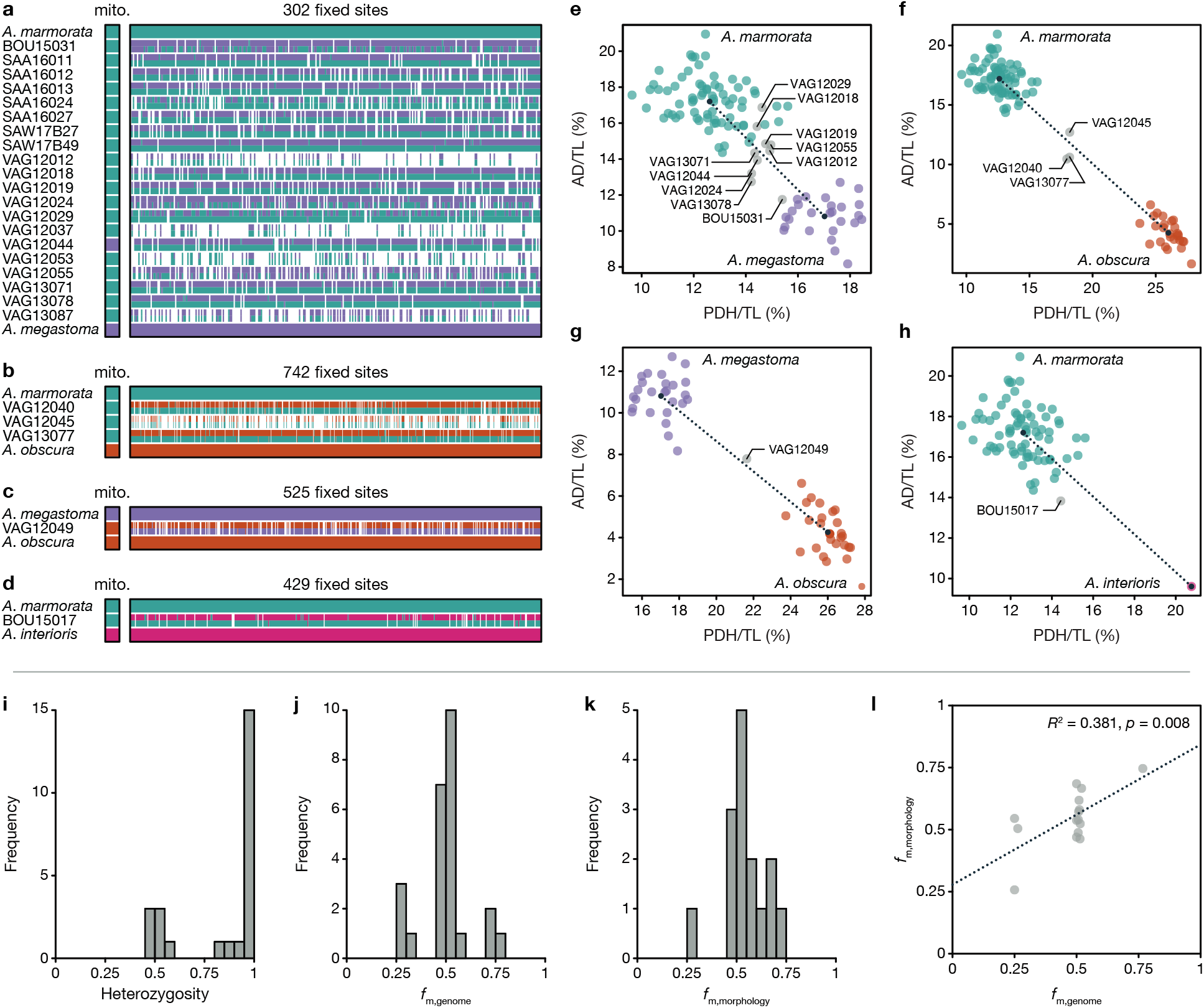
Contemporary hybridization among tropical eels. **a)** Ancestry painting for 20 hybrids between *A. marmorata* and *A. megastoma.* The top and bottom horizontal bars represent 302 sites that are fixed for different alleles between the two species; all other bars indicate the alleles at each of those sites. White color indicates missing data. Heterozygous alleles are shown with the top half in each bar matching the second parental species and vice versa. **b)** Ancestry painting for 3 contemporary hybrids between *A. marmorata* and *A. obscura*, based on 742 sites fixed between these two species. **c)** Ancestry painting for one hybrid between *A. megastoma* and *A. obscura*, based on 525 fixed sites. **d)** Ancestry painting for one hybrid between *A. marmorata* and *A. interioris*, based on 429 fixed sites. **e)** Morphological variation between *A. marmorata* and *A. megastoma*. Hybrids identified in **a)** are marked with specimen IDs. Mean phenotypes per species are marked with black dots that are connected by a dashed line **f-h)** as **e)** but for the hybrids identified in b)-d). **i)** Histogram of heterozygosity observed in hybrids. **j)** Histogram of the proportions of hybrid genomes derived from the maternal species (according to mitochondrial sequence data). **k)** Histogram of the relative morphological similarities between hybrids and the maternal species, measured as the relative proximity to the mean maternal phenotypes, compared to the proximity to the mean paternal phenotype. **l)** Comparison of the proportions of hybrids’ genomes derived from the maternal species and the similarity to the mean maternal species’ phenotype. The dotted line indicates a significant positive correlation between the two measures (*p* < 0.01; *R*^2^ = 0.381). mito: mitochondrial genome; AD: distance between the dorsal fin and the anus; PDH: predorsal length without head; TL: total length.

In contrast to their intermediate size-standardized overall morphology, hybrids in some cases had certain transgressive characters, exceeding the range of the parental phenotypes (Rieseberg et al., 1999; Supplementary Figures 14,15). This was the case for the total length and the length of the pectoral fin in VAG13071 and VAG12044, two F1 hybrids between *A. marmorata* and *A. megastoma* that were sampled in two successive years in Gaua, Vanuatu (Supplementary Table 1; Supplementary Figure 15). With a TL of 139 and 142 cm, the sizes of the two hybrids exceeded those of 229 other individuals (counting *A. marmorata*, *A. megastoma*, and their hybrids) for which this information was available by at least four centimeters (3%). Under a null hypothesis of no relation between hybridization and transgression, the probability that the largest two individuals are among the 18 hybrids for which TL was measured is *p* = 18/231 × 17/230 = 0.006; thus, it appears that transgression resulting from complementary gene action in hybrids (Stelkens and Seehausen, 2009) is responsible for their large sizes (considering only individuals from Gaua to account for possible location-size effects, this probability is *p* = 11/69 × 10/68 = 0.023). As we observed transgression only in hybrids between *A. marmorata* and *A. megastoma* (Supplementary Figure 15), but not in the hybrids of the more recently diverged species pair *A. marmorata* and *A. obscura* (Supplementary Figure 16), our results are consistent with the predicted increase of transgression with genetic distance among parental species (Stelkens et al., 2009; Stelkens and Seehausen, 2009; Arntzen et al., 2018).

### Evidence of past introgression

Multiple independent approaches revealed highly variable signatures of past introgression among species pairs of tropical eels. First, we found discordance between the Bayesian species trees based on the multi-species coalescent model (Fig. 1d) and an additional maximum-likelihood tree inferred with IQ-TREE (Nguyen et al., 2015; Supplementary Figure 17) from 1,360 concatenated RAD loci selected for high SNP density (Supplementary Figure 1). Even though both types of trees received full node support, their topologies differed in the position of *A. interioris*, which appeared next to *A. marmorata* and *A. luzonensis* in the Bayesian species trees (Fig. 1d, Supplementary Figure 6), but as the sister to *A. bicolor* and *A. obscura* in the maximum-likelihood tree, in agreement with mitochondrial phylogenies (Minegishi et al., 2005; Jacobsen et al., 2014). We applied an approach recently implemented in IQ-TREE (Minh et al., 2018) to assess per-locus and per-site concordance factors as additional measures of node support in the maximum-likelihood tree. These concordance factors were substantially lower than bootstrap-support values and showed that as few as 4.7% of the individual RAD loci and no more than 39.7% of all sites supported the position of *A. interioris* as the sister to *A. bicolor* and *A. obscura*.

To further test whether the tree discordance is due to past introgression or other forms of model misspecification, we applied genealogy interrogation (Arcila et al., 2017), comparing the likelihood of different topological hypotheses for each of the 1,360 RAD loci (Fig. 3a). We find that neither the topology of the Bayesian species trees nor the topology of the maximum-likelihood tree received most support from genealogy interrogation. Instead, 773 loci (62% of the informative loci) had a better likelihood when *A. interioris* was the sister to a clade formed by *A. marmorata*, *A. luzonensis*, *A. bicolor*, and *A. obscura*, compared to the topology of the Bayesian species tree (*A. interioris* as the sister to *A. marmorata* and *A. luzonensis*; Fig. 1d). The position of *A. interioris* as the sister to the other four species also had a better likelihood than the topology of the maximum-likelihood tree (*A. interioris* as the sister to *A. bicolor* and *A. obscura*; Supplementary Figure 17) for 659 loci (53% of the informative loci). We thus assumed that the topology supported by genealogy interrogation (with *A. interioris* being the sister to *A. marmorata*, *A. luzonensis*, *A. bicolor*, and *A. obscura*) is our best estimate of the true species-tree topology. However, we observed an imbalance in the numbers of loci supporting the two alternative topologies, as 541 loci had a better likelihood when *A. interioris* was the sister to *A. marmorata* and *A. luzonensis*, whereas 685 loci had a better likelihood when *A. interioris* was the sister to *A. bicolor* and *A. obscura* (Fig. 3a). As incomplete lineage sorting would be expected to produce equal support for both alternative topologies but the imbalance is too large to arise stochastically (exact binomial test *p* < 10^−4^), genealogy interrogation supports past introgression among *A. interioris*, *A. bicolor*, and *A. obscura*.

**Figure 3:**
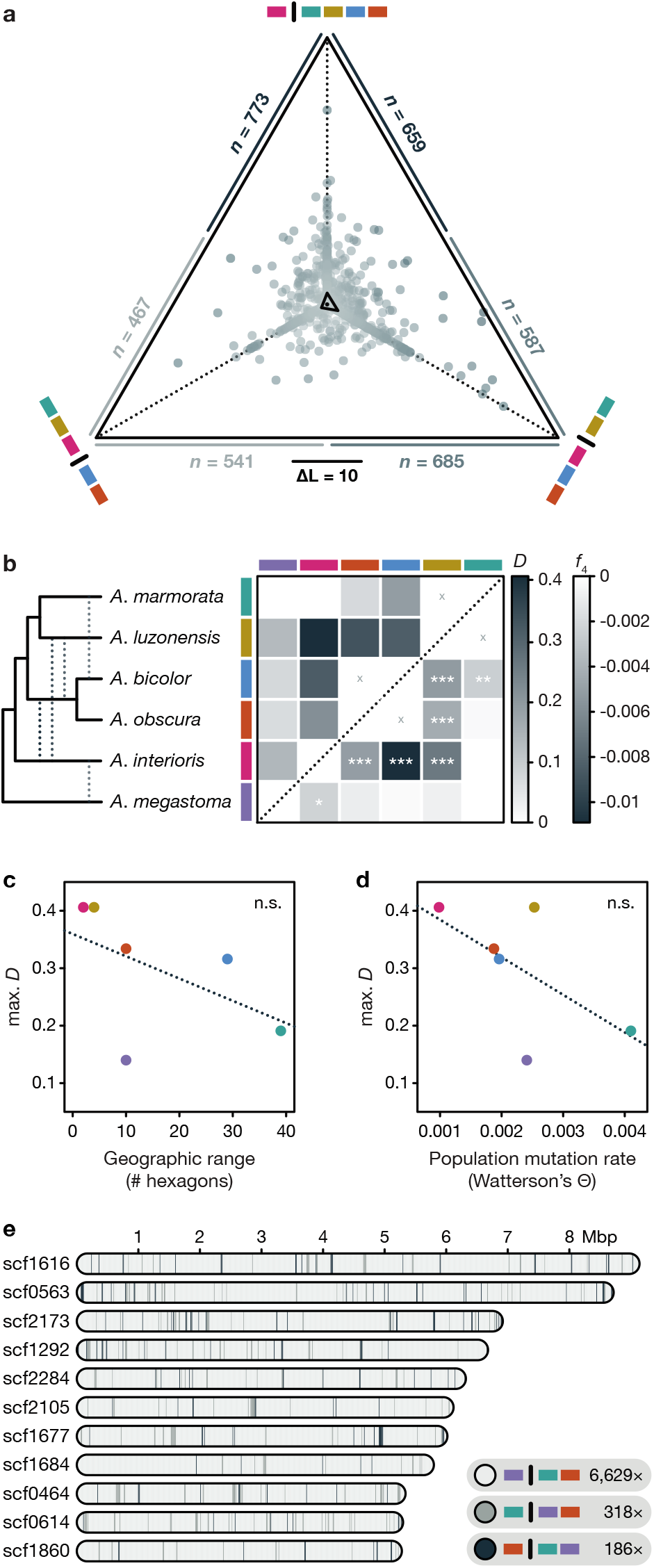
Past introgression among tropical eels. **a)** Likelihood support of individual RAD loci for different relationships of *A. interioris*: As sister to *A. marmorata* and *A. luzonensis* (bottom left), as sister to *A. obscura* and *A. bicolor* (bottom right), and as sister to a clade formed by those four species (top). The position of each dot shows the relative likelihood support of one RAD locus for each of the three tested relationships, with a distance corresponding to a log-likelihood difference of 10 indicated by the scale bar. The central triangle connects the mean relative likelihood support for each relationship. A black dot inside that triangle marks the central position corresponding to equal support for all three relationships. The two numbers outside each triangle edge report the number of loci that support each of the two competing relationships connected by that edge. **b)** Heatmap indicating maximum pairwise *D* (above diagonal) and *f*_4_ (below diagonal) statistics (see Table 1). Combinations marked with “x” symbols indicate sister taxa; introgression between these could not be assessed. Asterisks indicate the significance of *f*_4_ values (*: *p* < 0.05; **: *p* < 0.01; ***: *p* < 0.001), estimated with the F4 software (Meyer et al., 2017). The cladogram on the left summarizes the species-tree topology according to a) and the significant signals of introgression according to b). **c-d)** Comparisons of the maximum *D* value per species with the species’ geographic range or population mutation rate Θ. Geographic range was measured as the number of geographic hexagons (see Fig. 1) in which the species is present, and Watterson’s estimator (Watterson, 1975) was used for the population mutation rate Θ. n.s., not significant. **e)** Genomic patterns of phylogenetic relationships among *A. marmorata, A. obscura*, and *A. megastoma*, based on WGS reads mapped to the eleven largest scaffolds (those longer than 5 Mbp) of the *A. anguilla* reference genome. Blocks in light gray show 20,000-bp regions (incremented by 10,000 bp) in which *A. marmorata* and *A. obscura* appear as sister species, in agreement with the inferred species tree; in other blocks, *A. megastoma* appears closer to either *A. obscura* (gray) or *A. marmorata* (dark gray).

**Table 1:** Past introgression supported by *D* and *f*_4_ statistics. Only comparisons that are compatible with the inferred phylogenetic relationships and result in positive *D* values are shown (for all comparisons see Supplementary Table 9). All except the comparison in the last row are based on RAD-sequencing derived SNP data; the last comparison is based on WGS reads of a single individual of the three species. Either *A. mossambica, A. megastoma, A. interioris*, or *A. anguilla* (in the comparison based on WGS data) were used as outgroups and the comparison resulting in the largest *D* value is reported when multiple of these outgroups were used. *n*: number of sites variable among the included species; *C*_ABBA_: number of sites at which species P2 and P3 share the derived allele; *C*_BABA_: number of sites at which P1 and P3 share the derived allele.

| P1 | P2 | P3 | $n$ | $C_{ABBA}$ | $C_{BABA}$ | $D$ | $f_4$ | $p$ |
| --- | --- | --- | --- | --- | --- | --- | --- | --- |
| *A. marmorata* | *A. luzonensis* | *A. interioris* | 10,290 | 182.7 | 77.1 | 0.406 | -0.0070 | 0.000 |
| *A. marmorata* | *A. luzonensis* | *A. obscura* | 15,689 | 186.6 | 93.0 | 0.334 | -0.0043 | 0.000 |
| *A. marmorata* | *A. bicolor* | *A. interioris* | 7,772 | 266.3 | 138.4 | 0.316 | -0.0109 | 0.000 |
| *A. marmorata* | *A. luzonensis* | *A. bicolor* | 11,542 | 158.1 | 82.8 | 0.313 | -0.0052 | 0.000 |
| *A. marmorata* | *A. obscura* | *A. interioris* | 10,208 | 307.9 | 197.8 | 0.218 | -0.0051 | 0.000 |
| *A. obscura* | *A. bicolor* | *A. interioris* | 8,304 | 123.8 | 84.1 | 0.191 | -0.0030 | 0.005 |
| *A. obscura* | *A. bicolor* | *A. marmorata* | 11,372 | 104.7 | 71.2 | 0.191 | -0.0025 | 0.002 |
| *A. obscura* | *A. bicolor* | *A. luzonensis* | 12,557 | 113.4 | 80.0 | 0.173 | -0.0022 | 0.008 |
| *A. marmorata* | *A. interioris* | *A. megastoma* | 9,951 | 96.4 | 72.7 | 0.140 | -0.0023 | 0.026 |
| *A. marmorata* | *A. luzonensis* | *A. megastoma* | 13,129 | 69.0 | 52.9 | 0.133 | -0.0008 | 0.201 |
| *A. luzonensis* | *A. marmorata* | *A. bicolor* | 14,675 | 105.4 | 84.5 | 0.110 | -0.0011 | 0.106 |
| *A. luzonensis* | *A. bicolor* | *A. interioris* | 14,246 | 228.4 | 191.0 | 0.089 | -0.0015 | 0.062 |
| *A. luzonensis* | *A. interioris* | *A. megastoma* | 13,632 | 82.4 | 70.2 | 0.080 | -0.0007 | 0.192 |
| *A. marmorata* | *A. bicolor* | *A. megastoma* | 11,134 | 110.9 | 95.0 | 0.077 | -0.0003 | 0.430 |
| *A. luzonensis* | *A. marmorata* | *A. obscura* | 15,500 | 111.7 | 96.5 | 0.073 | -0.0003 | 0.406 |
| *A. marmorata* | *A. obscura* | *A. megastoma* | 11,647 | 126.1 | 110.0 | 0.068 | -0.0009 | 0.241 |
| *A. bicolor* | *A. obscura* | *A. marmorata* | 11,303 | 80.0 | 73.0 | 0.046 | -0.0007 | 0.261 |
| *A. obscura* | *A. bicolor* | *A. megastoma* | 11,761 | 64.7 | 59.5 | 0.042 | -0.0002 | 0.447 |
| *A. bicolor* | *A. obscura* | *A. luzonensis* | 15,856 | 78.1 | 72.1 | 0.040 | -0.0010 | 0.141 |
| *A. obscura* | *A. interioris* | *A. megastoma* | 11,017 | 96.2 | 90.8 | 0.029 | -0.0011 | 0.137 |
| *A. luzonensis* | *A. bicolor* | *A. megastoma* | 14,602 | 97.0 | 93.1 | 0.020 | 0.0002 | 0.416 |
| *A. bicolor* | *A. interioris* | *A. megastoma* | 10,451 | 84.0 | 82.0 | 0.012 | -0.0007 | 0.213 |
| *A. luzonensis* | *A. obscura* | *A. interioris* | 15,143 | 227.1 | 221.7 | 0.012 | 0.0005 | 0.300 |
| *A. luzonensis* | *A. obscura* | *A. megastoma* | 15,405 | 107.9 | 106.2 | 0.008 | -0.0001 | 0.461 |
| *A. obscura* | *A. marmorata* | *A. megastoma* | 23,165,451 | 596786.0 | 587910.0 | 0.007 | — | — |

We further quantified both Patterson’s *D* statistic (Green et al., 2010; Durand et al., 2011) and the *f*_4_ statistic (Reich et al., 2009) from biallelic SNPs, for all species quartets compatible with the species tree supported by genealogy interrogation. Both of these statistics are expected to be zero in the absence of introgression and thus support past introgression when they are found to differ from zero. As the distribution of these statistics is not usually normally-distributed across the genome (Meyer et al., 2017), we avoided block-jackknife resampling and instead assessed the significance of the *f*_4_ statistic with coalescent simulations in the software F4 (Meyer et al., 2017). We found that the *f*_4_ statistic was significant in no less than 29 out of 60 species quartets (Supplementary Table 9). The most extreme *D* and *f*_4_ values were observed in quartets in which *A. mossambica* was in the outgroup position, *A. marmorata* was in the position of the unadmixed species (P1), and *A. interioris* was in a position (P3) sharing gene flow with either *A. luzonensis* (*D* = 0.41) or *A. bicolor* (*f*_4_ = −0.011) (P2). The sum of the analyses of *D* and *f*_4_ suggests pervasive introgression among tropical eel species (Table 1), with significant support for gene flow between *A. interioris* and each of the three species *A. luzonensis*, *A. bicolor*, *A. obscura*, and *A. megastoma*, between *A. luzonensis* and both *A. bicolor* and *A. obscura*, and between *A. marmorata* and *A. bicolor* (Fig. 3b). While the pervasiveness of these signals prevents a clear resolution of introgression scenarios, the patterns could potentially be explained by a minimum of five introgression events: introgression between *A. megastoma* and *A. interioris*, between *A. interioris* and the common ancestor of *A. bicolor* and *A. obscura*, between *A. interioris* and *A. luzonensis*, between *A. luzonensis* and the common ancestor of *A. bicolor* and *A. obscura*, and between *A. bicolor* and *A. marmorata* (Fig. 3b). The four different populations of *A. marmorata* all showed nearly the same signal of gene flow with *A. bicolor*, indicating that the introgression between these species predates the origin of the observed spatial within-species differentiation in *A. marmorata* (Supplementary Table 10).

Interestingly, it appears that the species with the most restricted geographic distributions — *A. interioris* and *A. luzonensis* — are those with the strongest signals of past introgression (Fig. 3c), even though we identified only a single instance of contemporary hybridization involving one of these species (the first-generation hybrid BOU15017 with an *A. marmorata* mother and an *A. interioris* father; Fig. 2). In contrast, *A. marmorata* and *A. megastoma*, which both have a high population mutation rate Θ indicative of a large effective population size *N*_e_ (as Θ = 4*N_eμ_*), are those with the weakest signals of introgression (Fig. 3d) despite a high frequency of hybrids between them. While our sampling scheme does not allow us to exclude an effect of unequal sample sizes, this observation could be explained if introgressed alleles are over time more effectively purged by purifying selection from the genomes of species with larger effective population sizes (Harris and Nielsen, 2016; Juric et al., 2016). Particularly large effective population sizes in *A. marmorata* and *A. megastoma* are in fact supported by the WGS data produced for one individual of both species as well as *A. obscura.* When analyzed with the pairwise sequentially Markovian coalescent (PSMC; Li and Durbin, 2011), these data yielded estimates of a contemporary *N*_e_ between 9.9 × 10^4^ and 6.0 × 10^5^ for *A. marmorata* and between 2.3 × 10^5^ and 2.0 × 10^6^ for *A. megastoma*, whereas a comparatively lower *N*_e_ between 3.4 × 10^4^ and 7.4 × 10^4^ was estimated for the third species with WGS data, *A. obscura* (Supplementary Figure 18).

Low levels of introgression in the genomes of *A. marmorata* and *A. megastoma* were also supported by these WGS data. Aligning the WGS reads of *A. marmorata, A. megastoma*, and *A. obscura* to the *A. anguilla* reference-genome assembly (Jansen et al., 2017) resulted in an alignment with 23,165,451 genome-wide SNPs. Based on these SNPs, and using *A. anguilla* as the outgroup, the *D* value supporting gene flow between *A. marmorata* and *A. megastoma* was only 0.007 (Table 1). Phylogenetic analyses for 7,133 blocks of 20,000 bp, incremented by 10,000 bp, on the eleven largest scaffolds of the *A. anguilla* assembly showed that as many as 6,629 blocks (93%) support the species-tree topology, in which *A. marmorata* and *A. obscura* appear more closely related to each other than to *A. megastoma* (Fig. 3e). The alternative topologies with either *A. obscura* or *A. marmorata* being closer to *A. megastoma* were supported by 318 (4%) and 186 (3%) blocks, respectively. Notably, we did not observe long sets of adjacent blocks supporting the alternative topologies, which would be expected if the individuals had hybrids in their recent ancestry (Fu et al., 2015). The longest set of blocks supporting *A. marmorata* and *A. megastoma* as most closely related encompassed merely 80,000 bp (positions 4,890,000 to 4,970,000 on scaffold scf1677). While the lack of phasing information and a recombination map prevents a statistical test of time since admixture (Fu et al., 2015), the absence of longer sets of blocks most likely excludes hybrid ancestors within the last 10-20 generations.

## Discussion

As species diverge, genetic incompatibilities accumulate (Bateson, 1909; Dobzhansky, 1936; Muller, 1942) and reduce the viability of hybrids (Orr and Turelli, 2001). However, the absolute timescale on which hybrid inviability evolves vastly exceeds the ages of species in many diversifying clades, indicating that species boundaries in these groups are maintained by reproductive barriers that act after the F1 stage (Prager and Wilson, 1975; Coyne and Orr, 1989, 1997; Price and Bouvier, 2002; Bolnick and Near, 2008; Arntzen et al., 2009; Stelkens et al., 2010, 2015). For anguillid eels, laboratory experiments have produced hybrids between several species pairs, including *A. anguilla* and *A. australis* (Burgerhout et al., 2011), *A. anguilla* and *A. japonica* (Okamura et al., 2004; Müller et al., 2012), and *A. australis* and *A. dieffenbachii* (Lokman and Young, 2000). These species pairs diverged in some of the earliest divergence events within the genus (Supplementary Figure 6), suggesting that the limits of hybrid viability are not reached in anguillid eels. Our observation of frequent hybridization in four different species pairs, including two pairs involving *A. megastoma* with a divergence time around 10 Ma (Fig. 1d), supports this conclusion in a natural system, indicating that prezygotic reproductive barriers may generally be weak in tropical eels. This interpretation is strengthened by the fact that the 25 hybrids in our dataset were sampled in five different years (Supplementary Table 7), suggesting that natural hybridization in tropical eels occurs continuously, rather than, for example, being the result of an environmental change that ephemerally caused spatially and temporally overlapping spawning (Pujolar et al., 2014). Moreover, the seven identified backcrosses demonstrate that hybrids, at least those between *A. marmorata* and *A. megastoma*, can successfully reproduce naturally, indicating that, just like prezygotic barriers, postzygotic barriers are also incomplete in tropical eels, even after 10 million years of divergence.

Nevertheless, by considering both hybridization frequencies and introgression signals across multiple species pairs, our analyses reveal how tropical eel species have succeeded to prevent species collapse (and even diversify) despite their great potential for genomic homogenization. First, with a single exception, all of the 24 hybrids with *A. marmorata* as a parental species possessed the mitochondrial genome of this species, indicating that it is almost exclusively female *A. marmorata* that are involved in successful hybridization events. This asymmetry extends to later generations, because all seven backcrosses had the *A. marmorata* mitochondrial genome, and thus the mother’s mother must have been an *A. marmorata* for all backcrosses. Such asymmetry indicates differential viability of hybrids depending on the directionality of mating and could result from cytonuclear incompatibilities (Turelli and Moyle, 2007; Bolnick and Near, 2008; Arntzen et al., 2009; Gagnaire et al., 2012; Jacobsen et al., 2017). Second, the lower frequency of backcrosses compared to F1 hybrids and the lack of both F2 hybrids and later-generation backcrosses also suggest decreased fitness of hybrids. This hypothesis is supported by the observation that the *A. marmorata* and *A. megastoma* individuals selected for WGS apparently did not have recent hybrid ancestors, even though these individuals were sampled at the hybridization hotspot of Gaua, Vanuatu, where over 20% of all specimens are hybrids. Thus, it is possible that hybrid breakdown, affecting the viability and fertility of later-generation hybrids to a greater extent than F1 hybrids (Price and Bouvier, 2002; Wiley et al., 2009; Stelkens et al., 2015), is common in tropical eels and reduces the amount of introgression generated by backcrossing. Finally, the degree of introgression present in the genomes of tropical eel species appears to depend more on their population sizes than their hybridization frequencies, which could suggest that most introgressed alleles are purged from the recipient species by purifying selection (Harris and Nielsen, 2016; Juric et al., 2016). The combination of these mechanisms may thus effectively reduce gene flow among tropical eels to a trickle that is not strong enough to break down species boundaries. Over the last 10 million years, this trickle might nevertheless have contributed to the evolutionary success of anguillid eels by providing the potential for adaptive introgression (Abbott et al., 2013; Marques et al., 2019), whenever environmental changes required it. The identification of signatures of such introgression based on population-level whole-genome resequencing in tropical eels will be a promising goal for future studies.

## Methods

### Sample collection

A total of 456 *Anguilla* specimens were obtained from 14 main localities over 14 years (2003-2016, Fig. 1, Supplementary Table 1). Sampling localities included South Africa (AFC: *n* = 16), Swaziland (AFS: *n* = 1), Mayotte (MAY: *n* = 18), Réunion (REU: *n* = 10), Indonesia (JAV: *n* = 30), Philippines (PHC/PHP: *n* = 58), Taiwan (TAI: *n* = 30), Bougainville Island (BOU: *n* = 30), Solomon Islands (SOK/SOL/SON/SOR/SOV: *n* = 31), Vanuatu (VAG: *n* = 79), New Caledonia (NCA: *n* = 45), Samoa (SAW: *n* = 71), and American Samoa (SAA: *n* = 38). Sampling was performed as described by Schabetsberger et al. (2015) and Gubili et al. (2019), targeting elvers, yellow eels, and silver eels by electrofishing and with handnets in estuaries, rivers, and lakes. Small fin clips were extracted from the pectoral fin of each specimen and stored in 98% ethanol, to be used in subsequent genetic analyses. Permits were obtained prior to sampling from the responsible authorities.

### Morphological analyses

Morphological variation was assessed based on the following measurements: total length (TL), weight, preanal length (PA), predorsal length (PD), head length (HL), mouth length, eye distance, eye size (horizontal and vertical), pectoral fin size, head width, and girth (Watanabe et al., 2009). We further calculated the distance between the anus and the dorsal fin (AD = PA-PD), predorsal length without head length (PDH = PD-HL), tail length (T = TL-PA), and preanal length without head length (TR = PA-HL). Morphological variation was assessed with PCA in the program JMP v.7.0 (SAS Institute Inc.; www.jmp.com) based on the ratios of PA, T, HL, TR, PD, PDH, and AD to TL; this analysis was performed for 161 individuals for which all measurements were available (100 *A. marmorata*, 30 *A. megastoma*, 30 *A. obscura*, and 1 *A. interioris*). Principal-component scores were used to delimit “core” groups of putatively unadmixed individuals for the three species *A. marmorata* (73 individuals), *A. megastoma* (26 individuals), and *A. obscura* (26 individuals). In addition to PCA, we plotted the ratios of AD and PDH to TL, which were found to be particularly diagnostic for *Anguilla* species (Watanabe et al., 2004).

### Sequencing and quality filtering

Genomic DNA was extracted using the DNeasy Blood and Tissue Kit (Qiagen) as per the manufacturer’s instructions, or using a standard phenol chloroform procedure (Sambrook et al., 1989). DNA quality of each sample was evaluated on an agarose gel and quantified on a Qubit Fluorometer 2.0 (Thermo Fisher Scientific). Double-digest restriction-site associated DNA sequencing (ddRAD) was completed following Peterson et al. (2012) with minor modifications; this protocol is described in Supplementary Note 2.

Returned demultiplexed reads were processed using the software STACKS v.2.0-beta9 and v.2.2 (Catchen et al., 2013), following the protocol described by Rochette and Catchen (2017). In brief, the reads were checked for correct cut sites and adaptor sequences using the “process_radtags” tool and subsequently mapped against the European eel (*A. anguilla*) genome assembly (Jansen et al., 2017) using BWA MEM v.0.7.17 (Li and Durbin, 2009). As this assembly does not include the mitochondrial genome, mitochondrial reads were identified by separately mapping against the *A. japonica* mitochondrial genome (NCBI accession CM002536). Mapped reads were sorted and indexed using SAMTOOLS v.1.4 (Li, 2009, 2011). Species identification was verified for all individuals by comparing mitochondrial sequences with the NCBI Genbank database using BLAST v.2.7.1 (Altschul et al., 1990). Individuals with low-quality sequence data (with a number of reads below 600,000, a number of mapped reads below 70%, or a proportion of singletons above 5%) were excluded (*n* = 26). Variants were called using the “gstacks” tool, requiring a minimum mapping quality of 20 and an insert size below 500. Called variants were exported to variant call format (VCF) and haplotype format using the “populations” tool, allowing maximally 20% missing data and an observed heterozygosity below 75%, returning 1,518,299 SNPs.

The VCF file was further processed in two separate ways to generate suitable datasets for phylogenetic and population genetic analyses based on SNPs. For phylogenetic analyses, the VCF file was filtered with BCFTOOLS v.1.6 (Li, 2011) to mask genotypes if the per-sample read depth was below 5 or above 50 or if the genotype quality was below 30. Sites were excluded from the dataset if they appeared no longer polymorphic after the above modifications, if genotypes were missing for 130 or more of the 460 individuals (30%), or if their heterozygosity was above 50%. The resulting VCF file contained 619,353 SNPs (Supplementary Figure 1).

For analyses of genomic variation within and among species, filtering was done using VCFTOOLS v.0.1.14 (Danecek et al., 2011) and PLINK v.1.9 (Purcell et al., 2007). Sites were excluded if the mean read depth was above 50, the minor allele frequency was below 0.02, or heterozygosity excess was supported with *p* < 0.05 (rejecting the null hypothesis of no excess). In addition, individual genotypes were masked if they had a read depth below 5 or a genotype quality below 30. The resulting VCF file contained 155,896 SNPs (Supplementary Figure 1).

For each of the three species *A. marmorata*, *A. megastoma*, and *A. obscura*, one individual (VAG12030, VAG12032, and VAG12050, respectively) sampled in Gaua, Vanuatu, was subjected to WGS. Genomic DNA was extracted using the DNeasy Blood and Tissue Kit (Qiagen) according to the manufacturer’s protocol. DNA quality was evaluated on an agarose gel and quantified on a Qubit Fluorometer 2.0 (Thermo Fisher Scientific). All samples were sequenced on an Illumina HiSeq X Ten system at Macrogen (Korea) with the TruSeq DNA PCR-Free library kit (350 bp insert size) using 150 bp paired-end reads.

### Genome assembly

WGS reads for the three different species were error-corrected and trimmed for adapters with “merTrim” from the Celera Assembler software (Miller et al., 2008; downloaded from the CVS Concurrent Version System repository on 21 June 2017) using a k-mer size of 22 and the Illumina adapters option (Tørresen et al., 2017). Celera Assembler was run with the following options: merThreshold=0, merDistinct=0.9995, merTotal=0.995, unitigger=bogart, doOBT=0, doToggle=0; default settings were used for all other parameters. After assembly, the reads were mapped back to the assemblies using BWA MEM v.0.7.12, and consensus was recalled using Pilon v.1.22 (Walker et al., 2014). The completeness of the three different assemblies was assessed with BUSCO v.3.0.1 (Waterhouse et al., 2018) based on the vertebrate gene set.

### Analysis of mitochondrial haplotypes

RAD-sequencing reads mapping to the mitochondrial genome were converted to FASTA format using SAMTOOLS v.1.3, BCFTOOLS v.1.6, and SE-QTK v.1.0 (https://github.com/lh3/seqtk). Sequences corresponding to regions 10,630–10,720 and 12,015–12,105 of the *A. japonica* mitochondrial genome were aligned with default settings in MAFFT v.7.397 (Katoh and Standley, 2013) and the two resulting alignments were concatenated. The genealogy of mitochondrial haplotypes was reconstructed based on the GTRCAT substitution model in RAxML v.8.2.11 (Stamatakis, 2014) and used jointly with the concatenated alignment to produce a haplotype-genealogy graph with the software Fitchi v.1.1.4 (Matschiner, 2016).

### Species-tree inference

To estimate a time-calibrated species tree for the seven sampled *Anguilla* species, we applied the Bayesian molecular-clock approach of Stange et al. (2018) to a subset of the dataset of 619,353 SNPs, containing data for the maximally five individuals per species with the lowest proportions of missing data (28 individuals in total: 1 *A. mossambica*, 3 *A. interioris*, 4 *A. bicolor*, and 5 of each remaining species). By employing the SNAPP v.1.3 (Bryant et al., 2012) package for the program BEAST 2 v.2.5.0 (Bouckaert et al., 2019), the approach of Stange et al. (2018) integrates over all possible trees at each SNP and therefore allows accurate phylogenetic inference in the presence of incomplete lineage sorting. As the SNAPP model assumes a single rate of evolution for all substitution types, all SNAPP analyses were conducted separately for transitions and transversions. A maximum of 5,000 SNPs was used in both cases to reduce run times of the computationally demanding SNAPP analyses. After exploratory analyses unambiguously supported a position of *A. mossambica* outside of the other six sampled anguillid species, the root of the species tree was calibrated according to published estimates for the divergence time of *A. mossambica*. Specifically, we constrained this divergence to 13.76 Ma (with a standard deviation of 0.1 myr), as reported by Jacobsen et al. (2014) based on mitochondrial genomes of 15 anguillid species and three outgroup species. A justification of this timeline is given in Supplementary Note 3. Five replicate Markov-chain Monte Carlo (MCMC) analyses were conducted and convergence was confirmed with effective sample sizes (ESS) greater than 200, measured with the software Tracer v.1.7 (Rambaut et al., 2018). The posterior distributions of run replicates were merged after discarding the first 10% of each MCMC as burn-in, and maximum-clade-credibility (MCC) trees with node heights set to mean age estimates were generated with TreeAnnotator (Heled and Bouckaert, 2013). The robustness of divergence-time estimates was tested in a series of additional analyses, in which (i) alternative topologies were specified to fix the position of *A. interioris* (see below), (ii) species with strong signals of past introgression, *A. luzonensis* and *A. interioris* (see below), were excluded, (iii) genome assemblies of *A. marmorata, A. obscura*, and *A. megastoma* were used in combination with sequences and age constraints from Musilova et al. (2019), or (iv) mitochondrial sequences for the same three species were used jointly with sequences and age constraints from Rabosky et al. (2018). A full description of these additional analyses is presented in Supplementary Note 4.

The relationships among the seven sampled species *A. marmorata*, *A. luzonensis*, *A. bicolor*, *A. obscura, A. interioris, A. megastoma*, and *A. mossambica* were further investigated based on maximum likelihood, using the software IQ-TREE (Nguyen et al., 2015) and the same 28 individuals as in SNAPP analyses. RAD loci were filtered to exclude those with completely missing sequences and those with fewer than 20 (19,276 loci) or more than 40 variable sites (1 locus). The resulting dataset contained sequences from 1,360 loci with a total length of 393,708 bp and 0.18% of missing data (Supplementary Figure 1). The maximum-likelihood phylogeny was estimated from this set of loci with IQ-TREE’s edge-linked proportional-partition model that automatically selects the best-fitting substitution model for each locus. Node support was estimated with three separate measures: 1,000 ultrafast bootstrap-approximation replicates (Hoang et al., 2018) and gene- and site-specific concordance factors (Minh et al., 2018). These two types of concordance factors quantify the percentage of loci and sites, respectively, that support a given branch, and thus are a useful complement to bootstrap support values that are known to often overestimate confidence with phylogenomic data (Liu et al., 2015).

### Assessing genomic variation among and within species

Genome-wide variation was estimated based on the dataset of 155,896 SNPs, after excluding sites linked within 10-kb windows with *R*^2^ > 0.8 (Supplementary Figure 1). We performed PCA using smartPCA in EIGENSOFT v.6.0.1 (Patterson et al., 2006), including the function “lsqproject” to account for missing data, and through model-based clustering using ADMIXTURE v.1.3 (Alexander et al., 2009). Five replicates, each testing for one to eight clusters (*K*) and 10-fold cross-validation was performed.

The software fineRADstructure v.0.3.1 (Malinsky et al., 2018b) was used to infer genomic variation among individuals by clustering them according to similarity of their RAD haplotypes in a coancestry matrix. Haplotypes were exported using “populations” in Stacks (see above), additionally filtering for a minor allele frequency above 0.02 and a mean log likelihood greater than −10.0. The script “Stacks2fineRAD.py” (Malinsky et al., 2018b) was used to converte haplotypes of loci with maximally 20 variable sites to the fineRADstructure input format, resulting in a set of haplotypes for 65,912 RAD loci (Supplementary Figure 1). The coancestry matrix was inferred using RADpainter, and the MCMC clustering algorithm in fineSTRUCTURE v.4 (Lawson et al., 2012) was used to infer clusters of shared ancestry, setting the number of burnin iterations to 100,000, the sample iterations to 100,000, and the thinning interval to 1,000. Finally, to reflect the relationships within the co-ancestry matrix, the inferred clusters were arranged according to a tree inferred with fineSTRUCTURE, using 100,000 hill-climbing iterations and allowing all possible tree comparisons.

### Detecting contemporary hybridization

Based on the results of morphological and genomic PCA (Fig. 1, Supplementary Figures 3,5), analyses with ADMIXTURE (Supplementary Figure 7) and fineRADstructure (Supplementary Figure 8), and previous reports (Schabetsberger et al., 2015; Gubili et al., 2019), we suspected that our dataset included recent hybrids between four species pairs: *A. marmorata* and *A. megastoma, A. marmorata* and *A. obscura, A. megastoma* and *A. obscura*, and *A. marmorata* and *A. interioris.* To verify these putative hybrids, we determined sites that were fixed in each of the four species pairs, considering only the “core”-group individuals for *A. marmorata, A. megastoma*, and *A. obscura* (see section “Morphological analyses”; 73, 26, and 26 individuals, respectively) and the three available individuals for *A. interioris* (Supplementary Table 1). At each fixed site for which no more than 20% of genotypes were missing, we then assessed the genotypes of the putative hybrids and plotted these in the form of “ancestry paintings” (Runemark et al., 2018). We expected that first-generation (F1) hybrids would be consistently heterozygous at nearly all sites fixed for different alleles between parental species (some few loci that appear fixed between the sampled individuals of the parental species might not be entirely fixed in those species), and that backcrossed individuals would show a heterozygosity (*h*_fixed_) of around 50% or less at these sites. For each verified F1 or backcrossed hybrid, we further quantified the proportion of its genome derived from the maternal species, *f*_m,genome_, based on its genotypes at the sites fixed between parents and assuming that its mitochondrial genome reliably indicates the species of its mother. Finally, we also quantified the relative morphological similarity to the maternal species, *f*_m,morphology_, for each hybrid, corresponding to the position of the hybrid on an axis connecting the mean morphology of the maternal species with the mean morphology of the paternal species.

Specifically we calculated this relative similarity as

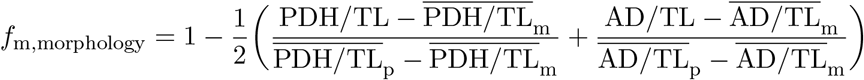

where 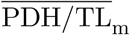 is the mean PDH divided by TL of the maternal species, 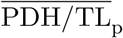 is the mean PDH divided by TL of the paternal species, 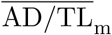 is the mean AD divided by TL of the maternal species, and 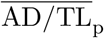 is the mean AD divided by TL of the paternal species.

### Detecting past introgression

As our analyses of contemporary hybridization identified several backcrossed individuals, we assumed that, despite their old divergence times, tropical eel species may have remained connected by continuous or episodic gene flow. We thus tested for signals of past introgression among the seven species using multiple complementary approaches. Our first approach was motivated by the observation that *A. interioris* clustered with *A. marmorata* and *A. luzonensis* in the Bayesian species-tree analyses with SNAPP, but appeared as the sister to *A. bicolor* and *A. obscura* in the maximum-likelihood phylogeny generated with IQ-TREE, with strong support in both cases. Assuming that this discordance might have resulted from past introgression (e.g., Martin et al., 2019), we thus applied genealogy interrogation (Arcila et al., 2017) to the dataset used for IQ-TREE analyses, composed of 1,360 RAD loci with a total length of 393,708 bp. For each of these loci, we separately calculated the likelihood of three different topological hypotheses (H1-H3): *A. interioris* forming a monophyletic group with *A. marmorata* and *A. luzonensis* to the exclusion of *A. bicolor* and *A. obscura* (H1), *A. interioris* forming a monophyletic group with *A. bicolor* and *A. obscura* to the exclusion of *A. marmorata* and *A. luzonensis* (H2), or *A. marmorata, A. luzonensis, A. bicolor*, and *A. obscura* forming a monophyletic group to the exclusion of *A. interioris* (H3). These likelihood calculations were performed using IQ-TREE with the GTR substitution model, and two replicate analyses were conducted for each combination of locus and hypothesis. Per locus, we then compared the three resulting likelihoods and quantified the numbers of loci supporting H1 over H2, H2 over H1, H1 over H3, H3 over H1, H2 over H3, and H3 over H2. We expected that the true species-tree topology would be supported by the largest number of loci, and that introgression would, if present, increase the support for one of the alternative hypotheses relative to the other (Schumer et al., 2016, Meyer et al. 2017).

As a second approach for the detection of past introgression, we calculated Patterson’s *D* statistic (Green et al., 2010; Durant et al., 2011) from biallelic SNPs included in the RAD-sequencing derived dataset of 619,353 SNPs (Supplementary Table 1). As this statistic is applicable to quartets of species in which one is the outgroup to all others and two species (labeled P1 and P2) are sister taxa, we calculated the *D* statistic separately for all species quartets compatible with the species tree inferred through genealogy interrogation. In this species tree, *A. mossambica* is the sister to all other species and *A. interioris* is the sister to a clade formed by the two species pairs *A. marmorata* and *A. luzonensis* and *A. bicolor* and *A. obscura.* Per species quartet, the *D* statistic was calculated as

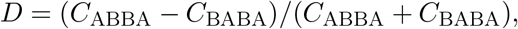

where *C*_ABBA_ is the number of sites at which P2 and the third species (P3) share a derived allele and *C_BABA_* is the number of sites at which P1 and P3 share the derived allele. If sites were not fixed within species, allele frequencies were taken into account following Martin et al. (2015). In the absence of introgression, *D* is expected to be zero; positive *D* values are expected when introgression took place between P2 and P3, and negative *D* values result from introgression between P1 and P3.

In addition to the above analyses based on RAD-sequencing derived SNPs, the WGS data for *A. marmorata, A. megastoma*, and *A. obscura*, in combination with the available reference-genome assembly for *A. anguilla* (Jansen et al., 2017), allowed us to calculate *D* statistics for this species quartet from a fully genomic dataset. To this end, WGS reads of the three species were mapped against the *A. anguilla* reference assembly using BWA MEM, and sorted and indexed using SAMTOOLS. Duplicates were marked using PICARD-TOOLS v.2.6.0 (http://broadinstitute.github.io/picard/), and indels were realigned using GATK v.3.4.64 (McKenna et al., 2010). Per-species mean read coverage (71.31×, 64.80×, and 48.97× for *A. marmorata, A. megastoma*, and *A. obscura*, respectively) was calculated with BEDTOOLS v.2.26.0 (Quinlan and Hall, 2010). SNP calling was performed using SAMTOOLS’ “mpileup” command, requiring a minimum mapping quality (MQ) of 30 and a base quality (BQ) greater than 30, before extracting the consensus sequence using BCFTOOLS v.1.6. The consensus sequences were converted to FASTQ format via SAMTOOLS’ “vcfutils” script for bases with a read depth (DP) between 15 and 140, and subsequently used to calculate the genome-wide *D* statistic with *A. obscura* as P1, *A. marmorata* as P2, *A. megastoma* as P3, and *A. anguilla* as the outgroup.

The dataset of 619,353 RAD-sequencing derived SNPs (Supplementary Table 1) was further used to calculated the *f*_4_ statistic (Reich et al., 2009) as a separate measure of introgression signals, for the same species quartets as the *D* statistic. The *f*_4_ statistic is based on allele-frequency differences between the species pair formed by P1 and P2 and the species pair formed by P3 and the outgroup (as the *f*_4_ statistic does not assume a rooted topology, P3 and the outgroup form a pair when P1 and P2 are monophyletic), and like the *D* statistic, the *f*_4_ statistic is expected to be zero in the absence of introgression. We calculated the *f*_4_ statistic with the F4 program v.0.92 (Meyer et al., 2017). As the distribution of the *f*_4_ statistic across the genome is usually not normally distributed, block-jackknife resampling is not an appropriate method to assess its significance; thus, we estimated *p*-values based on coalescent simulations as described in Meyer et al. (2017). In brief, these simulations are also conducted with the F4 program, internally employing fastsimcoal v.2.5.2 (Excoffier et al., 2013) to run each individual simulation. After a burnin period required to adjust settings for divergence times and population sizes in the simulations, the set of simulations allows the estimation of the *p*-value for the hypothesis of no introgression as the proportion of simulations that resulted in an *f*_4_ statistic as extreme or more extreme than the *f*_4_ statistic of the empirical species quartet.

The genome-wide consensus sequences for *A. marmorata, A. megastoma*, and *A. obscura*, aligned to the *A. anguilla* reference-genome assembly (Jansen et al., 2017), were further used to test for introgressed regions on the largest scaffolds of the reference genome (11 scaffolds with lengths greater than 5 Mbp). To this end, maximum-likelihood phylogenies of the four species were generated with IQ-TREE for blocks of 20,000 bp, incremented by 10,000 bp, with IQ-TREE settings as described above for species-tree inference.

### Estimating effective population sizes

Distributions of genome-wide coalescence times were inferred from WGS reads of *A. marmorata, A. megastoma*, and *A. obscura* using the pairwise sequentially Markovian coalescent (PSMC; Li and Durbin, 2011). Heterozygous sites were detected from consensus sequences in FASTQ format (see above) using the script “fq2psmcfa” (Li and Durbin, 2011), applying a window size of 20 bp (1.4% of windows contained more then one heterozygous site), and a scaffold-good-size of 10,000 bp. The PSMC analyses were run for 30 iterations, setting the initial effective population size to 15, the initial Θ to five, and the time-intervals option to “4 × 4 + 13 × 2 + 4 × 4 + 6”, corresponding to 22 free parameters. To assess confidence intervals, 100 bootstrap replicates were performed using the script “splitfa” (Li and Durbin, 2011). The PSMC plots were scaled using generation times reported by Jacoby et al. (2015); these were 12 years, 10 years, and 6 years for *A. marmorata, A. megastoma*, and *A. obscura*, respectively. Mutation rates were calculated based on pairwise genetic distances and divergence-time estimates inferred in our phylogenetic analyses. Uncorrected *p*-distances were 1.199% between *A. marmorata* and *A. megastoma*, 1.307% between *A. megastoma* and *A. obscura*, and 1.141% between *A. marmorata* and *A. obscura.* In combination with the divergence time of *A. megastoma* at 9.6954 Ma and the divergence time between *A. marmorata* and *A. obscura* at 7.2023 Ma, these distances resulted in mutation-rate estimates per site per generation of *r* = 8.6 × 10^−9^, 5.6 × 10^−9^, and 5.2 × 10^−9^ for *A. marmorata*, *A. megastoma*, and *A. obscura*, respectively.

## Data availability

The raw RADseq data will be deposited on the NCBI SRA database. Genome assemblies for *A. marmorata, A. megastoma*, and *A. obscura* are deposited on ENA with project number PRJEB32187. Morphological measurements, SNP datasets in VCF format, and input and output of phylogenetic analyses will be deposited on Dryad. Code for computational analyses is available from Github (http://github.com/mmatschiner/anguilla).

## Acknowledgements

Funding for this study was provided by the Austrian Science Fund (FWF, project P28381-B29 to R.Sc.) and the Norwegian Research Council (FRIPRO project 275869 to M.M.). We thank Anthony Acou, Davi Boseto, Donna Kalfatak, Rilloy Leaana, Finn Økland, Christine Pöllabauer, Alexander Scheck, Ursula Sichrowsky, and Meelis Tambets for assistance with field work, Franz Gassner for help with data analysis, and Ian Goodhead for his support in the laboratory. We further thank Olaf L. F. Weyl and acknowledge the DST/NRF South African Research Chair in Inland Fisheries and Freshwater Ecology (Grant No 110507) and the NRF-SAIAB Collections Platform for the provision of genetic tissue samples. All computational work was performed on the Abel Supercomputing Cluster (Norwegian Metacenter for High-Performance Computing (NOTUR) and the University of Oslo), operated by the Research Computing Services group at USIT, the University of Oslo IT Department. Peter Comes provided valuable comments on the manuscript.

## Author contributions

R.Sc. and R.J. conceived the project. R.Sc., R.J., C.G., M.M., J.M.I.B., and R.So. planned and oversaw the project. R.Sc., C.G., Y.-S.H., and E.F. contributed specimens. C.G., R.J., and R.Sc. organized RAD sequencing. S.W. performed morphological analyses. J.M.I.B. and B.E. prepared genomic datasets. O.K.T. performed genome assembly. M.M. and J.M.I.B. performed population genomic and phylogenomic analyses. M.M. and J.M.I.B. prepared the figures. M.M. and J.M.I.B. wrote the manuscript with input from all authors.

## References

Abbott, R. et al. Hybridization and speciation. J. Evol. Biol. 26, 229–246 (2013).

Albert, V. Jónsson, B. & Bernatchez, L. Natural hybrids in Atlantic eels (*Anguilla anguilla, A. rostrata*): evidence for successful reproduction and fluctuating abundance in space and time. Mol. Ecol. 15, 1903–1916 (2006).

Alexander, D. H. Novembre, J. & Lange, K. Fast model-based estimation of ancestry in unrelated individuals. Genome Res. 19, 1655–1664 (2009).

Altschul, S. F. Gish, W. Miller, W. Myers, E. W. & Lipman, D. J. Basic local alignment search tool. J. Mol. Biol. 215, 403–410 (1990).

Aoyama, J. Nishida, M. & Tsukamoto, K. Molecular phylogeny and evolution of the freshwater eel, genus *Anguilla*. Mol. Phyl. Evol. 20, 450–459 (2001).

Arai, T. Biology and Ecology of Anguillid Eels (CRC Press, Boca Raton, Florida, 2016).

Arcila, D. et al. Genome-wide interrogation advances resolution of recalcitrant groups in the tree of life. Nat. Ecol. Evol. 1, 1–10 (2017).

Arntzen, J. W. Jehle, R. Bardakci, F. Burke, T. & Wallis G. P. Asymmetric viability of reciprocal-cross hybrids between crested and marbled newts (*Triturus cristatus* and *T. marmoratus*). Evolution 63, 1191–1202 (2009).

Arntzen, J. W. Üzüm, N. Ajduković, M. D. Ivanović, A. & Wielstra, B. Absence of heterosis in hybrid crested newts. PeerJ 6, e5317 (2018).

Avise, J. C. et al. The evolutionary genetic status of Icelandic eels. Evolution 44, 1254–1262 (1990).

Bateson, W. in Darwin and Modern Science (ed Seward, A. C.) 85–101 (Cambridge University Press, Cambridge, 1909).

Bolnick, D. I. & Near, T. J. Tempo of hybrid inviability in centrarchid fishes (Teleostei: Centrarchidae). Evolution 59, 1754–1767 (2008).

Bouckaert, R. R. et al. BEAST 2.5: An advanced software platform for Bayesian evolutionary analysis. PLoS Comput. Biol. 15, e1006650 (2019).

Bryant, D. Bouckaert, R. Felsenstein, J. Rosenberg, N. A. & Choudhury, A. R. Inferring species trees directly from biallelic genetic markers: Bypassing gene trees in a full coalescent analysis. Mol. Biol. Evol. 29, 1917–1932 (2012).

Burgerhout, E. et al. First artificial hybrid of the eel species *Anguilla australis* and *Anguilla anguilla*. BMC Dev. Biol. 11, 16 (2011).

Catchen, J. Hohenlohe, P. A. Bassham, S. Amores, A. & Cresko W. A. Stacks: an analysis tool set for population genomics. Mol. Ecol. 22, 3124–3140 (2013).

Coyne, J. A. & Orr, H. A. Patterns of speciation in *Drosophila*. Evolution 43, 362–381 (1989).

Coyne, J. A. & Orr H. A. “Patterns of speciation in *Drosophila*” revisited. Evolution 51, 295–303 (1997).

Danecek, P. et al. The variant call format and VCFtools. Bioinformatics 27, 2156–2158 (2011).

Dobzhansky, T. Studies on hybrid sterility. II. Localization of sterility factors in *Drosophila pseudoobscura* hybrids. Genetics 21, 113–135 (1936).

Durand, E. Y. Patterson, N. Reich, D. & Slatkin, M. Testing for ancient admixture between closely related populations. Mol. Biol. Evol. 28, 2239–2252 (2011).

Edelman, N. B. et al. Genomic architecture and introgression shape a butterfly radiation. Preprint at https://www.biorxiv.org/content/10.1101/466292v1 (2018).

Excoffier, L. Dupanloup, I. Huerta-Sánchez, E. Soussa, V. & Foll, M. Robust demographic inference from genomic and SNP data. PLoS Genet. 9, e1003905 (2013).

Fu, Q. et al. An early modern human from Romania with a recent Neanderthal ancestor. Nature 524, 216–219 (2015).

Gagnaire, P.-A. et al. Within population structure highlighted by differential introgression across semipermeable barriers to gene flow in *Anguilla marmorata*. Evolution 65, 3413–3427 (2011).

Gagnaire, P.-A. Normandeau, E. & Bernatchez, L. Comparative genomics reveals adaptive protein evolution and a possible cytonuclear incompatibility between European and American eels. Mol. Biol. Evol. 29, 2909–2919 (2012).

GBIF.org GBIF Home Page. Available from: https://www.gbif.org (2019).

Green, R. E. et al. A draft sequence of the Neandertal genome. Science 328, 710–722 (2010).

Gubili, C. et al. High genetic diversity and lack of pronounced population structure in five species of sympatric Pacific eels. Fish. Manag. Ecol. 26, 31–41 (2019).

Harris, K. & Nielsen, R. The genetic cost of Neanderthal introgression. Genetics 203, 881–891 (2016).

Heled, J. & Bouckaert, R. R. Looking for trees in the forest: summary tree from posterior samples. BMC Evol. Biol. 13, 221 (2013).

Hench, K. Vargas, M. Höppner, M. P. McMillan, W. O. Puebla, O. Inter-chromosomal coupling between vision and pigmentation genes during genomic divergence. Nat. Ecol. Evol. 3, 657–667 (2019).

Hoang, D. T. Vinh, L. S. Flouri, T. Stamatakis, A. & von Haeseler, A. MPBoot: fast phylogenetic maximum parsimony tree inference and bootstrap approximation. BMC Evol. Biol. 18, 11 (2018).

Ishikawa, S. Tsukamoto, K. & Nishida, M. Genetic evidence for multiple geographic populations of the giant mottled eel *Anguilla marmorata* in the Pacific and Indian oceans. Ichthyol. Res. 51, 343–353 (2004).

Jacobsen, M. W. et al. Speciation and demographic history of Atlantic eels (*Anguilla anguilla* and *A. rostrata*) revealed by mitogenome sequencing. Heredity 113, 432–442 (2014).

Jacobsen, M. W. et al. Assessing pre- and post-zygotic barriers between North Atlantic eels (*Anguilla anguilla* and *A. rostrata*). Heredity 118, 266–275 (2017).

Jacoby, D. M. P. et al. Synergistic patterns of threat and the challenges facing global anguillid eel conservation. Glob. Ecol. Conserv. 4, 321–333 (2015).

Jansen, H. J. et al. Rapid de novo assembly of the European eel genome from nanopore sequencing reads. Sci. Rep. 7, 7213 (2017).

Juric, I. Aeschbacher, S. & Coop, G. The strength of selection against Neanderthal introgression. PLoS Genet. 12, e1006340 (2016).

Katoh, K. & Standley, D. M. MAFFT multiple sequence alignment software version 7: Improvements in performance and usability. Mol. Biol. Evol. 30, 772–780 (2013).

Kozak, K. M. McMillan, W. O. Joron, M. & Jiggins, C. D. Genome-wide admixture is common across the *Heliconius* radiation. Preprint at https://www.biorxiv.org/content/10.1101/414201v1 (2018).

Kuroki, M. et al. Offshore spawning for the newly discovered anguillid species *Anguilla luzonensis* (Teleostei: Anguillidae) in the Western North Pacific. Pac. Sc. 66, 497–507 (2012).

Lamichhaney, S. et al. Rapid hybrid speciation in Darwin’s finches. Science 359, 224–228 (2018).

Lawson, D. J. Hellenthal, G. Myers, S. & Falush, D. Inference of population structure using dense haplotype data. PLoS Genet. 8, e1002453 (2012).

Li, H. The Sequence Alignment/Map format and SAMtools. Bioinformatics 25, 2078–2079 (2009).

Li, H. A statistical framework for SNP calling, mutation discovery, association mapping and population genetical parameter estimation from sequencing data. Bioinformatics 27, 2987–2993 (2011).

Li, H. & Durbin, R. Fast and accurate short read alignment with Burrows-Wheeler transform. Bioinformatics 25, 1754–1760 (2009).

Li, H. & Durbin, R. Inference of human population history from individual whole-genome sequences. Nature 475, 493–496 (2011).

Liu, L. Xi, Z. Wu, S. Davis, C. C. Edwards, S. V. Estimating phylogenetic trees from genome-scale data. Ann. New York Acad. Sci. 1360, 36–53 (2015).

Lokman, P. M. & Young, G. Induced spawning and early ontogeny of New Zealand freshwater eels. New Zealand J. Mar. Freshw. Res. 34, 135–145 (2000).

Malinsky, M. et al. Whole-genome sequences of Malawi cichlids reveal multiple radiations interconnected by gene flow. Nat. Ecol. Evol. 2, 1940–1955 (2018a).

Malinsky, M. Trucchi, E. Lawson, D. J. & Falush, D. RADpainter and fineRADstructure: Population inference from RADseq data. Mol. Biol. Evol. 35, 1284–1290 (2018b).

Mallet, J. Hybridisation as an invasion of the genome. Trends Ecol. Evol. 20, 229–237 (2005).

Mallet, J. Hybrid speciation. Nature 446, 279–283 (2007).

Mallet, J. Beltrán, M. Neukirchen, W. & Linares, M. Natural hybridization in heliconiine butterflies: the species boundary as a continuum. BMC Evol. Biol. 7, 28 (2007).

Marques, D. A. Meier, J. I. & Seehausen, O. A combinatorial view on speciation and adaptive radiation. Trends Ecol. Evol. Advance access at https://www.cell.com/trends/ecology-evolution/fulltext/S0169-5347(19)30055-2 (2019).

Martin, S. H. Davey, J. W. & Jiggins, C. D. Evaluating the use of ABBA-BABA statistics to locate introgressed loci. Mol. Biol. Evol. 32, 244–257 (2015).

Martin, S. H. Davey, J. W. Salazar, C. & Jiggins, C. D. Recombination rate variation shapes barriers to introgression across butterfly genomes. PLoS Biol. 17, e2006288 (2019).

Matschiner, M. Fitchi: haplotype genealogy graphs based on the Fitch algorithm. Bioinformatics 32, 1250–1252 (2016).

Matschiner, M. et al. Bayesian phylogenetic estimation of clade ages supports trans-Atlantic dispersal of cichlid fishes. Syst. Biol. 66, 3–22 (2017).

McKenna, A. et al. The Genome Analysis Toolkit: a MapReduce framework for analyzing next-generation DNA sequencing data. Genome Res. 20, 1297–1303 (2010).

Meier, J. I. et al. Ancient hybridization fuels rapid cichlid fish adaptive radiations. Nat. Commun. 8, 14363 (2017).

Meyer, B. S. Matschiner, M. & Salzburger, W. Disentangling incomplete lineage sorting and introgression to refine species-tree estimates for Lake Tanganyika cichlid fishes. Syst. Biol. 66, 531–550 (2017).

Miller, J. R. et al. Aggressive assembly of pyrosequencing reads with mates. Bioinformatics 24, 2818–2824 (2008).

Minegishi, Y. et al. Molecular phylogeny and evolution of the freshwater eels genus *Anguilla* based on the whole mitochondrial genome sequences. Mol. Phyl. Evol. 34, 134–146 (2005).

Minegishi, Y. Aoyama, J. & Tsukamoto, K. Multiple population structure of the giant mottled eel, *Anguilla marmorata*. Mol. Ecol. 17, 3109–3122 (2008).

Minh, B. Q. Hahn, M. & Lanfear, R. New methods to calculate concordance factors for phylogenomic datasets. Preprint at https://www.biorxiv.org/content/10.1101/487801v1 (2018).

Muller, H. J. Isolating mechanisms, evolution and temperature. Biol. Symp. 6, 71–125 (1942).

Müller, T. et al. Artificial hybridization of Japanese and European eel (*Anguilla japonica × A. anguilla*) by using cryopreserved sperm from freshwater reared males. Aquaculture 350–353, 130–133 (2012).

Musilova, Z. et al. Vision using multiple distinct rod opsins in deep-sea fishes. Science 364, 588–592 (2019).

Nelson, J. S. Grande, T. C. & Wilson, M. V. H. Fishes of the World, 5th ed. (John Wiley & Sons, Inc. Hoboken, New Jersey, 2018).

Nguyen, L. T. Schmidt, H. A. von Haeseler, A. & Minh, B. Q. IQ-TREE: A fast and effective stochastic algorithm for estimating maximum-likelihood phylogenies. Mol. Biol. Evol. 32, 268–274 (2015).

Okamura, A. et al. Artificial hybrid between *Anguilla anguilla* and A. japonica. J. Fish Biol. 64, 1450–1454 (2004).

Orr, H. A. & Turelli, M. The evolution of postzygotic isolation: Accumulating Dobzhansky-Muller incompatibilities. Evolution 55, 1085–1094 (2001).

Patterson, N, Price, A. L. & Reich D. Population structure and eigenanalysis. PLoS Genet. 2, e190 (2006).

Peterson, B. K. Weber, J. N. Kay, E. H. Fisher, H. S. & Hoekstra, H. E. Double digest RADseq: An inexpensive method for de novo SNP discovery and genotyping in model and non-model species. PLoS ONE 7, e37135 (2012).

Prager, E. M. & Wilson, A. C. Slow evolutionary loss of the potential for interspecific hybridization in birds: A manifestation of slow regulatory evolution. Proc. Natl. Acad. Sci. USA 72, 200–204 (1975).

Price, T. D. & Bouvier, M. The evolution of F1 postzygotic incompatibilities in birds. Evolution 56, 2083–2089 (2002).

Pujolar, J. M. et al. Assessing patterns of hybridization between North Atlantic eels using diagnostic single-nucleotide polymorphisms. Heredity 112, 627–637 (2014).

Pujolar, J. M. & Maes, G. in Biology and Ecology of Anguillid Eels (ed Arai, T.) 36–51 (CRC Press, Boca Raton, Florida, 2016).

Purcell, S. et al. PLINK: a tool set for whole-genome association and population-based linkage analyses. Am. J. Human Genet. 81, 559–575 (2007).

Quinlan, A. R. & Hall, I. M. BEDTools: a flexible suite of utilities for comparing genomic features. Bioinformatics 26, 841–842 (2010).

Rabosky, et al. An inverse latitudinal gradient in speciation rate for marine fishes. Nature 559, 392–395 (2018).

Rambaut, A. Drummond, A. J. Xie, D. Baele, G. & Suchard, M. A. Posterior summarization in Bayesian phylogenetics using Tracer 1.7. Syst. Biol. 67, 901–904 (2018).

Reich, D. Thangaraj, K. Patterson, N. Price, A. L. & Singh, L. Reconstructing Indian population history. Nature 461, 489–494 (2009).

Rieseberg, L. H. Archer M. A. & Wayne R. K. Transgressive segregation, adaptation and speciation. Heredity 83, 363–372 (1999).

Rochette, N. C. & Catchen, J. M. Deriving genotypes from RAD-seq short-read data using Stacks. Nat. Protoc. 12, 2640–2659 (2017).

Runemark, A. et al. Variation and constraints in hybrid genome formation. Nat. Ecol. Evol. 2, 549–556 (2018).

Sambrook, J. Fritsch, E. F. & Maniatis, T. Molecular Cloning: A Laboratory Manual, 2nd ed. (Cold Spring Harbor Laboratory Press, Cold Spring Harbor, NY, 1989).

Schabetsberger, R. et al. Genetic and migratory evidence for sympatric spawning of tropical Pacific eels from Vanuatu. Marine Ecol. Prog. Ser. 521, 171–187 (2015).

Schabetsberger, R. et al. Hydrographic features of anguillid spawning areas: Potential signposts for migrating eels. Mar. Ecol. Prog. Ser. 554, 141–155 (2016).

Schumer, M. Cui, R. Powell, D. L. Rosenthal, G. G. & Andolfatto, P. Ancient hybridization and genomic stabilization in a swordtail fish. Mol. Ecol. 25, 2661–2679 (2016).

Stamatakis, A. RAxML version 8: a tool for phylogenetic analysis and post-analysis of large phylogenies. Bioinformatics 30, 1312–1313 (2014).

Stange, M. Sánchez-Villagra, M. R. Salzburger, W. & Matschiner, M. Bayesian divergence-time estimation with genome-wide SNP data of sea catfishes (Ariidae) supports Miocene closure of the Panamanian Isthmus. Syst. Biol. 67, 681–699 (2018).

Stelkens, R. B. Schmid, C. Selz, O. & Seehausen, O. Phenotypic novelty in experimental hybrids is predicted by the genetic distance between species of cichlid fish. BMC Evol. Biol. 9, 283 (2009).

Stelkens, R. B. Schmid, C. & Seehausen, O. Hybrid breakdown in cichlid fish. PLoS ONE 10, e0127207 (2015).

Stelkens, R. B. & Seehausen, O. Genetic distance between species predicts novel trait expression in their hybrids. Evolution 63, 884–897 (2009).

Stelkens, R. B. Young, K. A. & Seehausen, O. The accumulation of reproductive incompatibilities in African cichlid fish. Evolution 64, 617–633 (2010).

Taylor, S. A. & Larson E. L. Insights from genomes into the evolutionary importance and prevalence of hybridisation in nature. Nature Ecol. Evol. 3, 170–177 (2019).

Teng, H.-Y. Lin, Y.-S. & Tzeng C.-S. A new *Anguilla* species and a reanalysis of the phylogeny of freshwater eels. Zool. Studies 48, 808–822 (2009).

Tørresen, O. K. et al. An improved genome assembly uncovers prolific tandem repeats in Atlantic cod. BMC Genomics 18, 95 (2017).

Tseng, M.-C. in Biology and Ecology of Anguillid Eels (ed Arai, T.) 21–35 (CRC Press, Boca Raton, Florida, 2016).

Turelli, M. & Moyle L. C. Asymmetric postmating isolation: Darwin’s corollary to Haldane’s rule. Genetics 176, 1059–1088 (2007).

Walker, B. J. et al. Pilon: An integrated tool for comprehensive microbial variant detection and genome assembly improvement. PLoS ONE 9, e112963 (2014).

Watanabe, S. Aoyama, J. & Tsukamoto, K. Reexamination of Ege’s (1939) use of taxonomic characters of the genus Anguilla. Bull. Mar. Sci. 74, 337–351 (2004).

Watanabe, S. et al. Evidence of population structure in the giant mottled eel, *Anguilla marmorata*, using total number of vertebrae. Copeia 2008, 680–688 (2008).

Watanabe, S. Miller, M. J. Aoyama, J. & Tsukamoto, K. Morphological and meristic evaluation of the population structure of *Anguilla marmorata* across its range. J. Fish Biol. 74, 2069–2093 (2009).

Waterhouse, R. M. et al. BUSCO applications from quality assessments to gene prediction and phylogenomics. Mol. Biol. Evol. 35, 543–548 (2018).

Watterson, G. A. On the number of segregating sites in genetical models without recombination. Theor. Popul. Biol. 7, 256–276 (1975).

Wielgoss, S. Meyer, A. Gilabert, A. & Wirth T. Introgressive hybridization and latitudinal admixture clines in North Atlantic eels. BMC Evol. Biol. 14, 61 (2014).

Wiley, C. Qvarnström, A. Andersson, G. Borge, T. & Saetre, G.-P. Postzygotic isolation over multiple generations of hybrid descendents in a natural hybrid zone: how well do single-generation estimates reflect reproductive isolation? Evolution 63, 1731–1739 (2009).

