## Supplementary Information for "Stable Species Boundaries Despite Ten Million Years of Hybridization in Tropical Eels"

**Addresses:**
[1]Zoological Institute, University of Basel, Basel, Switzerland.
[2]Fisheries Research Institute, Hellenic Agricultural Organisation-DEMETER, Nea Peramos, Kavala, Greece.
[3]Department of Palaeontology and Museum, University of Zurich, Zurich, Switzerland.
[4]Centre for Ecological and Evolutionary Synthesis, Department of Biosciences, University of Oslo, Oslo, Norway.
[5]Faculty of Agriculture, Kindai University, Nara, Japan.
[6]Institute of Fisheries Science, College of Life Science, National Taiwan University, Taipei, Taiwan.
[7]UMR Biologie des Organismes et Écosystèmes Aquatiques (BOREA UMR 7208), Muséum National d'Histoire Naturelle, Sorbonne Université, Université Pierre et Marie Curie, Université de Caen Normandie, Université des Antilles, CNRS, IRD, Paris, France.
[8]MNHN - Station Marine de Dinard, Centre de Recherche et d'Enseignement Sur les Systèmes Côtiers (CRESCO), Dinard, France.
[9]Department of Ecology, University of Innsbruck, Innsbruck, Austria.
[10]School of Environment and Life Sciences, University of Salford, Salford, UK.
[11]Department of Biosciences, University of Salzburg, Salzburg, Austria.
[12]Present address: Center for Active Learning (CAL), Department of Biology, ETH Zurich, Zurich, Switzerland.
[13]These first authors contributed equally: Julia M. I. Barth, Chrysoula Gubili, Michael Matschiner.
[14]These senior authors contributed equally: Robert Jehle, Robert Schabetsberger.

### 1 Supplementary Notes

**Supplementary Note 1:** Contrasting patterns of within-species genomic variation.
Our extensive sampling scheme of tropical eels permitted detailed analyses of genomic variation within *A. marmorata*, *A. megastoma* and *A. obscura*, as we sampled each of the three species at multiple sites throughout their geographic distribution (Fig. 1a; Supplementary Table 1). These analyses were based on a dataset of 155,896 RAD-sequencing derived single-nucleotide polymorphisms (SNPs), partitioned according to species and subsequently filtered to exclude invariant sites (minor allele count $> 2$) and missing data ($> 20\%$; Supplementary Figure 1). Using these partitioned datasets, principal-component analysis (PCA) of genomic variation was performed with smartPCA in EIGENSOFT v.6.0.1 (Patterson *et al.* 2006), including the function "lsqproject" to account for missing data. For *A. marmorata*, PCA separated four populations present in the western Indian Ocean (South Africa, Reunion, Mayotte), in Indonesia (Java), the South China Sea (Philippines and Taiwan), and the western South Pacific (Bougainville Island, Solomon Islands, Vanuatu, New Caledonia, Samoa, and American Samoa). The latter three, however, were only discernible on the second principal-component axis (explaining 2.2% of genetic variation), along which the individuals from Java appeared intermediate between those from the western Indian Ocean and the western South Pacific (Supplementary Figure 4a). Our results are thus consistent with divergence among Indian and Pacific ocean populations (Ishikawa *et al.* 2004; Minegishi *et al.* 2008; Watanabe *et al.* 2008; Gagnaire *et al.* 2011) and with the region of Java representing a contact zone between those populations (Gagnaire *et al.* 2011). Conversely, no population structure was detected in either *A. megastoma* or *A. obscura* (Supplementary Figures 4c-f), supporting the hypothesized single spawning area for the two species in the western South Pacific (Schabetsberger *et al.* 2015, 2016).

**Supplementary Note 2:** Double-digest restriction-site associated DNA (ddRAD) sequencing.
Following Peterson *et al.* (2012), 20 units of EcoRI-HF (New England Biolabs) and 20 units of MspI (New England Biolabs) were used to digest 400 ng of genomic DNA per sample in a 37 °C incubation for 8 hours. Digests were purified with homemade paramagnetic carboxyl-modified beads (Sera-Mag, Fisher Scientific; Rohland & Reich 2012). Moreover, samples were randomly placed in PCR plates, ensuring a wide coverage of geographic sampling locations per plate during library preparation. T4 DNA ligase (New England Biolabs) was applied to ligate 100 ng of each digested DNA fragment to a EcorRI-specific P1 adapter that contained a 5-bp barcode and the MspI-specific P2 adapter in room temperature, followed by an enzyme heat-kill at 65 °C for 10 min. Twenty-four unique barcodes were used so that the ligated DNA fragments from 24 individuals could be pooled (according to one index) to form a single ddRAD-seq library. Ligations were cleaned with homemade paramagnetic carboxyl-modified beads. Fragments in the range of 300-400 bp were selected using AMPure XP beads (Agilent Technologies) and were subsequently amplified by 12 rounds of PCR with the following conditions: 98 °C for 60 s; 12 cycles of 98 °C for 10 s, 60 °C for 30 s, 72 °C for 30 s; 72 °C for 10 min using Q5® High Fidelity polymerase (New England Biolabs), and the Illumina sequencing primers (PCR Primer 1 and Index added PCR Primer 2; four unique 6-bp multiplexing indices were used). In total, six separate PCR reactions of the same index were amplified, pooled

and cleaned with homemade paramagnetic carboxyl-modified beads. Library quality was assessed on a TapeStation 2200 (Agilent Technologies) to confirm fragment recovery on the selected range and was quantified using a Qubit Fluorometer 2.0. To avoid index-hopping (Kircher *et al.* 2012; Sinha *et al.* 2018), none of the samples shared both the P1 barcode and the multiplexing index in a given sequencing run. In total, 20 libraries, each containing 24 barcodes, were sent to Macrogen (Korea) for 100 bp paired-end Illumina HiSeq 4000 sequencing.

**Supplementary Note 3:** The reliability of published age estimates for the genus *Anguilla*.
We time calibrated the species tree of tropical eels according to age estimates reported by Jacobsen *et al.* (2014) on the basis of mitochondrial genomes. In their study, Jacobsen *et al.* (2014) used the earliest fossil records of the family Anguillidae, *Eoanguilla leptoptera* from Monte Bolca, Italy (Patterson 1993; Carnevale *et al.* 2014), to constrain the divergence between Anguillidae and Serrivomeridae to 55-50 Ma. Given that the age of the Monte Bolca deposits is 49.4-49.1 Ma (Benton *et al.* 2015; Matschiner *et al.* 2017), it is indeed likely that Anguillidae originated before 50 Ma. However, since fossils do not directly constrain maximum ages, the upper boundary of 55 Ma was arbitrarily specified by Jacobsen *et al.* (2014), and the divergence times estimated in their study would likely be underestimated if Anguillidae in fact originated earlier than 55 Ma. Nevertheless, we consider the timeline proposed by Jacobsen *et al.* (2014) plausible for the following reasons: (i) According to this timeline, European and American anguillid species (*A. anguilla* and *A. rostrata*) diverged from Indo-Pacific members of the genus around 10.8 Ma, which is consistent with the Messinian age (7.2-5.3 Ma) of the earliest fossils of the genus, known from the Gessoso Solififera Formation in Northern Italy (Dela Pierre *et al.* 2011); (ii) the timeline is consistent with those of two other recent studies based on genome-wide data (Musilova *et al.* 2019) and a massive taxon set (Rabosky *et al.* 2018), as the most recent common ancestor of *A. anguilla* and *A. japonica* (the only species pair included in all three studies) was estimated at 13.8 Ma in Jacobsen *et al.* (2014), at 12.4 Ma in Musilova *et al.* (2019), and at 12.9 Ma in Rabosky *et al.* (2018).

**Supplementary Note 4:** Assessing the robustness of divergence-time estimates.
To test how robust the divergence-time estimates are to alternative phylogenetic positions of *A. interioris*, SNAPP analyses were repeated separately with two fixed topologies in which *A. interioris* is either the sister of *A. bicolor* and *A. obscura* or the sister to the clade formed by *A. marmorata*, *A. luzonensis*, *A. bicolor*, and *A. obscura*. Furthermore, to test the robustness of divergence-time estimates to introgression involving *A. luzonensis* and *A. interioris*, the analyses were repeated after excluding these two species.

Additionally, we compiled a multi-locus phylogenetic dataset based on genome assemblies of the five species *A. anguilla*, *A. japonica*, *A. marmorata*, *A. obscura*, and *A. megastoma* to estimate divergence times among *Anguilla* species independently of the timeline of Jacobsen *et al.* (2014). Of these five species, genome assemblies of *A. anguilla* (NCBI accession GCA_000695075; Henkel *et al.* 2012a) and *A. japonica* (NCBI accession GCA_000470695; Henkel *et al.* 2012b) were included in the large-scale phylogenomic analysis of Musilova *et al.* (2019), in which their divergence was estimated at around 12.37 Ma. We thus extracted ortholog sequences, corresponding to the loci used in Musilova *et al.* (2019), from the three new genome assemblies of *A. marmorata*, *A. obscura*, and *A. megastoma* (Supplementary Table 5), and aligned these jointly with those of *A. anguilla* and *A. japonica*. Alignments were then filtered according to the protocol of Musilova *et al.* (2019), excluding 10 of the 113 genes used in Musilova *et al.* (2019) due to missing sequences. Alignments for the remaining 103 nuclear genes were concatenated and split into two separate partitions for first- and second-codon positions; third-codon positions were excluded. Together, these two partitions included 92,530 bp with 0.07% missing data. The concatenated alignment was used for phylogenetic analyses with the software BEAST 2, time calibrating the phylogeny according to the timeline estimated by Musilova *et al.* (2019). Specifically, we used 12.37 Ma, the estimated age for the divergence of *A. anguilla* and *A. japonica*, as a constraint on the age of the most recent common ancestor of the five *Anguilla* species, after initial analyses suggested an position of *A. anguilla* outside of a clade formed by the other four species. The model used in this analysis included a GTR substitution-rate matrix (Tavaré 1986), gamma-distributed among-site rate variation, a strict molecular clock, and the Yule process of species diversification (Yule 1925). The BEAST 2 analysis was performed with 10 million MCMC iterations. Convergence was again assessed with Tracer, and the posterior tree distribution was summarized in an MCC tree generated with TreeAnnotator.

As another alternative to the divergence-time estimates of Jacobsen *et al.* (2014), we also implemented age constraints according to the timeline of Rabosky *et al.* (2018). The large-scale time-calibrated phylogeny of Rabosky *et al.* (2018), based on molecular data for 11,638 ray-finned fishes and 139 fossil constraints, includes 20 species and subspecies of the genus *Anguilla* and places the earliest divergence within the genus, the separation of *A. australis*, at 21.57 Ma. As the genus was mostly (82%) represented by mitochondrial sequences in the dataset of Rabosky *et al.* (2018), we extracted homologous mitochondrial sequences from the new genome assemblies of *A. marmorata*, *A. megastoma*, and *A. obscura* (Supplementary Table 5) using BLAST, and integrated these sequences with the mitochondrial data for *Anguilla* compiled by Rabosky *et al.* (2018). This integrated mitochondrial dataset was concatenated into a single alignment and used again for phylogenetic inference with BEAST 2. According to the phylogeny of Rabosky *et al.* (2018), *A. australis* was constrained to be the outgroup of the other *Anguilla* species and the divergence of *A. australis* was fixed at 21.57 Ma. The model used in this analysis was identical to that used for nuclear data, except that the GTR substitution-rate matrix was replaced with an HKY matrix (Hasegawa *et al.* 1985) because some GTR rate parameters appeared unidentifiable in preliminary analyses. The BEAST 2 analysis was again performed for 10 million MCMC iterations, convergence was assessed with Tracer, and a MCC summary tree was generated with TreeAnnotator.

#### 2 Supplementary Figures

**Supplementary Figure 1:** Molecular datasets used in this study.
Flow chart illustrating how RAD sequencing data was filtered and used for various analyses. File names are shown in yellow and analyses are highlighted in cyan.

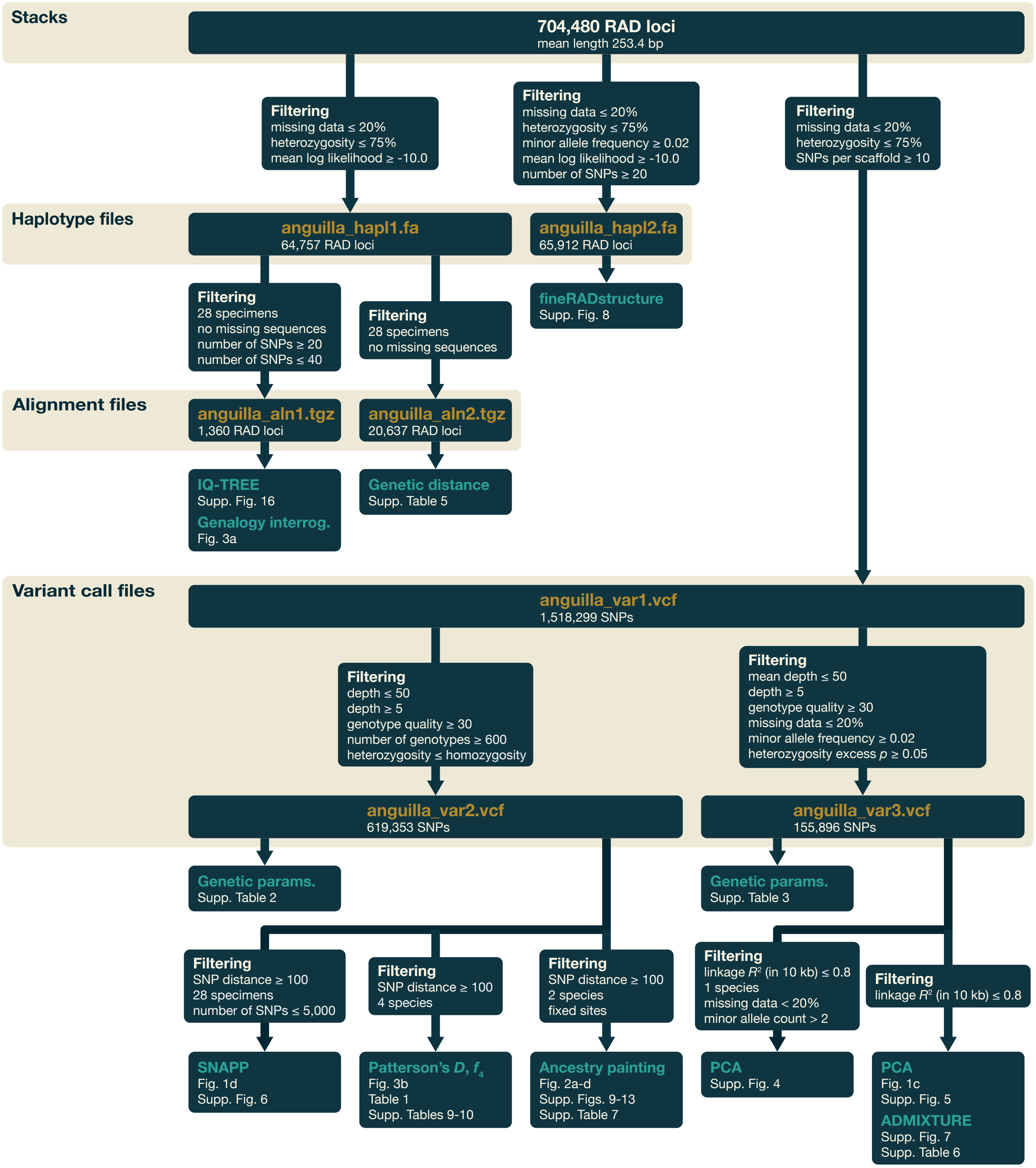

**Supplementary Figure 2:** Haplotype-genealogy graph based on mitochondrial sequences. Graph generated with the software Fitchi v.1.1.4 (Matschiner 2016) for two concatenated RAD loci mapping to positions 10630-10720 and 12015-12105 of the *Anguilla japonica* mitochondrial genome (NCBI accession CM002536). The genealogy of mitochondrial sequences for all 456 individuals was produced using RAxML v. 8.2.11 (Stamatakis 2014) with the GTRCAT model of sequence evolution.

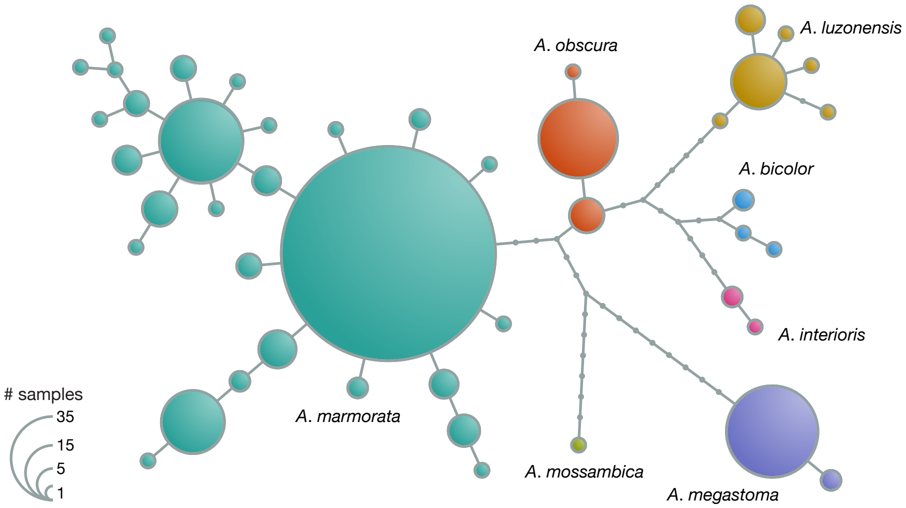

**Supplementary Figure 3:** Morphological variation among tropical eel species.
a) Following Watanabe *et al.* (2009), the predorsal length without the head (PDH) and the distance between the dorsal fin and the anus (AD) were measured for 161 individuals available for morphological analyses (100 × *A. marmorata*, 30 × *A. megastoma*, 30 × *A. obscura*, and 1 × *A. interioris*) and standardized by terminal length (TL). Color code is identical to Supplementary Figure 2. Individuals selected as putatively unadmixed "core" group representatives of *A. marmorata*, *A. megastoma*, and *A. obscura* are marked with dark gray outlines. Specimen IDs (see Supplementary Table 1) are given for putative hybrids and one representative of *A. interioris*. b) First and second principal components of morphological variation. "Core" individuals were selected according to clusters shown in this plot.

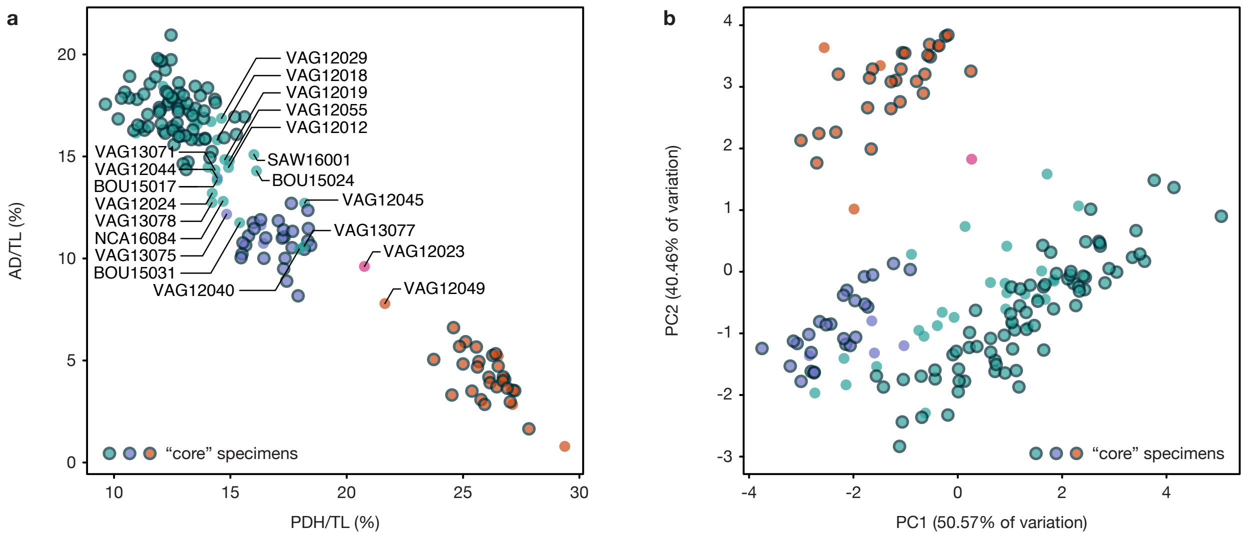


**Supplementary Figure 4 (next page):** Genomic variation within tropical eel species.
a-b) Comparison of first and second (a), and third and fourth (b), principal components of genomic variation in *A. marmorata*. c-d) Comparison of first and second (c), and third and fourth (d), principal components of genomic variation in *A. megastoma*. e-f) Comparison of first and second (e), and third and fourth (f), principal components of genomic variation in *A. obscura*. Putative between-species hybrids (see Supplementary Table 7) were exluded from this analysis. Stroke color indicates geographic origin as specified in Supplementary Figure 3.

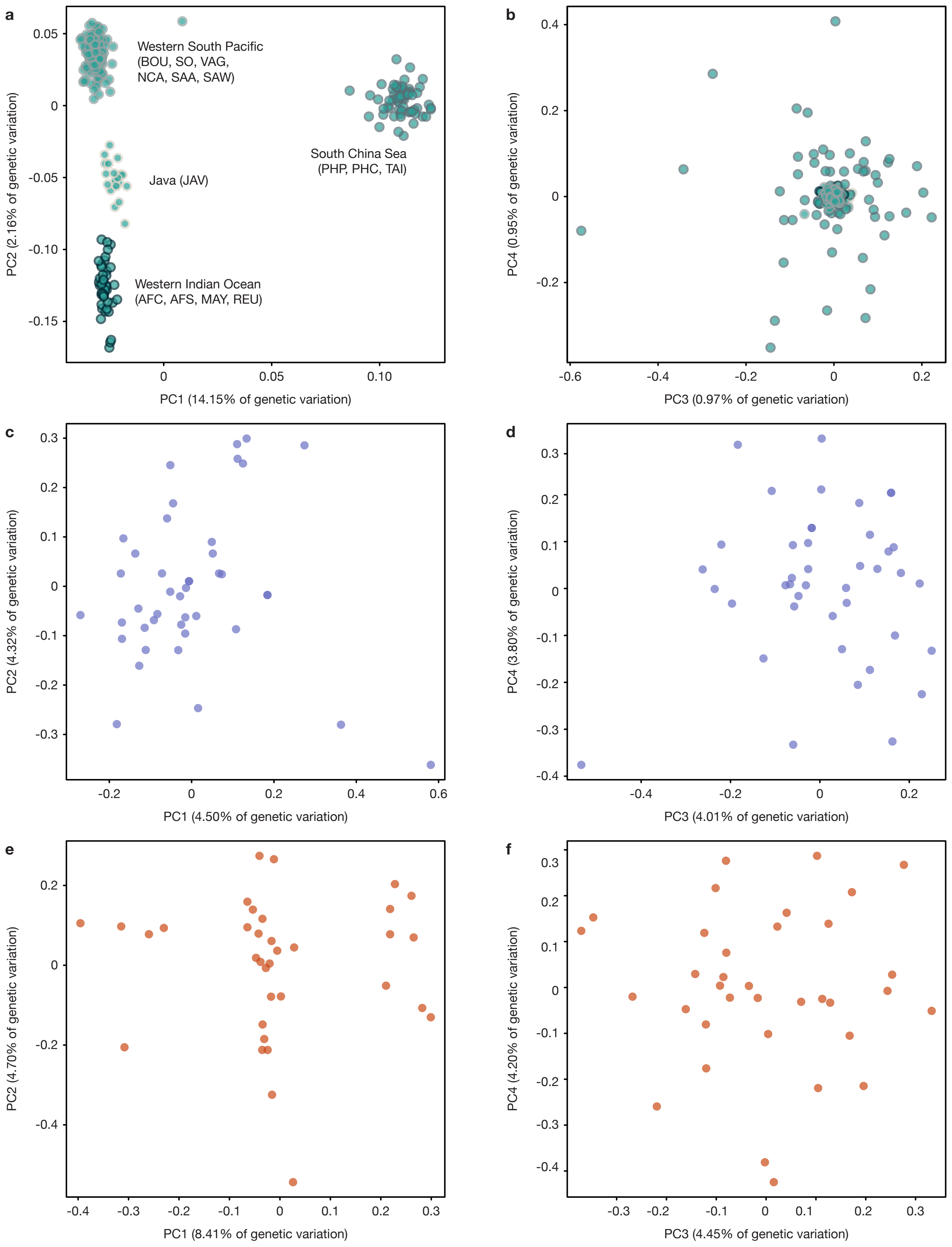
a
b
c
d
e
f
Western South Pacific
(BOU, SO, VAG,
NCA, SAA, SAW)
Java (JAV)
Western Indian Ocean
(AFC, AFS, MAY, REU)
South China Sea
(PHP, PHC, TAI)
PC1 (14.15% of genetic variation)
PC2 (2.16% of genetic variation)
PC3 (0.97% of genetic variation)
PC4 (0.95% of genetic variation)
PC1 (4.50% of genetic variation)
PC2 (4.32% of genetic variation)
PC3 (4.01% of genetic variation)
PC4 (3.80% of genetic variation)
PC1 (8.41% of genetic variation)
PC2 (4.70% of genetic variation)
PC3 (4.45% of genetic variation)
PC4 (4.20% of genetic variation)

**Supplementary Figure 5:** Genomic variation among tropical eel species.
a) First and second principal components of genomic variation among the seven species *A. marmorata* (cyan), *A. luzonensis* (brown), *A. megastoma* (purple), *A. obscura* (red), *A. bicolor* (blue), *A. interioris* (magenta), and *A. mossambica* (green). b) Third and fourth principal components of genomic variation among the seven species. c) First and second principal components of genomic variation, focusing on the four species *marmorata* (cyan), *A. megastoma* (purple), *A. obscura* (red), and *A. interioris* (magenta). d) Third and fourth principal components of genomic variation, for the same four species as in c). The two individuals VAG12033 and PHP14P16 are characterized by a large proportion of missing data (see Supplementary Table 1), which might explain their outlier positions.

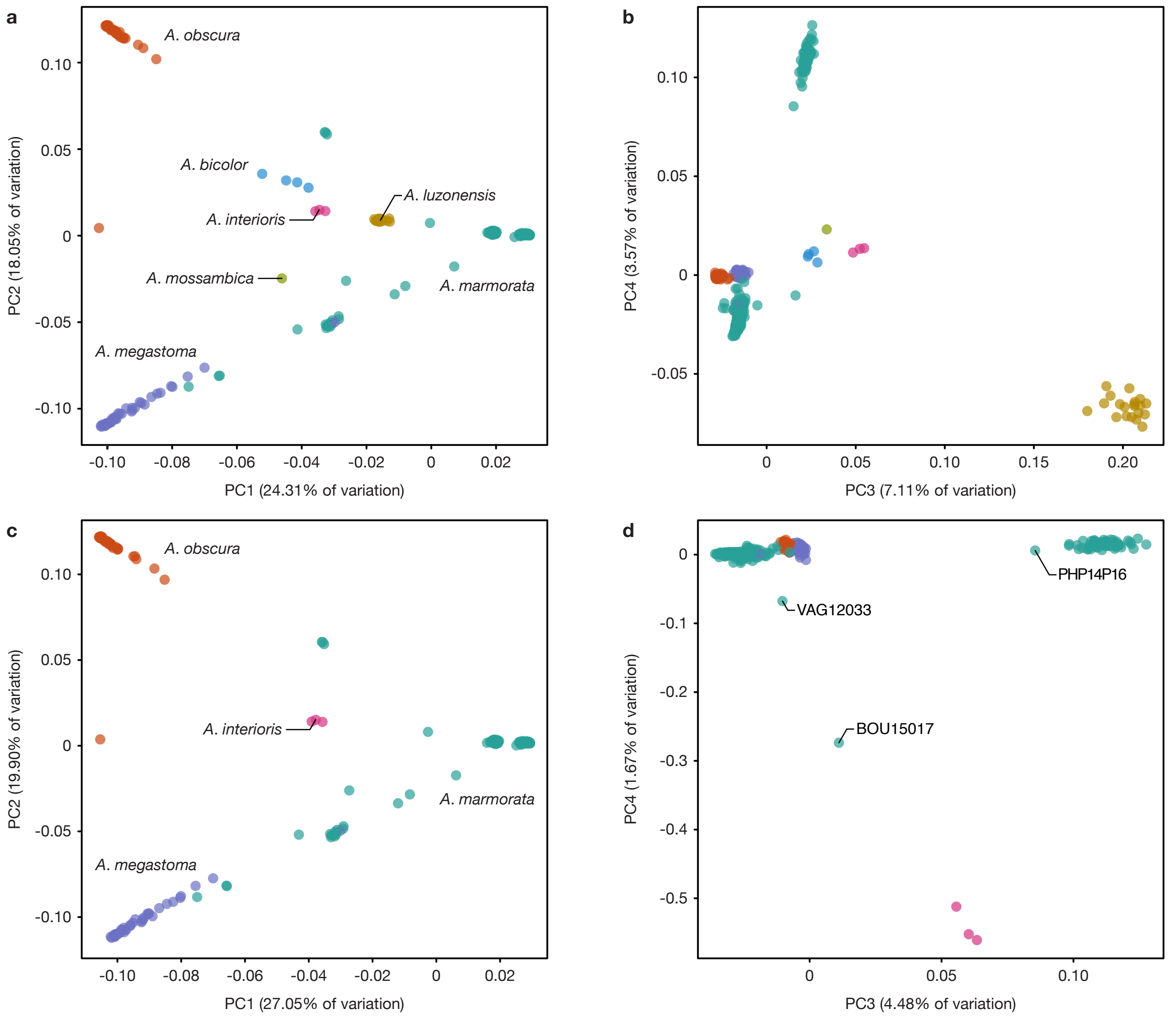

**Supplementary Figure 6 (next page):** Divergence times of tropical eel species.
a-b) Maximum-clade-credibility (MCC) summary trees of SNAPP analyses with 5,000 transition or transversion sites, respectively, without topology constraints and a single age constraint on the root divergence according to Jacobsen *et al.* (2014) (J2014). The gray area marks the group of species for which we find evidence of past and ongoing hybridization, and the dotted line indicates the crown age of this group. c-d) As a-b), but with a topology constraint on the position of *A. interioris* as the sister species to a clade combining *A. bicolor* and *A. obscura*; this position is supported by maximum-likelihood inference with IQ-TREE (Supplementary Figure 17). e-f) As a-b), but with a topology constraint on the position of *A. interioris* as the outgroup to a formed by *A. bicolor*, *A. obscura*, *A. marmorata*, and *A. luzonensis*; this position is supported by genealogy interrogation (Fig. 3a). g-h) As a-b), but excluding the the two species *A. luzonensis* and *A. interioris* due to their strong signals of ancient hybridization. i) MCC tree of a BEAST analysis with 103 nuclear genes. Sequences of these genes from *A. anguilla* and *A. japonica* were included in Musilova *et al.* (2019) (M2019) and were here complemented with orthologs extracted from the new genome assemblies of *A. marmorata*, *A. megastoma*, and *A. obscura*. A single age constraint on the root was used for calibration according to Musilova *et al.* (2019). j) MCC tree of a BEAST analysis with four mitochondrial genes that were used in Rabosky *et al.* (2018) (R2018). The dataset of Rabosky *et al.* (2018) was here complemented with orthologs from the three new genome assemblies (*A. marmorata*, *A. megastoma*, and *A. obscura*), and a single age constraint on the root was used according to Rabosky *et al.* (2018). Note that the timescale in j) differs from a-i). Unless specified, all nodes received full Bayesian support.

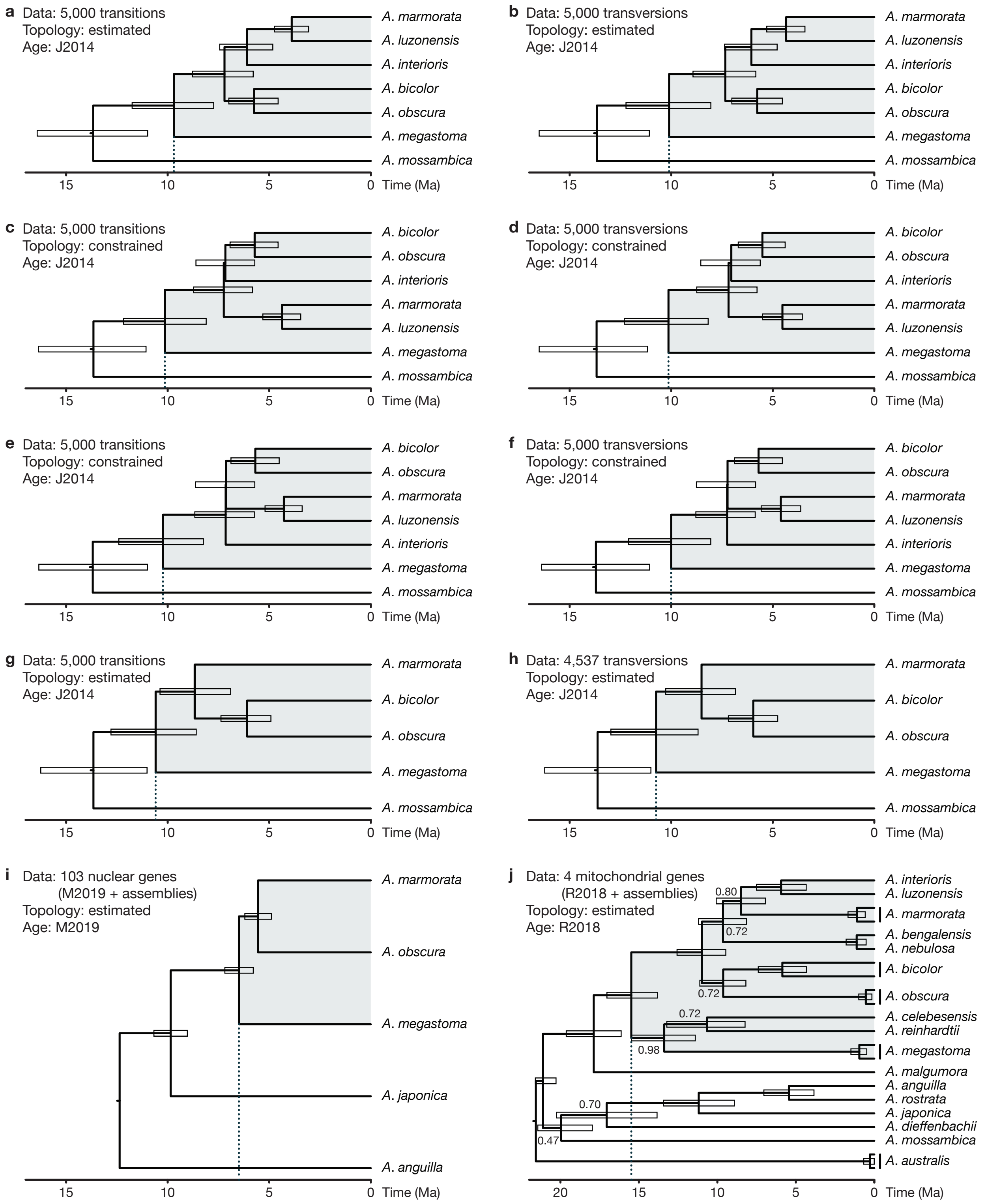
a
Data: 5,000 transitions
Topology: estimated
Age: J2014
A. marmorata
A. luzonensis
A. interioris
A. bicolor
A. obscura
A. megastoma
A. mossambica
15
10
5
0
Time (Ma)
b
Data: 5,000 transversions
Topology: estimated
Age: J2014
A. marmorata
A. luzonensis
A. interioris
A. bicolor
A. obscura
A. megastoma
A. mossambica
15
10
5
0
Time (Ma)
c
Data: 5,000 transitions
Topology: constrained
Age: J2014
A. bicolor
A. obscura
A. interioris
A. marmorata
A. luzonensis
A. megastoma
A. mossambica
15
10
5
0
Time (Ma)
d
Data: 5,000 transversions
Topology: constrained
Age: J2014
A. bicolor
A. obscura
A. interioris
A. marmorata
A. luzonensis
A. megastoma
A. mossambica
15
10
5
0
Time (Ma)
e
Data: 5,000 transitions
Topology: constrained
Age: J2014
A. bicolor
A. obscura
A. marmorata
A. luzonensis
A. interioris
A. megastoma
A. mossambica
15
10
5
0
Time (Ma)
f
Data: 5,000 transversions
Topology: constrained
Age: J2014
A. bicolor
A. obscura
A. marmorata
A. luzonensis
A. interioris
A. megastoma
A. mossambica
15
10
5
0
Time (Ma)
g
Data: 5,000 transitions
Topology: estimated
Age: J2014
A. marmorata
A. bicolor
A. obscura
A. megastoma
A. mossambica
15
10
5
0
Time (Ma)
h
Data: 4,537 transversions
Topology: estimated
Age: J2014
A. marmorata
A. bicolor
A. obscura
A. megastoma
A. mossambica
15
10
5
0
Time (Ma)
i
Data: 103 nuclear genes
(M2019 + assemblies)
Topology: estimated
Age: M2019
A. marmorata
A. obscura
A. megastoma
A. japonica
A. anguilla
15
10
5
0
Time (Ma)
j
Data: 4 mitochondrial genes
(R2018 + assemblies)
Topology: estimated
Age: R2018
0.80
0.72
0.72
0.72
0.98
0.70
0.47
A. interioris
A. luzonensis
A. marmorata
A. bengalensis
A. nebulosa
A. bicolor
A. obscura
A. celebesensis
A. reinhardtii
A. megastoma
A. malgumora
A. anguilla
A. rostrata
A. japonica
A. dieffenbachii
A. mossambica
A. australis
20
15
10
5
0
Time (Ma)

**Supplementary Figure 7:** Maximum-likelihood ancestry inference.
Ancestry proportions displayed in bars per individual were inferred for the models $K = 1$ to $K = 8$ based on 117,638 variable sites using the software ADMIXTURE (Alexander *et al.* 2009). The inset shows the cross-validation (CV) error for five replicates per model. For *A. marmorata*, individuals are labeled according to geographic origin: Indian Ocean (sampling locations AFC, AFS, MAY, REU), Java, South China Sea (PHP, PHC, TAI), and Pacific (BOU, NCA, SAA, SAW, SO, VAG). Label 1 marks 20 individuals that appear admixed between *A. marmorata* and *A. megastoma*: BOU15031, SAA16011, SAA16012, SAA16013, SAA16024, SAA16027, SAW17B27, SAW17B49, VAG12012, VAG12018, VAG12019, VAG12024, VAG12029, VAG12037, VAG12044, VAG12053, VAG12055, VAG13071, VAG13078, and VAG13087. Labels 2-4 indicate species *A. luzonensis* (2), *A. bicolor* (3), and *A. mossambica* (4).

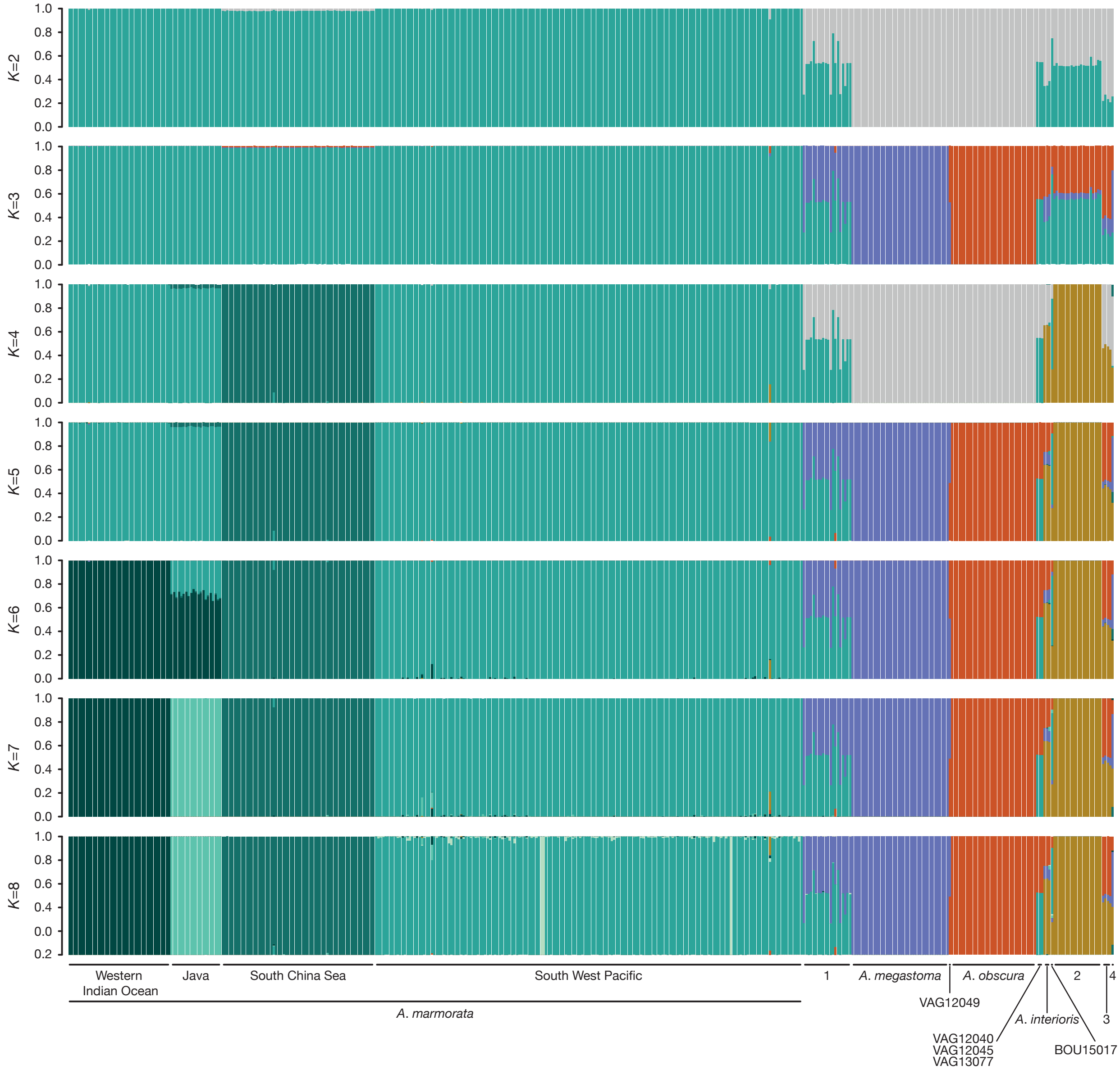

**Supplementary Figure 8:** Individual coancestry based on haplotype similarity.
Coancestry was investigated based on RAD loci with fineRADstructure (Malinsky *et al.* 2018). Heatmap colors indicate numbers of RAD loci with estimated shared coancestry. Individuals are listed on both axes in the same order, clustered according to the tree shown on top of the heatmap (Lawson *et al.* 2012). Note that even though *A. luzonensis* appears to have more shared coancestry with the South China Sea population of *A. marmorata* than with other populations, introgression between *A. luzonensis* and the South China Sea population of *A. marmorata* is not supported by $D$ and $f_4$ statistics (Supplementary Table 10).

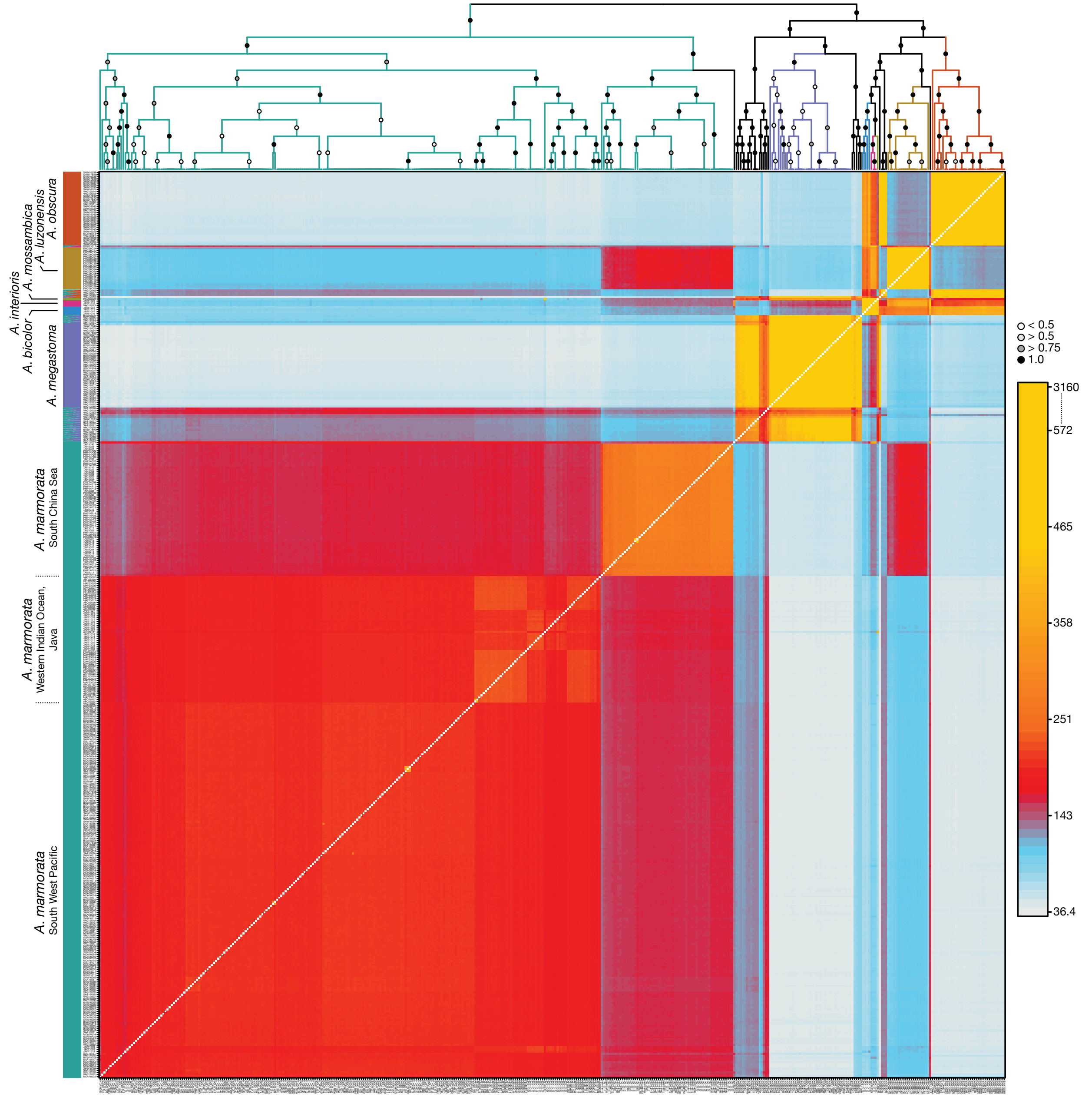

**Supplementary Figure 9 (next page):** Ancestry painting for *A. marmorata* and *A. megastoma*. Ancestry painting for 73 "core" *A. marmorata* individuals, 26 "core" *A. megastoma* individuals, and 20 recent hybrids between the two species. In addition, one *A. marmorata* individual (BOU15024) was included because it was initially assumed to be a hybrid based on morphological measurements (Supplementary Figure 3a); this assumption is not supported by the ancestry painting shown here. Horizontal bars indicate the genotypes at each of 302 sites fixed between the two parental species. White color indicates missing data. Heterozygous genotypes are shown with the top half in each bar matching the second parental species and vice versa. Light gray cells in the morphology column indicate individuals not classified into any of the "core" groups. The species' color code is identical to Supplementary Figures 1-4.

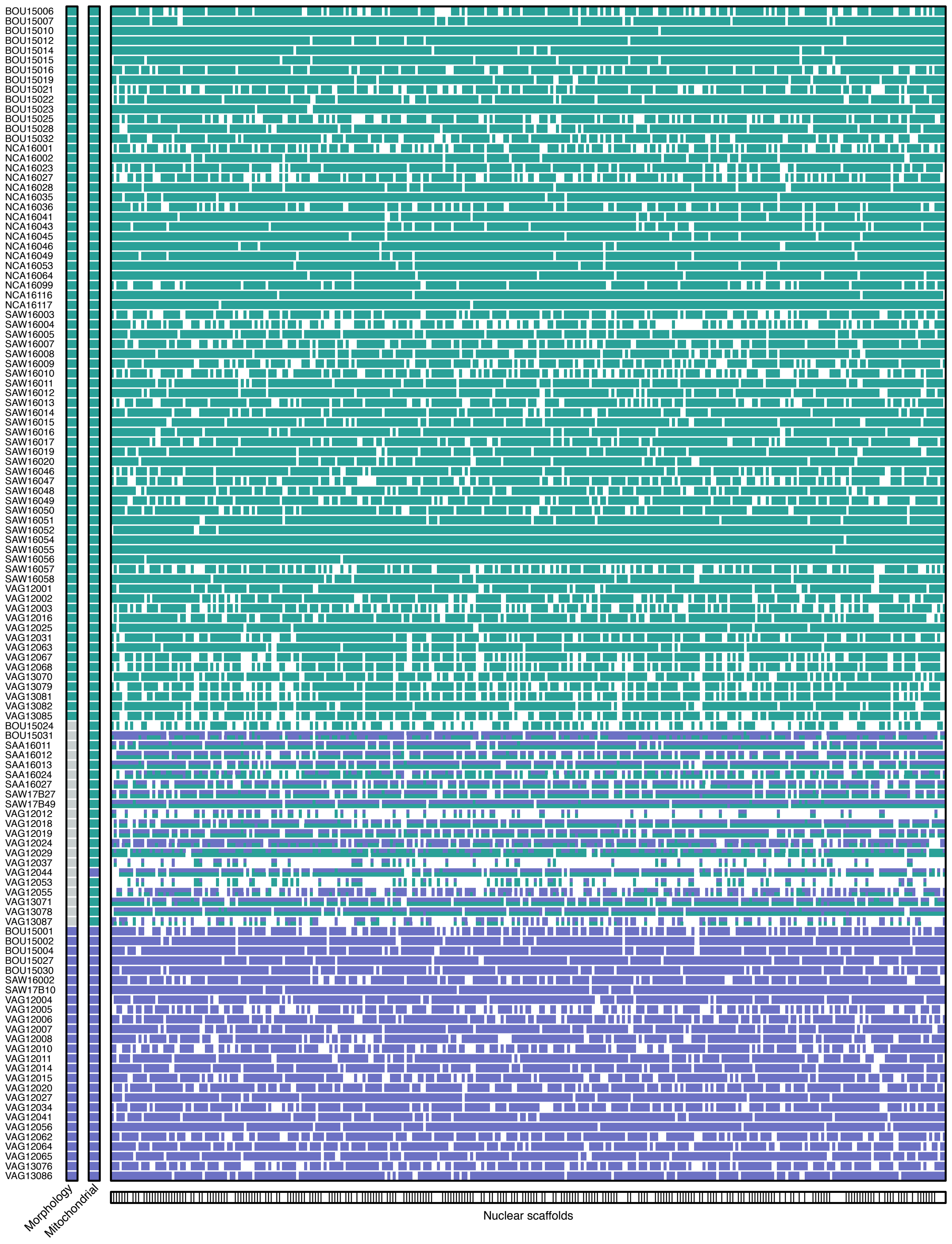
BOU15006
BOU15007
BOU15010
BOU15012
BOU15014
BOU15015
BOU15016
BOU15019
BOU15021
BOU15022
BOU15023
BOU15025
BOU15028
BOU15032
NCA16001
NCA16002
NCA16023
NCA16027
NCA16028
NCA16035
NCA16036
NCA16041
NCA16043
NCA16045
NCA16046
NCA16049
NCA16053
NCA16064
NCA16099
NCA16116
NCA16117
SAW16003
SAW16004
SAW16005
SAW16007
SAW16008
SAW16009
SAW16010
SAW16011
SAW16012
SAW16013
SAW16014
SAW16015
SAW16016
SAW16017
SAW16019
SAW16020
SAW16046
SAW16047
SAW16048
SAW16049
SAW16050
SAW16051
SAW16052
SAW16054
SAW16055
SAW16056
SAW16057
SAW16058
VAG12001
VAG12002
VAG12003
VAG12016
VAG12025
VAG12031
VAG12063
VAG12067
VAG12068
VAG13070
VAG13079
VAG13081
VAG13082
VAG13085
BOU15024
BOU15031
SAA16011
SAA16012
SAA16013
SAA16024
SAA16027
SAW17B27
SAW17B49
VAG12012
VAG12018
VAG12019
VAG12024
VAG12029
VAG12037
VAG12044
VAG12053
VAG12055
VAG13071
VAG13078
VAG13087
BOU15001
BOU15002
BOU15004
BOU15027
BOU15030
SAW16002
SAW17B10
VAG12004
VAG12005
VAG12006
VAG12007
VAG12008
VAG12010
VAG12011
VAG12014
VAG12015
VAG12020
VAG12027
VAG12034
VAG12041
VAG12056
VAG12062
VAG12064
VAG12065
VAG13076
VAG13086
Morphology
Mitochondrial
Nuclear scaffolds

**Supplementary Figure 10:** Ancestry painting for *A. marmorata* and *A. obscura*.
Ancestry painting as in Supplementary Figure 9, but for 73 "core" *A. marmorata* individuals, 26 "core" *A. obscura* individuals, and 3 recent hybrids between the two species. Horizontal bars indicate the genotypes at each of 742 sites fixed between the two parental species.

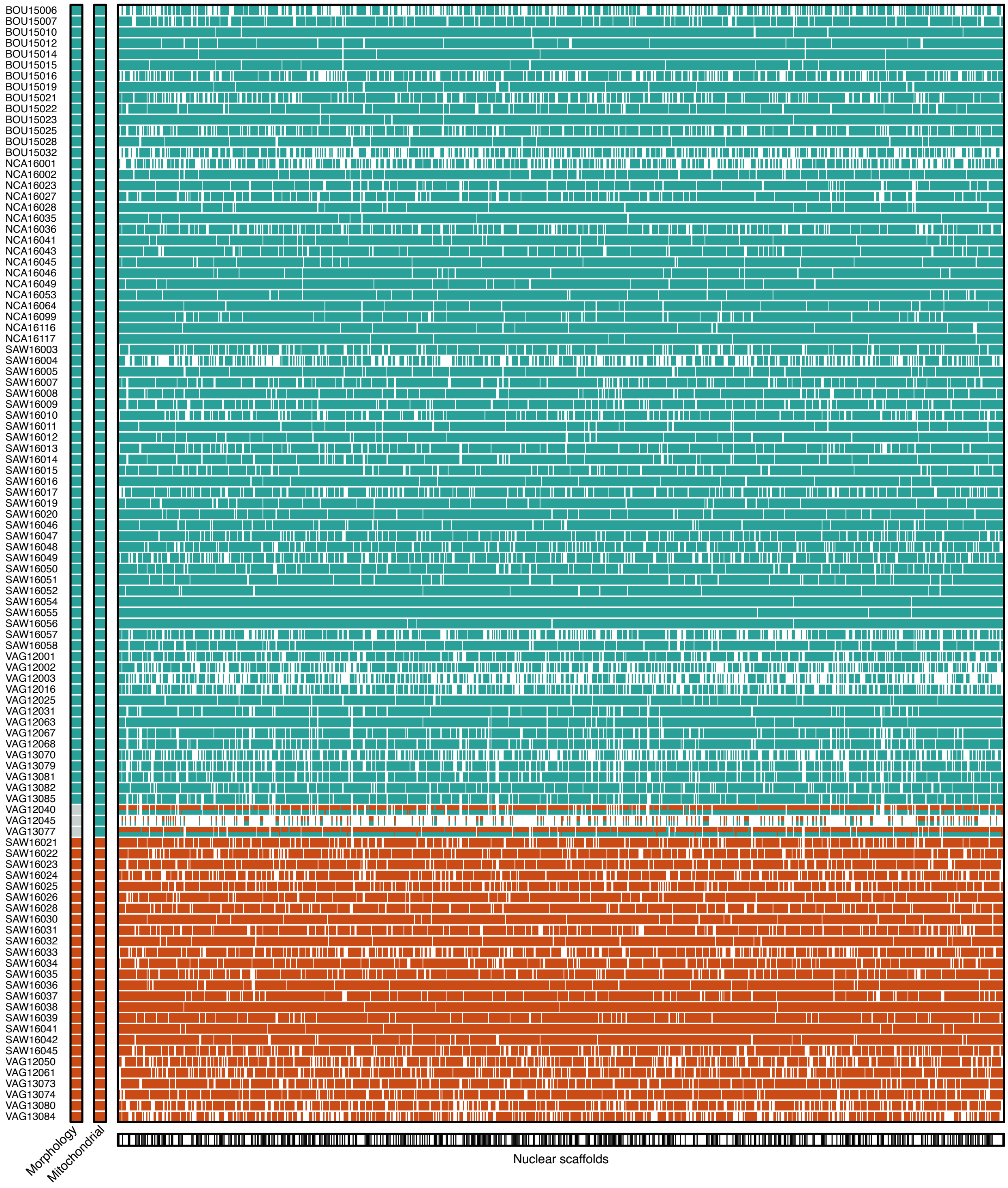

**Supplementary Figure 11:** Ancestry painting for *A. megastoma* and *A. obscura*.
Ancestry painting as in Supplementary Figure 9, but for 26 "core" *A. megastoma* individuals, 26 "core" *A. obscura* individuals, and 1 recent hybrid between the two species. Horizontal bars indicate the genotypes at each of 525 sites fixed between the two parental species.

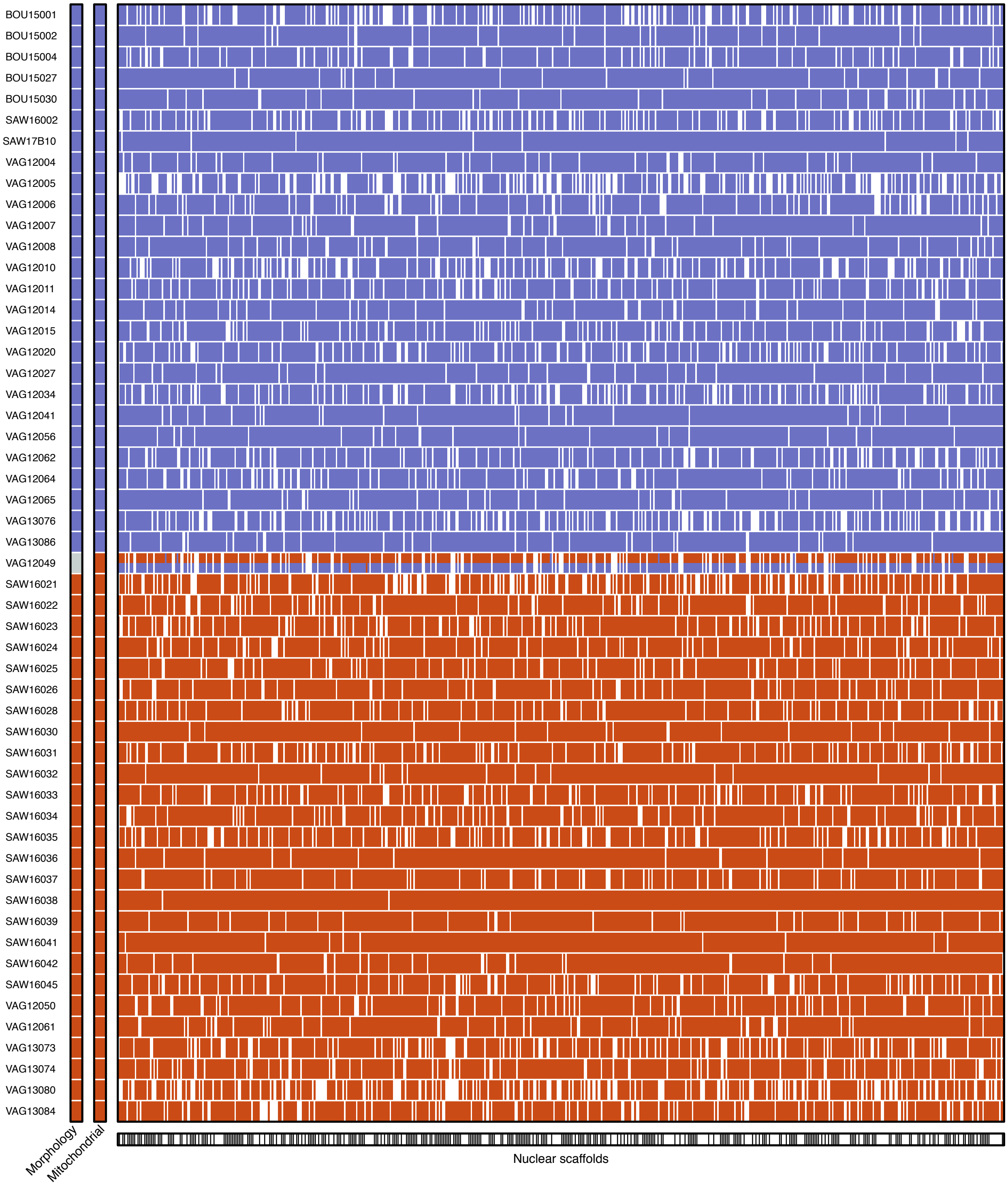

**Supplementary Figure 12:** Ancestry painting for *A. marmorata* and *A. interioris*.
Ancestry painting as in Supplementary Figure 9, but for 73 "core" *A. marmorata* individuals, 3 *A. interioris* individuals, and 1 recent hybrid between the two species. Horizontal bars indicate the genotypes at each of 429 sites fixed between the two parental species.

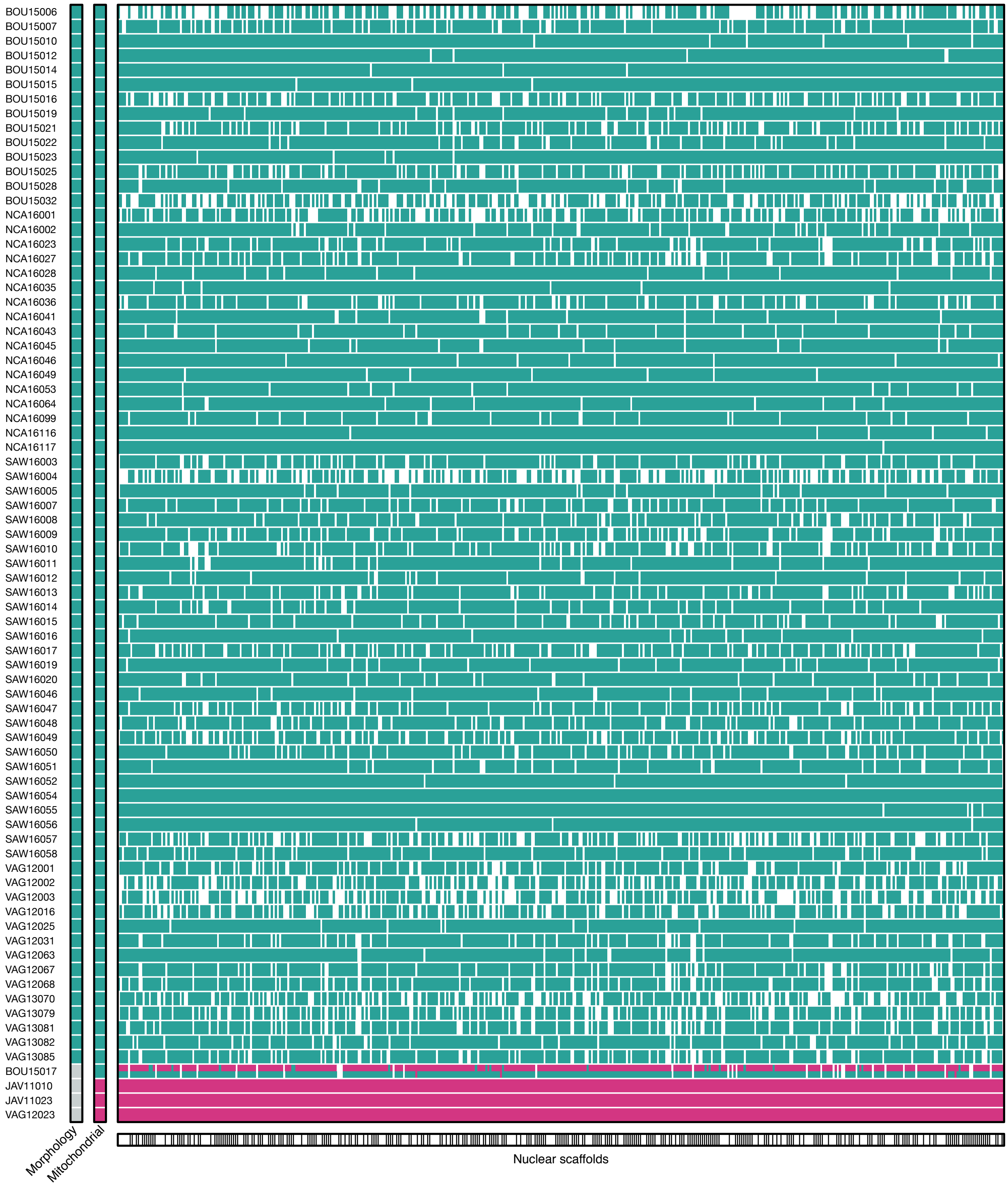

**Supplementary Figure 13:** Hybrid frequencies per sampling location.
Horizontal bars indicate the numbers of individuals collected at 14 sampling locations, counting only those with sufficient sequence quality that were used in genomic analyses. Hybrid frequencies are shown as black proportions of these bars.

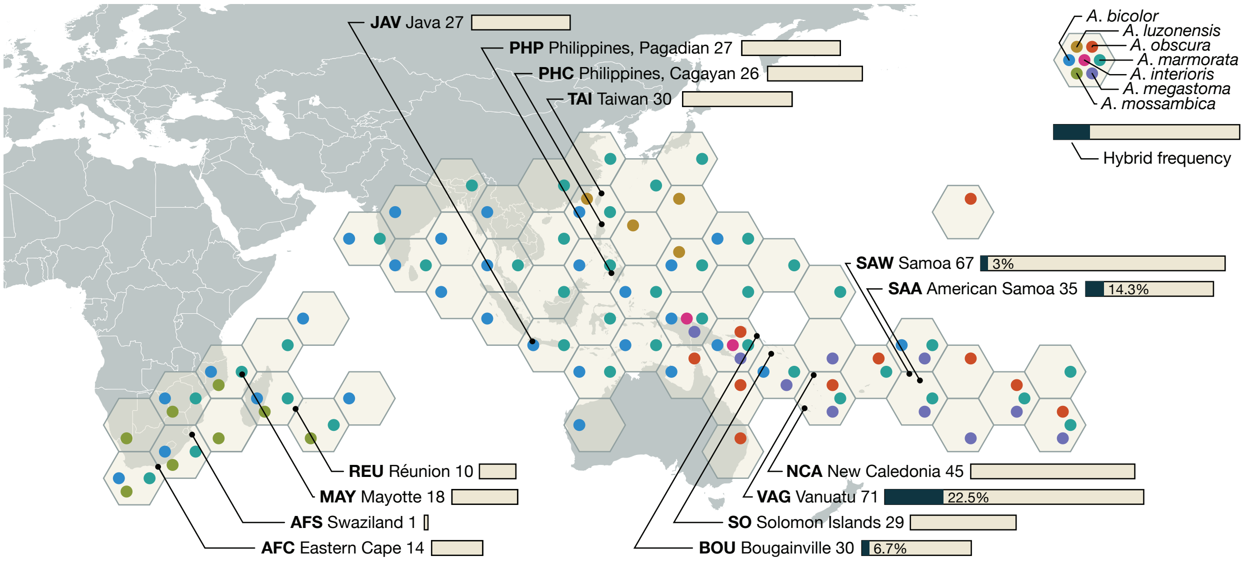

**Supplementary Figure 14:** Ancestry painting of long scaffolds for backcrossed hybrids between *A. marmorata* and *A. megastoma*.

Information shown here is a part of that presented in Supplementary Figure 9, focusing only on backcrossed hybrids and their genotypes on scaffolds with at least three fixed sites. Scaffold IDs are given on top and the positions of the first and last of the sites fixed on this scaffold are given below the ancestry painting. A single change from heterozygous to homozygous states or vice versa occurs six times among the seven individuals and two such changes on the same scaffold occur twice. With a mean distance of 1,708,024 bp and a maximum distance of 4,654,779 bp between the first and the last of the sites assessed on these 22 scaffolds, recombination breakpoints therefore appeared to be rare.

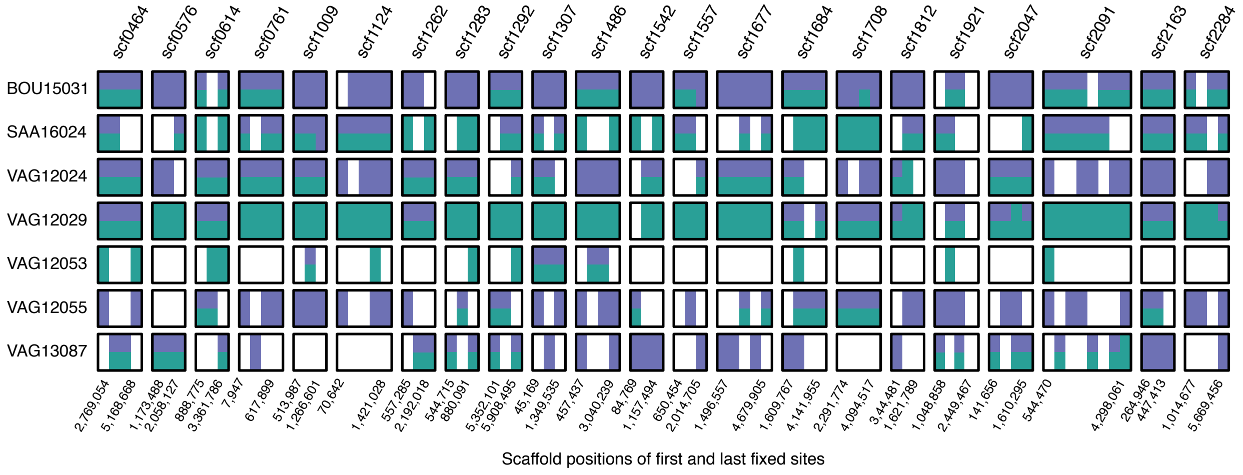

**Supplementary Figure 15:** Morphology of F1 and backcrossed hybrids between *A. marmorata* and *A. megastoma*.
Morphological measurements followed Watanabe *et al.* (2009). Labels indicate hybrids with transgressive phenotypes.

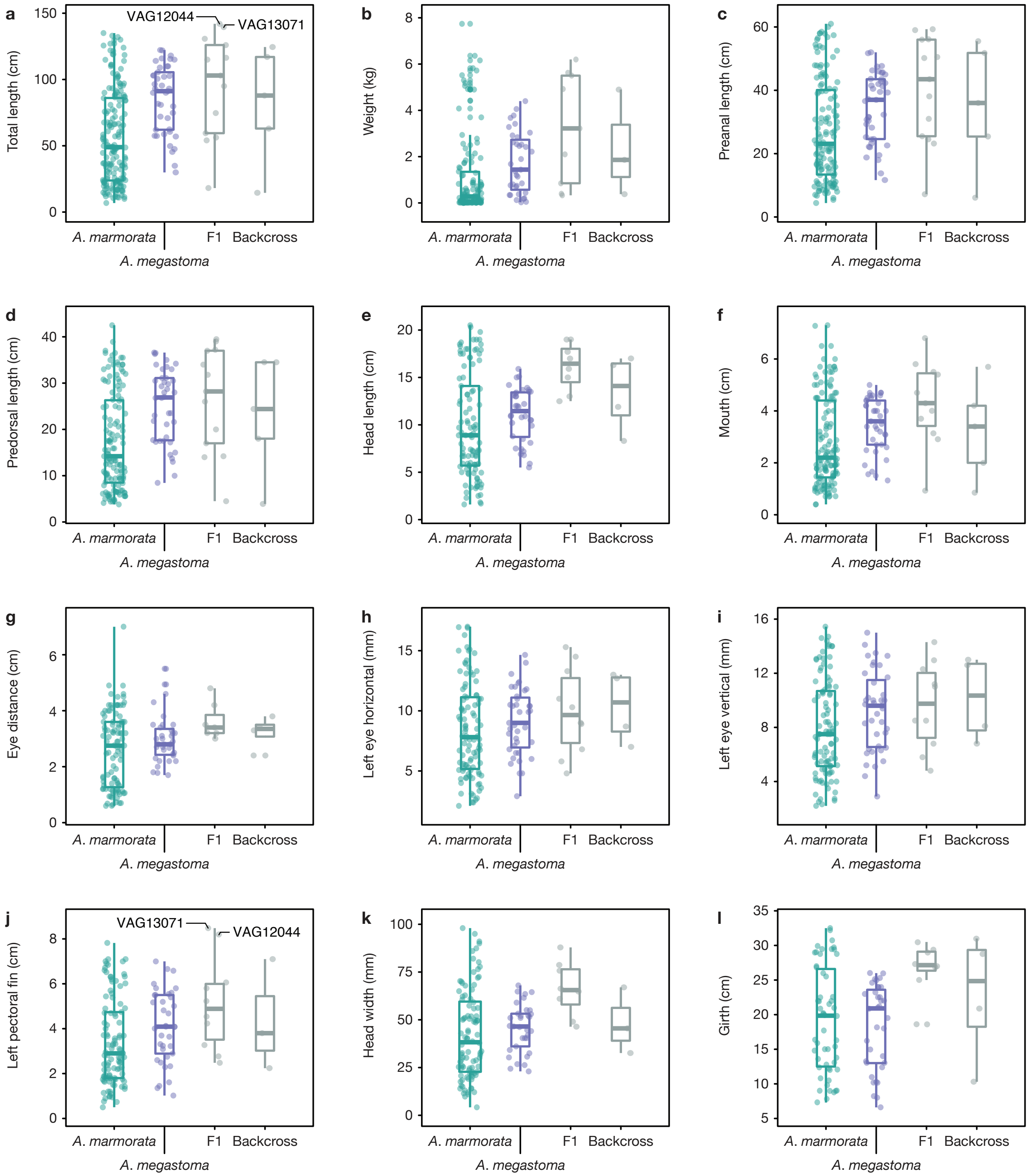

**Supplementary Figure 16:** Morphology of F1 hybrids between *A. marmorata* and *A. obscura*. Morphological measurements followed Watanabe *et al.* (2009).

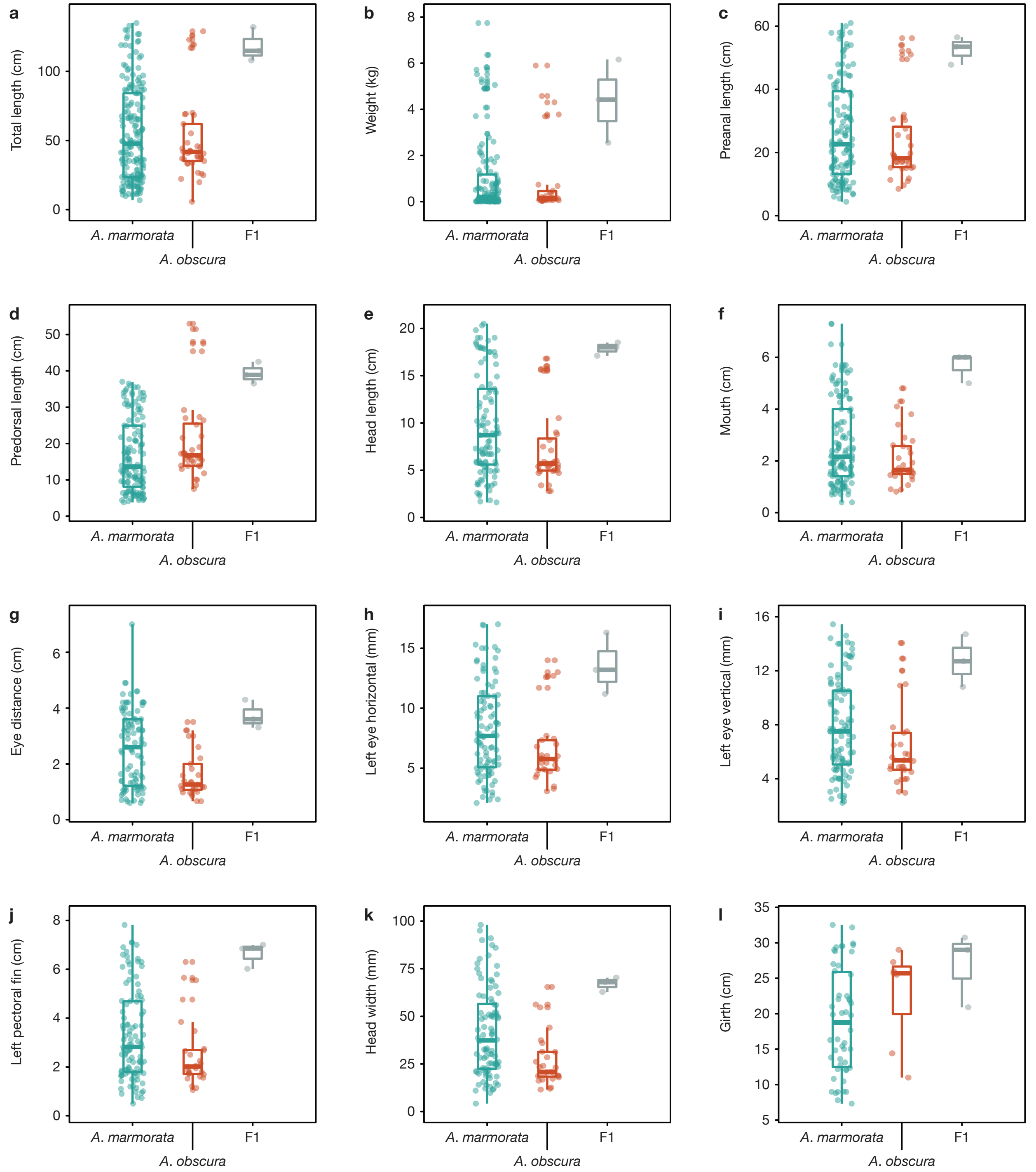

**Supplementary Figure 17:** Maximum-likelihood phylogenetic inference.
Phylogeny reconstructed with IQ-TREE (Nguyen *et al.* 2015) from 1,360 concatenated RAD loci without missing sequences and 20-40 variable sites per locus. Node labels indicate bootstrap support as well as per-locus (gCF) and per-site (sCF) concordance factors (Minh *et al.* 2018). For *A. marmorata*, *A. megastoma*, *A. luzonensis*, and *A. obscura*, only the five individuals with the lowest amount of missing data were used.

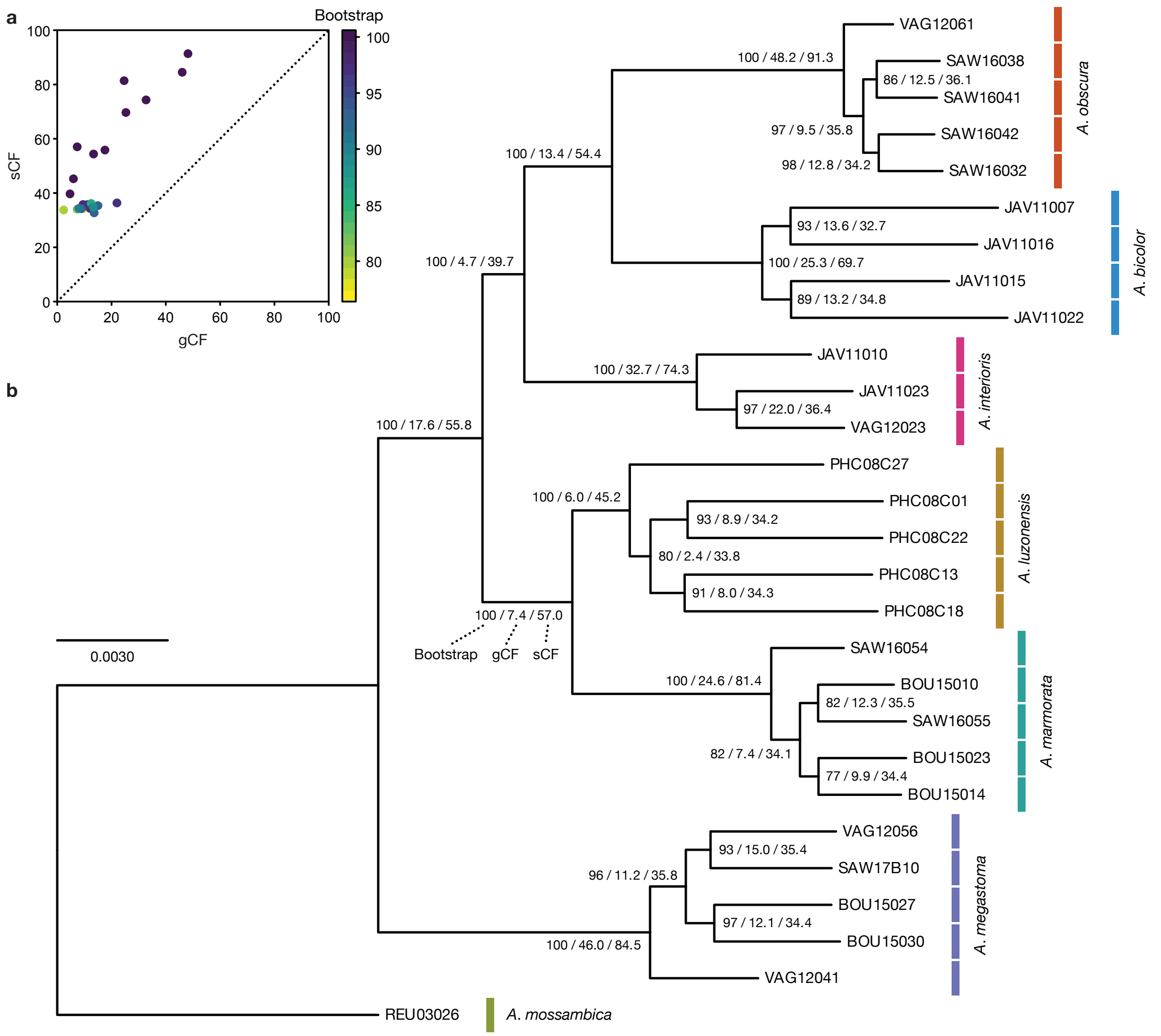

**Supplementary Figure 18:** Estimates of effective population size based on WGS data. Changes in effective population size ($N_e$; vertical axis) over time (the last 1 myr; horizontal axis) estimated using the pairwise sequential Markovian coalescent (PSMC). The PSMC was applied to WGS data of one individual for each of the three species *A. marmorata*, *A. megastoma*, and *A. obscura*. Estimates were based on assumed generation times ($g$) between 6 and 12 years and mutation rates ($\mu$) between 5.2 and 8.6 $\times 10^{-9}$ mutations/site/generation. Semi-transparent colored lines correspond to 100 bootstrap replicates. For visualization purposes, the range of contemporary $N_e$ values is truncated for *A. megastoma*; the maximum bootstrap value is $N_e = 2.0 \times 10^6$. Note that the apparent bottleneck pattern seen in all three species could be an artifact as it is expected even without actual population-size decline when parts of the genome are affected by introgression (Nielsen & Beaumont 2009).

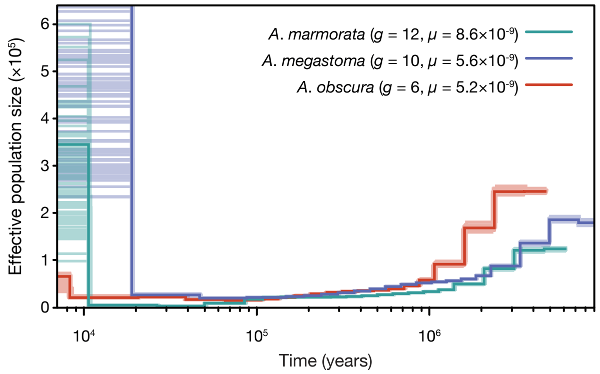

### 3 Supplementary Tables

**Supplementary Table 1:** Sampled specimens.
Underlined specimens were included in the "core" group of individuals for *A. marmorata*, *A. megastoma*, and *A. obscura*, based on morphological measurements characteristic for the species. Species assignment is based on mitochondrial sequence data, and additionally on morphological measurements when these were available. Solomon Islands sampling sites SOK (Kolombangara), SOL (Kolombangara), SON (Nggatokae), SOR (Ranongga), and SOV (Vangunu) are jointly labeled "SO" in Figure 1, Supplementary Figure 4a, and descriptions in the text. The availability of morphological information is indicated in the morphology column. Unless specified otherwise, all specimens for which morphology information was available were included in morphological principal component analysis. [1]specimen removed due to read number below 600,000; [2]specimen not included in morphological principal component analysis even though morphological information is available.

| Specimen ID | Species | Site | Lat. | Lon. | Date | # reads | Morphology |
|---|---|---|---|---|---|---|---|
| AFC09022 | *A. marmorata* | AFC | -33.615 | 25.667 | 2009/03/22 | 6,776,718 | partial |
| AFC09027 | *A. marmorata* | AFC | -33.615 | 25.667 | 2009/03/22 | 4,634,730 | partial |
| AFC09028 | *A. marmorata* | AFC | -33.615 | 25.667 | 2009/03/22 | 4,693,084 | partial |
| AFC09038 | *A. marmorata* | AFC | -33.615 | 25.667 | 2009/03/22 | 2,400,252 | partial |
| AFC09040 | *A. marmorata* | AFC | -33.615 | 25.667 | 2009/03/22 | 4,033,770 | partial |
| AFC09042 | *A. marmorata* | AFC | -33.615 | 25.667 | 2009/03/22 | 6,358,602 | partial |
| AFC09046 | *A. marmorata* | AFC | -33.615 | 25.667 | 2009/03/22 | 5,043,994 | partial |
| AFC09050 | *A. marmorata* | AFC | -33.615 | 25.667 | 2009/03/22 | 3,329,484 | partial |
| AFC09131 | *A. marmorata* | AFC | -33.046 | 26.662 | 2009/04/13 | 8,918,104 | partial |
| AFC09136 | *A. marmorata* | AFC | -33.046 | 26.662 | 2009/04/13 | 4,789,220 | partial |
| AFC09169 | *A. marmorata* | AFC | -33.615 | 25.667 | 2009/03/22 | 183,024[1] | partial |
| AFC09192 | *A. marmorata* | AFC | -33.615 | 25.667 | 2009/03/22 | 6,797,310 | partial |
| AFC09250 | *A. marmorata* | AFC | -33.615 | 25.667 | 2009/03/22 | 8,638,656 | partial |
| AFC09269 | *A. marmorata* | AFC | -33.615 | 25.667 | 2009/03/22 | 1,300,408 | partial |
| AFC09282 | *A. marmorata* | AFC | -33.615 | 25.667 | 2009/03/22 | 1,999,244 | partial |
| AFS03068 | *A. marmorata* | AFS | -26.713 | 31.979 | 2003/06/10 | 2,945,220 | no |
| BOU15001 | *A. megastoma* | BOU | -6.080 | 155.227 | 2015/04/04 | 10,524,168 | yes |
| BOU15002 | *A. megastoma* | BOU | -6.080 | 155.227 | 2015/04/04 | 8,753,268 | yes |
| BOU15003 | *A. megastoma* | BOU | -6.080 | 155.227 | 2015/04/04 | 12,765,510 | yes |
| BOU15004 | *A. megastoma* | BOU | -6.080 | 155.227 | 2015/04/04 | 8,763,996 | yes |
| BOU15005 | *A. megastoma* | BOU | -6.080 | 155.227 | 2015/04/05 | 6,346,666 | yes[2] |
| BOU15006 | *A. marmorata* | BOU | -5.982 | 155.365 | 2015/04/09 | 9,606,592 | yes |
| BOU15007 | *A. marmorata* | BOU | -5.982 | 155.365 | 2015/04/09 | 7,034,622 | yes |
| BOU15009 | *A. marmorata* | BOU | -5.982 | 155.365 | 2015/04/10 | 14,488,768 | yes |
| BOU15010 | *A. marmorata* | BOU | -5.982 | 155.365 | 2015/04/10 | 6,070,202 | yes |
| BOU15011 | *A. marmorata* | BOU | -5.982 | 155.365 | 2015/04/10 | 12,244,134 | yes |
| BOU15012 | *A. marmorata* | BOU | -5.982 | 155.365 | 2015/04/10 | 3,809,812 | yes |
| BOU15013 | *A. marmorata* | BOU | -5.982 | 155.365 | 2015/04/10 | 2,920,434 | yes[2] |
| BOU15014 | *A. marmorata* | BOU | -5.982 | 155.365 | 2015/04/10 | 4,069,002 | yes |
| BOU15015 | *A. marmorata* | BOU | -5.982 | 155.365 | 2015/04/10 | 4,142,280 | yes |
| BOU15016 | *A. marmorata* | BOU | -5.982 | 155.365 | 2015/04/10 | 8,406,776 | yes |

#### Supplementary Table 1 (continued)

| Specimen ID | Species | Site | Lat. | Lon. | Date | # reads | Morphology |
|---|---|---|---|---|---|---|---|
| BOU15017 | *A. marmorata* | BOU | -5.982 | 155.365 | 2015/04/11 | 6,477,100 | yes |
| BOU15018 | *A. marmorata* | BOU | -5.982 | 155.365 | 2015/04/12 | 16,346,782 | yes |
| BOU15019 | *A. marmorata* | BOU | -5.982 | 155.365 | 2015/04/12 | 3,438,714 | yes |
| BOU15020 | *A. marmorata* | BOU | -5.982 | 155.365 | 2015/04/12 | 4,981,164 | yes |
| BOU15021 | *A. marmorata* | BOU | -5.982 | 155.365 | 2015/04/12 | 1,490,284 | yes |
| BOU15022 | *A. marmorata* | BOU | -5.982 | 155.365 | 2015/04/13 | 2,212,684 | yes |
| BOU15023 | *A. marmorata* | BOU | -5.982 | 155.365 | 2015/04/13 | 4,800,140 | yes |
| BOU15024 | *A. marmorata* | BOU | -5.982 | 155.365 | 2015/04/12 | 12,316,926 | yes |
| BOU15025 | *A. marmorata* | BOU | -5.982 | 155.365 | 2015/04/14 | 1,569,478 | yes |
| BOU15027 | *A. megastoma* | BOU | -5.982 | 155.365 | 2015/04/14 | 4,104,044 | yes |
| BOU15028 | *A. marmorata* | BOU | -5.982 | 155.365 | 2015/04/14 | 3,353,376 | yes |
| BOU15029 | *A. marmorata* | BOU | -5.982 | 155.365 | 2015/04/14 | 715,048 | yes |
| BOU15030 | *A. megastoma* | BOU | -5.982 | 155.365 | 2015/04/14 | 2,705,922 | yes |
| BOU15031 | *A. marmorata* | BOU | -5.982 | 155.365 | 2015/04/14 | 4,074,020 | yes |
| BOU15032 | *A. marmorata* | BOU | -5.982 | 155.365 | 2015/04/12 | 8,950,896 | yes |
| JAV11001 | *A. marmorata* | JAV | -7.031 | 106.543 | 2011/06/NA | 1,911,324 | no |
| JAV11002 | *A. marmorata* | JAV | -7.031 | 106.543 | 2011/06/NA | 7,555,770 | no |
| JAV11003 | *A. marmorata* | JAV | -7.031 | 106.543 | 2011/06/NA | 1,090,534 | no |
| JAV11004 | *A. marmorata* | JAV | -7.031 | 106.543 | 2011/06/NA | 8,046,460 | no |
| JAV11005 | *A. marmorata* | JAV | -7.031 | 106.543 | 2011/06/NA | 2,310,702 | no |
| JAV11006 | *A. marmorata* | JAV | -7.031 | 106.543 | 2011/06/NA | 2,258,760 | no |
| JAV11007 | *A. bicolor* | JAV | -7.031 | 106.543 | 2011/06/NA | 2,688,928 | no |
| JAV11008 | *A. marmorata* | JAV | -7.031 | 106.543 | 2011/06/NA | 4,625,680 | no |
| JAV11009 | *A. marmorata* | JAV | -7.031 | 106.543 | 2011/06/NA | 1,036,900 | no |
| JAV11010 | *A. interioris* | JAV | -7.031 | 106.543 | 2011/06/NA | 6,425,584 | no |
| JAV11011 | *A. marmorata* | JAV | -7.031 | 106.543 | 2011/06/NA | 11,251,146 | no |
| JAV11012 | *A. marmorata* | JAV | -7.031 | 106.543 | 2011/06/NA | 6,491,164 | no |
| JAV11013 | *A. marmorata* | JAV | -7.031 | 106.543 | 2011/06/NA | 5,513,280 | no |
| JAV11014 | *A. marmorata* | JAV | -7.031 | 106.543 | 2011/06/NA | 9,067,812 | no |
| JAV11015 | *A. bicolor* | JAV | -7.031 | 106.543 | 2011/06/NA | 1,363,586 | no |
| JAV11016 | *A. bicolor* | JAV | -7.031 | 106.543 | 2011/06/NA | 2,046,036 | no |
| JAV11017 | *A. marmorata* | JAV | -7.031 | 106.543 | 2011/06/NA | 1,643,646 | no |
| JAV11018 | *A. marmorata* | JAV | -7.031 | 106.543 | 2011/06/NA | 7,677,132 | no |
| JAV11019 | *A. marmorata* | JAV | -7.031 | 106.543 | 2011/06/NA | 738,182 | no |
| JAV11020 | *A. marmorata* | JAV | -7.031 | 106.543 | 2011/06/NA | 272,336[1] | no |
| JAV11021 | *A. marmorata* | JAV | -7.031 | 106.543 | 2011/06/NA | 4,231,276 | no |
| JAV11022 | *A. bicolor* | JAV | -7.031 | 106.543 | 2011/06/NA | 11,179,098 | no |
| JAV11023 | *A. interioris* | JAV | -7.031 | 106.543 | 2011/06/NA | 5,903,968 | no |
| JAV11024 | *A. marmorata* | JAV | -7.031 | 106.543 | 2011/06/NA | 3,015,256 | no |
| JAV11025 | *A. marmorata* | JAV | -7.031 | 106.543 | 2011/06/NA | 1,471,420 | no |
| JAV11026 | *A. marmorata* | JAV | -7.031 | 106.543 | 2011/06/NA | 406,532[1] | no |
| JAV11027 | *A. marmorata* | JAV | -7.031 | 106.543 | 2011/06/NA | 1,378,514 | no |
| JAV11028 | *A. marmorata* | JAV | -7.031 | 106.543 | 2011/06/NA | 1,400,300 | no |
| JAV11029 | *A. marmorata* | JAV | -7.031 | 106.543 | 2011/06/NA | 374,992[1] | no |
| JAV11030 | *A. marmorata* | JAV | -7.031 | 106.543 | 2011/06/NA | 1,060,828 | no |
| MAY03001 | *A. marmorata* | MAY | -12.736 | 45.173 | 2003/11/09 | 3,635,428 | no |
| MAY03003 | *A. marmorata* | MAY | -12.736 | 45.173 | 2003/11/09 | 4,765,700 | no |

#### Supplementary Table 1 (continued)

| Specimen ID | Species | Site | Lat. | Lon. | Date | # reads | Morphology |
|---|---|---|---|---|---|---|---|
| MAY03005 | *A. marmorata* | MAY | -12.736 | 45.173 | 2003/11/09 | 4,158,434 | no |
| MAY03006 | *A. marmorata* | MAY | -12.736 | 45.173 | 2003/11/11 | 2,241,702 | no |
| MAY03007 | *A. marmorata* | MAY | -12.736 | 45.173 | 2003/11/09 | 1,725,756 | no |
| MAY03009 | *A. marmorata* | MAY | -12.736 | 45.173 | 2003/11/09 | 5,806,712 | no |
| MAY03013 | *A. marmorata* | MAY | -12.736 | 45.173 | 2003/11/09 | 2,181,048 | no |
| MAY03017 | *A. marmorata* | MAY | -12.736 | 45.173 | 2003/11/11 | 3,854,094 | no |
| MAY03018 | *A. marmorata* | MAY | -12.736 | 45.173 | 2003/11/09 | 2,476,808 | no |
| MAY03019 | *A. marmorata* | MAY | -12.736 | 45.173 | 2003/11/09 | 2,426,188 | no |
| MAY03020 | *A. marmorata* | MAY | -12.736 | 45.173 | 2003/11/09 | 1,422,048 | no |
| MAY03021 | *A. marmorata* | MAY | -12.736 | 45.173 | 2003/11/11 | 3,243,600 | no |
| MAY03022 | *A. marmorata* | MAY | -12.736 | 45.173 | 2003/11/11 | 4,022,620 | no |
| MAY03023 | *A. marmorata* | MAY | -12.736 | 45.173 | 2003/11/09 | 3,288,808 | no |
| MAY03024 | *A. marmorata* | MAY | -12.736 | 45.173 | 2003/11/11 | 3,669,560 | no |
| MAY03025 | *A. marmorata* | MAY | -12.736 | 45.173 | 2003/11/09 | 3,129,952 | no |
| MAY03027 | *A. marmorata* | MAY | -12.736 | 45.173 | 2003/11/11 | 2,452,074 | no |
| MAY03028 | *A. marmorata* | MAY | -12.736 | 45.173 | 2003/11/11 | 2,337,216 | no |
| NCA16001 | *A. marmorata* | NCA | -21.305 | 165.025 | 2016/07/27 | 1,463,126 | yes |
| NCA16002 | *A. marmorata* | NCA | -21.305 | 165.025 | 2016/07/27 | 3,965,922 | yes |
| NCA16003 | *A. marmorata* | NCA | -21.305 | 165.025 | 2016/07/27 | 885,746 | yes |
| NCA16004 | *A. marmorata* | NCA | -21.305 | 165.025 | 2016/07/27 | 3,202,396 | partial |
| NCA16005 | *A. marmorata* | NCA | -21.305 | 165.025 | 2016/07/27 | 3,506,424 | partial |
| NCA16006 | *A. marmorata* | NCA | -21.305 | 165.025 | 2016/07/27 | 6,456,506 | partial |
| NCA16007 | *A. marmorata* | NCA | -21.305 | 165.025 | 2016/07/27 | 4,056,230 | partial |
| NCA16008 | *A. marmorata* | NCA | -21.305 | 165.025 | 2016/07/27 | 2,850,918 | partial |
| NCA16009 | *A. marmorata* | NCA | -21.305 | 165.025 | 2016/07/27 | 4,937,224 | partial |
| NCA16010 | *A. marmorata* | NCA | -21.305 | 165.025 | 2016/07/27 | 4,202,782 | no |
| NCA16011 | *A. marmorata* | NCA | -21.305 | 165.025 | 2016/07/27 | 4,716,276 | no |
| NCA16014 | *A. marmorata* | NCA | -21.305 | 165.025 | 2016/07/27 | 4,179,094 | partial |
| NCA16015 | *A. obscura* | NCA | -21.305 | 165.025 | 2016/07/27 | 1,913,782 | partial |
| NCA16018 | *A. marmorata* | NCA | -21.302 | 165.029 | 2016/07/28 | 7,270,454 | partial |
| NCA16020 | *A. marmorata* | NCA | -21.302 | 165.029 | 2016/07/28 | 6,578,652 | partial |
| NCA16021 | *A. marmorata* | NCA | -21.302 | 165.029 | 2016/07/28 | 3,054,210 | partial |
| NCA16022 | *A. marmorata* | NCA | -21.302 | 165.029 | 2016/07/28 | 2,739,816 | partial |
| NCA16023 | *A. marmorata* | NCA | -21.302 | 165.029 | 2016/07/28 | 6,474,304 | yes |
| NCA16024 | *A. marmorata* | NCA | -21.302 | 165.029 | 2016/07/28 | 6,085,708 | partial |
| NCA16025 | *A. marmorata* | NCA | -21.302 | 165.029 | 2016/07/28 | 1,234,354 | partial |
| NCA16027 | *A. marmorata* | NCA | -21.302 | 165.029 | 2016/07/28 | 7,005,388 | yes |
| NCA16028 | *A. marmorata* | NCA | -21.302 | 165.029 | 2016/07/28 | 5,545,016 | yes |
| NCA16030 | *A. marmorata* | NCA | -21.302 | 165.029 | 2016/07/28 | 4,546,024 | partial |
| NCA16031 | *A. marmorata* | NCA | -20.491 | 164.258 | 2016/08/02 | 4,881,226 | partial |
| NCA16034 | *A. marmorata* | NCA | -20.491 | 164.258 | 2016/08/02 | 1,596,114 | partial |
| NCA16035 | *A. marmorata* | NCA | -20.491 | 164.258 | 2016/08/02 | 4,372,818 | yes |
| NCA16036 | *A. marmorata* | NCA | -20.491 | 164.258 | 2016/08/02 | 1,913,554 | yes |
| NCA16039 | *A. marmorata* | NCA | -20.491 | 164.258 | 2016/08/02 | 1,995,702 | partial |
| NCA16041 | *A. marmorata* | NCA | -20.491 | 164.258 | 2016/08/02 | 3,543,860 | yes |
| NCA16042 | *A. marmorata* | NCA | -20.491 | 164.258 | 2016/08/02 | 5,045,656 | partial |
| NCA16043 | *A. marmorata* | NCA | -20.491 | 164.258 | 2016/08/02 | 2,637,300 | yes |

#### Supplementary Table 1 (continued)

| Specimen ID | Species | Site | Lat. | Lon. | Date | # reads | Morphology |
|---|---|---|---|---|---|---|---|
| NCA16044 | *A. marmorata* | NCA | -20.491 | 164.258 | 2016/08/02 | 5,381,242 | partial |
| NCA16045 | *A. marmorata* | NCA | -20.491 | 164.258 | 2016/08/02 | 3,630,832 | yes |
| NCA16046 | *A. marmorata* | NCA | -22.111 | 166.423 | 2016/08/09 | 4,248,040 | yes |
| NCA16049 | *A. marmorata* | NCA | -22.111 | 166.423 | 2016/08/09 | 4,212,004 | yes |
| NCA16053 | *A. marmorata* | NCA | -22.136 | 166.367 | 2016/08/09 | 3,653,266 | yes |
| NCA16056 | *A. marmorata* | NCA | -22.136 | 166.367 | 2016/08/09 | 6,829,734 | partial |
| NCA16063 | *A. marmorata* | NCA | -22.136 | 166.367 | 2016/08/09 | 931,716 | yes[2] |
| NCA16064 | *A. marmorata* | NCA | -22.136 | 166.367 | 2016/08/09 | 4,739,518 | yes |
| NCA16084 | *A. marmorata* | NCA | -21.749 | 166.084 | 2016/08/11 | 5,874,506 | yes |
| NCA16099 | *A. marmorata* | NCA | -21.749 | 166.084 | 2016/08/11 | 7,063,454 | yes |
| NCA16110 | *A. marmorata* | NCA | -22.035 | 166.208 | 2016/08/18 | 4,691,496 | partial |
| NCA16116 | *A. marmorata* | NCA | -22.035 | 166.208 | 2016/08/18 | 4,702,550 | yes |
| NCA16117 | *A. marmorata* | NCA | -22.035 | 166.208 | 2016/08/18 | 4,790,108 | yes |
| NCA16120 | *A. marmorata* | NCA | -22.038 | 166.220 | 2016/08/18 | 6,610,270 | partial |
| PHC08C01 | *A. luzonensis* | PHC | 18.355 | 121.634 | 2008/09/08 | 4,385,110 | no |
| PHC08C02 | *A. luzonensis* | PHC | 18.355 | 121.634 | 2008/09/08 | 2,067,082 | no |
| PHC08C03 | *A. luzonensis* | PHC | 18.355 | 121.634 | 2008/09/08 | 6,693,386 | no |
| PHC08C04 | *A. luzonensis* | PHC | 18.355 | 121.634 | 2008/09/08 | 7,107,044 | no |
| PHC08C05 | *A. luzonensis* | PHC | 18.355 | 121.634 | 2008/09/08 | 8,824,514 | no |
| PHC08C06 | *A. luzonensis* | PHC | 18.355 | 121.634 | 2008/09/08 | 9,469,068 | no |
| PHC08C07 | *A. luzonensis* | PHC | 18.355 | 121.634 | 2008/09/08 | 6,707,486 | no |
| PHC08C08 | *A. luzonensis* | PHC | 18.355 | 121.634 | 2008/09/08 | 176,638[1] | no |
| PHC08C09 | *A. luzonensis* | PHC | 18.355 | 121.634 | 2008/09/08 | 6,465,580 | no |
| PHC08C10 | *A. marmorata* | PHC | 18.355 | 121.634 | 2008/09/08 | 6,767,542 | no |
| PHC08C11 | *A. luzonensis* | PHC | 18.355 | 121.634 | 2008/09/08 | 11,154,032 | no |
| PHC08C12 | *A. luzonensis* | PHC | 18.355 | 121.634 | 2008/09/08 | 6,774,270 | no |
| PHC08C13 | *A. luzonensis* | PHC | 18.355 | 121.634 | 2008/09/08 | 5,412,554 | no |
| PHC08C15 | *A. luzonensis* | PHC | 18.355 | 121.634 | 2008/09/08 | 3,325,444 | no |
| PHC08C16 | *A. luzonensis* | PHC | 18.355 | 121.634 | 2008/09/08 | 3,157,590 | no |
| PHC08C17 | *A. luzonensis* | PHC | 18.355 | 121.634 | 2008/09/08 | 35,658[1] | no |
| PHC08C18 | *A. luzonensis* | PHC | 18.355 | 121.634 | 2008/09/08 | 5,386,382 | no |
| PHC08C19 | *A. marmorata* | PHC | 18.355 | 121.634 | 2008/09/08 | 301,666[1] | no |
| PHC08C20 | *A. marmorata* | PHC | 18.355 | 121.634 | 2008/09/08 | 4,533,308 | no |
| PHC08C21 | *A. marmorata* | PHC | 18.355 | 121.634 | 2008/09/08 | 50,532[1] | no |
| PHC08C22 | *A. luzonensis* | PHC | 18.355 | 121.634 | 2008/09/08 | 6,089,432 | no |
| PHC08C23 | *A. luzonensis* | PHC | 18.355 | 121.634 | 2008/09/08 | 58,386[1] | no |
| PHC08C24 | *A. luzonensis* | PHC | 18.355 | 121.634 | 2008/09/08 | 778,644 | no |
| PHC08C25 | *A. marmorata* | PHC | 18.355 | 121.634 | 2008/09/08 | 3,036,944 | no |
| PHC08C26 | *A. luzonensis* | PHC | 18.355 | 121.634 | 2008/09/08 | 8,041,488 | no |
| PHC08C27 | *A. luzonensis* | PHC | 18.355 | 121.634 | 2008/09/08 | 3,149,128 | no |
| PHC08C28 | *A. luzonensis* | PHC | 18.355 | 121.634 | 2008/09/08 | 1,770,520 | no |
| PHC08C29 | *A. luzonensis* | PHC | 18.355 | 121.634 | 2008/09/08 | 1,787,774 | no |
| PHC08P20 | *A. marmorata* | PHC | 18.355 | 121.634 | 2008/09/08 | 3,961,354 | no |
| PHC08P22 | *A. marmorata* | PHC | 18.355 | 121.634 | 2008/09/08 | 4,094,926 | no |
| PHC08P23 | *A. marmorata* | PHC | 18.355 | 121.634 | 2008/09/08 | 5,894,520 | no |
| PHP14P01 | *A. marmorata* | PHP | 7.835 | 123.509 | 2014/02/14 | 4,783,294 | no |
| PHP14P02 | *A. marmorata* | PHP | 7.835 | 123.509 | 2014/02/14 | 5,889,742 | no |

#### Supplementary Table 1 (continued)

| Specimen ID | Species | Site | Lat. | Lon. | Date | # reads | Morphology |
|---|---|---|---|---|---|---|---|
| PHP14P03 | *A. marmorata* | PHP | 7.835 | 123.509 | 2014/02/14 | 6,371,944 | no |
| PHP14P04 | *A. marmorata* | PHP | 7.835 | 123.509 | 2014/02/14 | 4,141,222 | no |
| PHP14P05 | *A. marmorata* | PHP | 7.835 | 123.509 | 2014/02/14 | 5,544,038 | no |
| PHP14P06 | *A. marmorata* | PHP | 7.835 | 123.509 | 2014/02/14 | 2,983,648 | no |
| PHP14P07 | *A. marmorata* | PHP | 7.835 | 123.509 | 2014/02/14 | 4,178,472 | no |
| PHP14P08 | *A. marmorata* | PHP | 7.835 | 123.509 | 2014/02/14 | 3,034,380 | no |
| PHP14P09 | *A. marmorata* | PHP | 7.835 | 123.509 | 2014/02/14 | 4,966,488 | no |
| PHP14P10 | *A. marmorata* | PHP | 7.835 | 123.509 | 2014/02/14 | 5,286,256 | no |
| PHP14P11 | *A. marmorata* | PHP | 7.835 | 123.509 | 2014/02/14 | 6,830,854 | no |
| PHP14P12 | *A. marmorata* | PHP | 7.835 | 123.509 | 2014/02/14 | 3,904,972 | no |
| PHP14P13 | *A. marmorata* | PHP | 7.835 | 123.509 | 2014/02/14 | 3,827,650 | no |
| PHP14P14 | *A. marmorata* | PHP | 7.835 | 123.509 | 2014/02/14 | 5,505,830 | no |
| PHP14P15 | *A. marmorata* | PHP | 7.835 | 123.509 | 2014/02/14 | 2,567,042 | no |
| PHP14P16 | *A. marmorata* | PHP | 7.835 | 123.509 | 2014/02/14 | 770,766 | no |
| PHP14P17 | *A. marmorata* | PHP | 7.835 | 123.509 | 2014/02/14 | 4,868,568 | no |
| PHP14P18 | *A. marmorata* | PHP | 7.835 | 123.509 | 2014/02/14 | 4,753,592 | no |
| PHP14P19 | *A. marmorata* | PHP | 7.835 | 123.509 | 2014/02/14 | 4,047,700 | no |
| PHP14P21 | *A. marmorata* | PHP | 7.835 | 123.509 | 2014/02/14 | 1,969,502 | no |
| PHP14P24 | *A. marmorata* | PHP | 7.835 | 123.509 | 2014/02/14 | 6,280,804 | no |
| PHP14P25 | *A. marmorata* | PHP | 7.835 | 123.509 | 2014/02/14 | 6,543,116 | no |
| PHP14P26 | *A. marmorata* | PHP | 7.835 | 123.509 | 2014/02/14 | 7,983,072 | no |
| PHP14P27 | *A. marmorata* | PHP | 7.835 | 123.509 | 2014/02/14 | 9,373,838 | no |
| PHP14P28 | *A. marmorata* | PHP | 7.835 | 123.509 | 2014/02/14 | 8,095,200 | no |
| PHP14P29 | *A. marmorata* | PHP | 7.835 | 123.509 | 2014/02/14 | 6,411,040 | no |
| PHP14P30 | *A. marmorata* | PHP | 7.835 | 123.509 | 2014/02/14 | 6,390,714 | no |
| REU01002 | *A. marmorata* | REU | -20.983 | 55.685 | 2001/02/05 | 3,454,510 | no |
| REU01014 | *A. marmorata* | REU | -20.983 | 55.685 | 2001/02/05 | 2,737,978 | no |
| REU01016 | *A. marmorata* | REU | -20.983 | 55.685 | 2001/02/05 | 1,923,798 | no |
| REU03004 | *A. marmorata* | REU | -20.983 | 55.685 | 2003/11/04 | 3,841,652 | no |
| REU03008 | *A. marmorata* | REU | -20.983 | 55.685 | 2003/11/04 | 3,107,032 | no |
| REU03010 | *A. marmorata* | REU | -20.912 | 55.630 | 2003/11/04 | 4,676,038 | no |
| REU03011 | *A. marmorata* | REU | -20.983 | 55.685 | 2003/11/04 | 4,215,150 | no |
| REU03012 | *A. marmorata* | REU | -20.912 | 55.630 | 2003/11/04 | 8,125,876 | no |
| REU03015 | *A. marmorata* | REU | -20.912 | 55.630 | 2003/11/04 | 3,492,136 | no |
| REU03026 | *A. mossambica* | REU | -20.912 | 55.630 | 2003/11/04 | 3,450,942 | no |
| SAA16001 | *A. marmorata* | SAA | -14.304 | 170.816 | 2016/08/18 | 7,116,822 | partial |
| SAA16002 | *A. marmorata* | SAA | -14.304 | 170.816 | 2016/08/19 | 7,015,912 | partial |
| SAA16003 | *A. marmorata* | SAA | -14.304 | 170.816 | 2016/08/18 | 3,522,172 | partial |
| SAA16004 | *A. marmorata* | SAA | -14.304 | 170.816 | 2016/08/18 | 3,740,088 | partial |
| SAA16005 | *A. marmorata* | SAA | -14.304 | 170.816 | 2016/08/18 | 18,305,104 | no |
| SAA16006 | *A. marmorata* | SAA | -14.304 | 170.816 | 2016/08/18 | 6,540,862 | partial |
| SAA16007 | *A. marmorata* | SAA | -14.304 | 170.816 | 2016/08/18 | 3,685,986 | partial |
| SAA16008 | *A. marmorata* | SAA | -14.304 | 170.816 | 2016/08/18 | 5,828,106 | partial |
| SAA16009 | *A. marmorata* | SAA | -14.304 | 170.816 | 2016/08/18 | 5,979,728 | partial |
| SAA16010 | *A. marmorata* | SAA | -14.304 | 170.816 | 2016/08/18 | 1,269,150 | partial |
| SAA16011 | *A. marmorata* | SAA | -14.304 | 170.816 | 2016/08/18 | 3,557,730 | partial |
| SAA16012 | *A. marmorata* | SAA | -14.304 | 170.816 | 2016/08/18 | 7,436,372 | partial |

#### Supplementary Table 1 (continued)

| Specimen ID | Species | Site | Lat. | Lon. | Date | # reads | Morphology |
|---|---|---|---|---|---|---|---|
| SAA16013 | *A. marmorata* | SAA | -14.304 | 170.816 | 2016/08/18 | 2,752,316 | partial |
| SAA16014 | *A. megastoma* | SAA | -14.332 | 170.793 | 2016/08/19 | 3,018,560 | partial |
| SAA16015 | *A. megastoma* | SAA | -14.332 | 170.793 | 2016/08/19 | 5,022,542 | partial |
| SAA16016 | *A. marmorata* | SAA | -14.332 | 170.793 | 2016/08/19 | 7,564,634 | partial |
| SAA16017 | *A. marmorata* | SAA | -14.332 | 170.793 | 2016/08/19 | 881,890 | partial |
| SAA16018 | *A. marmorata* | SAA | -14.332 | 170.793 | 2016/08/19 | 6,003,128 | partial |
| SAA16019 | *A. marmorata* | SAA | -14.332 | 170.793 | 2016/08/19 | 1,456,892 | partial |
| SAA16020 | *A. marmorata* | SAA | -14.332 | 170.793 | 2016/08/19 | 4,554,650 | partial |
| SAA16021 | *A. marmorata* | SAA | -14.332 | 170.793 | 2016/08/19 | 1,433,016 | partial |
| SAA16022 | *A. marmorata* | SAA | -14.332 | 170.793 | 2016/08/19 | 5,381,110 | partial |
| SAA16023 | *A. marmorata* | SAA | -14.304 | 170.816 | 2016/08/20 | 5,759,626 | partial |
| SAA16024 | *A. marmorata* | SAA | -14.304 | 170.816 | 2016/08/20 | 7,833,688 | partial |
| SAA16025 | *A. marmorata* | SAA | -14.304 | 170.816 | 2016/08/20 | 3,393,344 | partial |
| SAA16026 | *A. marmorata* | SAA | -14.304 | 170.816 | 2016/08/20 | 6,973,470 | partial |
| SAA16027 | *A. marmorata* | SAA | -14.304 | 170.816 | 2016/08/20 | 8,727,666 | partial |
| SAA16028 | *A. marmorata* | SAA | -14.304 | 170.816 | 2016/08/20 | 12,240[1] | partial |
| SAA16029 | *A. marmorata* | SAA | -14.304 | 170.816 | 2016/08/20 | 9,310,208 | partial |
| SAA16030 | *A. marmorata* | SAA | -14.304 | 170.816 | 2016/08/20 | 4,058,900 | partial |
| SAA16031 | *A. marmorata* | SAA | -14.304 | 170.816 | 2016/08/20 | 4,624,388 | partial |
| SAA16032 | *A. marmorata* | SAA | -14.304 | 170.816 | 2016/08/20 | 5,149,616 | partial |
| SAA16033 | *A. marmorata* | SAA | -14.304 | 170.816 | 2016/08/20 | 266,538[1] | partial |
| SAA16034 | *A. marmorata* | SAA | -14.304 | 170.816 | 2016/08/20 | 7,292,490 | partial |
| SAA16035 | *A. marmorata* | SAA | -14.304 | 170.816 | 2016/08/20 | 6,989,336 | partial |
| SAA16036 | *A. marmorata* | SAA | -14.304 | 170.816 | 2016/08/20 | 7,190,652 | partial |
| SAA16037 | *A. marmorata* | SAA | -14.304 | 170.816 | 2016/08/20 | 58,520[1] | partial |
| SAA16038 | *A. marmorata* | SAA | -14.304 | 170.816 | 2016/08/20 | 3,566,502 | partial |
| SAW16001 | *A. marmorata* | SAW | -13.836 | 171.765 | 2016/08/16 | 2,750,534 | yes |
| SAW16002 | *A. megastoma* | SAW | -13.904 | 171.575 | 2016/08/16 | 8,122,668 | yes |
| SAW16003 | *A. marmorata* | SAW | -13.874 | 171.651 | 2016/08/31 | 7,612,856 | yes |
| SAW16004 | *A. marmorata* | SAW | -13.874 | 171.651 | 2016/08/31 | 1,578,510 | yes |
| SAW16005 | *A. marmorata* | SAW | -13.874 | 171.651 | 2016/08/31 | 5,100,238 | yes |
| SAW16006 | *A. marmorata* | SAW | -13.874 | 171.651 | 2016/08/31 | 47,706[1] | yes[2] |
| SAW16007 | *A. marmorata* | SAW | -13.874 | 171.651 | 2016/08/31 | 2,616,202 | yes |
| SAW16008 | *A. marmorata* | SAW | -13.874 | 171.651 | 2016/08/31 | 3,802,620 | yes |
| SAW16009 | *A. marmorata* | SAW | -13.874 | 171.651 | 2016/08/31 | 2,947,904 | yes |
| SAW16010 | *A. marmorata* | SAW | -13.874 | 171.651 | 2016/08/31 | 8,468,370 | yes |
| SAW16011 | *A. marmorata* | SAW | -13.874 | 171.651 | 2016/08/31 | 5,449,034 | yes |
| SAW16012 | *A. marmorata* | SAW | -13.874 | 171.651 | 2016/08/31 | 4,835,336 | yes |
| SAW16013 | *A. marmorata* | SAW | -13.874 | 171.651 | 2016/08/31 | 7,091,060 | yes |
| SAW16014 | *A. marmorata* | SAW | -13.874 | 171.651 | 2016/08/31 | 4,780,360 | yes |
| SAW16015 | *A. marmorata* | SAW | -13.874 | 171.651 | 2016/08/31 | 3,078,620 | yes |
| SAW16016 | *A. marmorata* | SAW | -13.874 | 171.651 | 2016/08/31 | 4,916,950 | yes |
| SAW16017 | *A. marmorata* | SAW | -13.874 | 171.651 | 2016/08/31 | 7,284,612 | yes |
| SAW16018 | *A. marmorata* | SAW | -13.874 | 171.651 | 2016/08/31 | 25,368[1] | yes[2] |
| SAW16019 | *A. marmorata* | SAW | -13.874 | 171.651 | 2016/08/31 | 3,451,030 | yes |
| SAW16020 | *A. marmorata* | SAW | -13.874 | 171.651 | 2016/08/31 | 3,764,650 | yes |
| SAW16021 | *A. obscura* | SAW | -14.025 | 171.431 | 2016/09/01 | 8,156,690 | yes |

#### Supplementary Table 1 (continued)

| Specimen ID | Species | Site | Lat. | Lon. | Date | # reads | Morphology |
|---|---|---|---|---|---|---|---|
| SAW16022 | *A. obscura* | SAW | -14.025 | 171.431 | 2016/09/01 | 3,012,538 | yes |
| SAW16023 | *A. obscura* | SAW | -14.025 | 171.431 | 2016/09/01 | 4,522,382 | yes |
| SAW16024 | *A. obscura* | SAW | -14.025 | 171.431 | 2016/09/01 | 3,250,488 | yes |
| SAW16025 | *A. obscura* | SAW | -14.025 | 171.431 | 2016/09/01 | 2,821,526 | yes |
| SAW16026 | *A. obscura* | SAW | -14.025 | 171.431 | 2016/09/01 | 3,173,878 | yes |
| SAW16027 | *A. obscura* | SAW | -14.025 | 171.431 | 2016/09/01 | 2,975,754 | yes[2] |
| SAW16028 | *A. obscura* | SAW | -14.025 | 171.431 | 2016/09/01 | 2,657,622 | yes |
| SAW16029 | *A. obscura* | SAW | -14.025 | 171.431 | 2016/09/01 | 399,402[1] | yes[2] |
| SAW16030 | *A. obscura* | SAW | -14.025 | 171.431 | 2016/09/01 | 6,092,366 | yes |
| SAW16031 | *A. obscura* | SAW | -14.025 | 171.431 | 2016/09/01 | 2,508,612 | yes |
| SAW16032 | *A. obscura* | SAW | -14.025 | 171.431 | 2016/09/01 | 4,808,164 | yes |
| SAW16033 | *A. obscura* | SAW | -14.025 | 171.431 | 2016/09/01 | 6,932,430 | yes |
| SAW16034 | *A. obscura* | SAW | -14.025 | 171.431 | 2016/09/01 | 2,710,582 | yes |
| SAW16035 | *A. obscura* | SAW | -14.025 | 171.431 | 2016/09/01 | 7,565,924 | yes |
| SAW16036 | *A. obscura* | SAW | -14.025 | 171.431 | 2016/09/01 | 6,343,662 | yes |
| SAW16037 | *A. obscura* | SAW | -14.025 | 171.431 | 2016/09/01 | 7,089,804 | yes |
| SAW16038 | *A. obscura* | SAW | -14.025 | 171.431 | 2016/09/01 | 5,230,320 | yes |
| SAW16039 | *A. obscura* | SAW | -14.025 | 171.431 | 2016/09/01 | 4,170,504 | yes |
| SAW16040 | *A. obscura* | SAW | -14.025 | 171.431 | 2016/09/01 | 9,956,058 | yes[2] |
| SAW16041 | *A. obscura* | SAW | -14.025 | 171.431 | 2016/09/01 | 5,220,798 | yes |
| SAW16042 | *A. obscura* | SAW | -14.025 | 171.431 | 2016/09/01 | 3,907,842 | yes |
| SAW16043 | *A. obscura* | SAW | -14.025 | 171.431 | 2016/09/01 | 451,282[1] | yes |
| SAW16044 | *A. obscura* | SAW | -14.025 | 171.431 | 2016/09/01 | 696,640 | yes |
| SAW16045 | *A. obscura* | SAW | -14.025 | 171.431 | 2016/09/01 | 3,211,904 | yes |
| SAW16046 | *A. marmorata* | SAW | -13.876 | 171.639 | 2016/09/01 | 6,048,220 | yes |
| SAW16047 | *A. marmorata* | SAW | -13.876 | 171.639 | 2016/09/02 | 7,660,116 | yes |
| SAW16048 | *A. marmorata* | SAW | -13.876 | 171.639 | 2016/09/02 | 2,626,456 | yes |
| SAW16049 | *A. marmorata* | SAW | -13.876 | 171.639 | 2016/09/02 | 7,328,702 | yes |
| SAW16050 | *A. marmorata* | SAW | -13.876 | 171.639 | 2016/09/02 | 3,279,138 | yes |
| SAW16051 | *A. marmorata* | SAW | -13.876 | 171.639 | 2016/09/02 | 2,574,796 | yes |
| SAW16052 | *A. marmorata* | SAW | -13.876 | 171.639 | 2016/09/02 | 3,483,700 | yes |
| SAW16054 | *A. marmorata* | SAW | -13.876 | 171.639 | 2016/09/02 | 5,107,548 | yes |
| SAW16055 | *A. marmorata* | SAW | -13.876 | 171.639 | 2016/09/02 | 5,252,892 | yes |
| SAW16056 | *A. marmorata* | SAW | -13.876 | 171.639 | 2016/09/02 | 5,137,008 | yes |
| SAW16057 | *A. marmorata* | SAW | -13.876 | 171.639 | 2016/09/02 | 1,370,838 | yes |
| SAW16058 | *A. marmorata* | SAW | -13.876 | 171.639 | 2016/09/02 | 3,511,122 | yes |
| SAW17B02 | *A. marmorata* | SAW | -13.968 | 171.862 | 2017/02/23 | 5,261,166 | no |
| SAW17B10 | *A. megastoma* | SAW | -13.968 | 171.862 | 2017/02/24 | 5,240,736 | yes |
| SAW17B13 | *A. megastoma* | SAW | -13.968 | 171.862 | 2017/02/24 | 5,383,126 | partial |
| SAW17B16 | *A. obscura* | SAW | -13.978 | 171.860 | 2017/02/25 | 4,077,174 | partial |
| SAW17B17 | *A. obscura* | SAW | -13.978 | 171.860 | 2017/02/25 | 5,551,346 | partial |
| SAW17B18 | *A. obscura* | SAW | -13.978 | 171.860 | 2017/02/25 | 5,749,306 | partial |
| SAW17B19 | *A. obscura* | SAW | -13.978 | 171.860 | 2017/02/25 | 7,420,046 | partial |
| SAW17B27 | *A. marmorata* | SAW | -13.968 | 171.862 | 2017/02/27 | 7,417,178 | partial |
| SAW17B48 | *A. megastoma* | SAW | -14.026 | 171.714 | 2017/03/01 | 7,218,260 | partial |
| SAW17B49 | *A. marmorata* | SAW | -14.026 | 171.714 | 2017/03/01 | 3,919,036 | partial |
| SAW17B55 | *A. megastoma* | SAW | -13.992 | 171.588 | 2017/03/09 | 4,638,202 | partial |

#### Supplementary Table 1 (continued)

| Specimen ID | Species | Site | Lat. | Lon. | Date | # reads | Morphology |
|---|---|---|---|---|---|---|---|
| SAW17B56 | *A. marmorata* | SAW | -13.968 | 171.862 | 2017/02/27 | 6,826,590 | partial |
| SAW17B57 | *A. marmorata* | SAW | -13.968 | 171.862 | 2017/02/27 | 3,774,558 | partial |
| SAW17B58 | *A. marmorata* | SAW | -13.968 | 171.862 | 2017/02/27 | 6,366,828 | partial |
| SOK16354 | *A. marmorata* | SOK | -8.060 | 156.973 | 2016/na/na | 5,215,368 | no |
| SOL16V01 | *A. marmorata* | SOL | -7.832 | 156.715 | 2016/na/na | 3,542,070 | no |
| SOL16V02 | *A. marmorata* | SOL | -7.832 | 156.715 | 2016/na/na | 4,502,312 | no |
| SOL16V03 | *A. marmorata* | SOL | -7.861 | 156.696 | 2016/na/na | 7,042,480 | no |
| SOL16V04 | *A. marmorata* | SOL | -7.861 | 156.696 | 2016/na/na | 7,799,996 | no |
| SOL16V06 | *A. marmorata* | SOL | -7.861 | 156.696 | 2016/na/na | 3,521,598 | no |
| SOL16V07 | *A. marmorata* | SOL | -7.861 | 156.696 | 2016/na/na | 4,174,210 | no |
| SOL16V08 | *A. marmorata* | SOL | -7.861 | 156.696 | 2016/na/na | 7,173,740 | no |
| SOL16V10 | *A. marmorata* | SOL | -7.861 | 156.696 | 2016/na/na | 998,684 | no |
| SOL16V11 | *A. marmorata* | SOL | -7.861 | 156.696 | 2016/na/na | 681,170 | no |
| SOL16V12 | *A. marmorata* | SOL | -7.861 | 156.696 | 2016/na/na | 6,531,962 | no |
| SOL16V13 | *A. marmorata* | SOL | -7.861 | 156.696 | 2016/na/na | 8,609,022 | no |
| SON16364 | *A. marmorata* | SON | -8.804 | 158.203 | 2016/na/na | 1,516,152 | no |
| SON16370 | *A. marmorata* | SON | -8.804 | 158.203 | 2016/na/na | 4,083,426 | no |
| SON16371 | *A. marmorata* | SON | -8.804 | 158.203 | 2016/na/na | 2,774,878 | no |
| SON16375 | *A. marmorata* | SON | -8.764 | 158.004 | 2016/na/na | 1,279,182[1] | no |
| SON16376 | *A. marmorata* | SON | -8.804 | 158.203 | 2016/na/na | 1,854,150[1] | no |
| SOR16R01 | *A. marmorata* | SOR | -8.055 | 156.582 | 2016/na/na | 4,643,324 | no |
| SOR16R02 | *A. marmorata* | SOR | -8.055 | 156.582 | 2016/na/na | 5,046,382 | no |
| SOR16R03 | *A. marmorata* | SOR | -8.055 | 156.582 | 2016/na/na | 4,317,646 | no |
| SOR16R06 | *A. marmorata* | SOR | -8.055 | 156.582 | 2016/na/na | 5,291,718 | no |
| SOR16R07 | *A. marmorata* | SOR | -8.055 | 156.582 | 2016/na/na | 5,497,242 | no |
| SOR16R09 | *A. marmorata* | SOR | -8.084 | 156.600 | 2016/na/na | 16,663,830 | no |
| SOR16R12 | *A. marmorata* | SOR | -8.084 | 156.600 | 2016/na/na | 6,932,478 | no |
| SOR16R13 | *A. marmorata* | SOR | -8.036 | 156.536 | 2016/na/na | 7,161,788 | no |
| SOR16R20 | *A. marmorata* | SOR | -8.036 | 156.536 | 2016/na/na | 5,295,472 | no |
| SOR16R21 | *A. marmorata* | SOR | -8.036 | 156.536 | 2016/na/na | 5,314,182 | no |
| SOR16R22 | *A. marmorata* | SOR | -8.036 | 156.536 | 2016/na/na | 2,375,032 | no |
| SOR16R23 | *A. marmorata* | SOR | -8.036 | 156.536 | 2016/na/na | 6,237,660 | no |
| SOV16374 | *A. megastoma* | SOV | -8.817 | 158.189 | 2016/na/na | 4,844,384 | no |
| SOV16377 | *A. marmorata* | SOV | -8.764 | 158.004 | 2016/na/na | 5,086,458 | no |
| TAI15001 | *A. marmorata* | TAI | 24.716 | 121.835 | 2015/06/05 | 3,759,770 | no |
| TAI15002 | *A. marmorata* | TAI | 24.716 | 121.835 | 2015/06/05 | 2,931,064 | no |
| TAI15003 | *A. marmorata* | TAI | 24.716 | 121.835 | 2015/06/05 | 5,276,728 | no |
| TAI15004 | *A. marmorata* | TAI | 24.716 | 121.835 | 2015/06/05 | 3,095,594 | no |
| TAI15005 | *A. marmorata* | TAI | 24.716 | 121.835 | 2015/06/05 | 4,486,336 | no |
| TAI15006 | *A. marmorata* | TAI | 24.716 | 121.835 | 2015/06/05 | 4,999,952 | no |
| TAI15007 | *A. marmorata* | TAI | 24.716 | 121.835 | 2015/06/05 | 5,063,884 | no |
| TAI15008 | *A. marmorata* | TAI | 24.716 | 121.835 | 2015/06/05 | 4,908,062 | no |
| TAI15009 | *A. marmorata* | TAI | 24.716 | 121.835 | 2015/06/05 | 5,224,604 | no |
| TAI15010 | *A. marmorata* | TAI | 24.716 | 121.835 | 2015/06/05 | 4,291,306 | no |
| TAI15011 | *A. marmorata* | TAI | 24.716 | 121.835 | 2015/06/05 | 1,850,660 | no |
| TAI15012 | *A. marmorata* | TAI | 24.716 | 121.835 | 2015/06/05 | 4,193,860 | no |
| TAI15013 | *A. marmorata* | TAI | 24.716 | 121.835 | 2015/06/05 | 4,946,272 | no |

#### Supplementary Table 1 (continued)

| Specimen ID | Species | Site | Lat. | Lon. | Date | # reads | Morphology |
|---|---|---|---|---|---|---|---|
| TAI15014 | *A. marmorata* | TAI | 24.716 | 121.835 | 2015/06/05 | 3,472,668 | no |
| TAI15015 | *A. marmorata* | TAI | 24.716 | 121.835 | 2015/06/05 | 2,665,630 | no |
| TAI15016 | *A. marmorata* | TAI | 24.716 | 121.835 | 2015/06/05 | 4,922,922 | no |
| TAI15017 | *A. marmorata* | TAI | 24.716 | 121.835 | 2015/06/05 | 4,966,356 | no |
| TAI15018 | *A. marmorata* | TAI | 24.716 | 121.835 | 2015/06/05 | 3,745,542 | no |
| TAI15019 | *A. marmorata* | TAI | 24.716 | 121.835 | 2015/06/05 | 4,070,564 | no |
| TAI15020 | *A. marmorata* | TAI | 24.716 | 121.835 | 2015/06/05 | 3,253,040 | no |
| TAI15021 | *A. marmorata* | TAI | 24.716 | 121.835 | 2015/06/05 | 4,451,956 | no |
| TAI15022 | *A. marmorata* | TAI | 24.716 | 121.835 | 2015/06/05 | 3,582,168 | no |
| TAI15023 | *A. marmorata* | TAI | 24.716 | 121.835 | 2015/06/05 | 5,037,786 | no |
| TAI15024 | *A. marmorata* | TAI | 24.716 | 121.835 | 2015/06/05 | 6,443,022 | no |
| TAI15025 | *A. marmorata* | TAI | 24.716 | 121.835 | 2015/06/05 | 5,691,686 | no |
| TAI15026 | *A. marmorata* | TAI | 24.716 | 121.835 | 2015/06/05 | 6,249,920 | no |
| TAI15027 | *A. marmorata* | TAI | 24.716 | 121.835 | 2015/06/05 | 6,987,072 | no |
| TAI15028 | *A. marmorata* | TAI | 24.716 | 121.835 | 2015/06/05 | 4,543,658 | no |
| TAI15029 | *A. marmorata* | TAI | 24.716 | 121.835 | 2015/06/05 | 5,325,064 | no |
| TAI15030 | *A. marmorata* | TAI | 24.716 | 121.835 | 2015/06/05 | 5,234,126 | no |
| VAG12001 | *A. marmorata* | VAG | -14.275 | 167.548 | 2012/01/17 | 2,943,378 | yes |
| VAG12002 | *A. marmorata* | VAG | -14.275 | 167.548 | 2012/01/18 | 1,740,398 | yes |
| VAG12003 | *A. marmorata* | VAG | -14.275 | 167.548 | 2012/01/19 | 1,406,994 | yes |
| VAG12004 | *A. megastoma* | VAG | -14.275 | 167.548 | 2012/01/21 | 6,051,382 | yes |
| VAG12005 | *A. megastoma* | VAG | -14.275 | 167.548 | 2012/01/21 | 6,977,268 | yes |
| VAG12006 | *A. megastoma* | VAG | -14.275 | 167.548 | 2012/01/21 | 2,449,900 | yes |
| VAG12007 | *A. megastoma* | VAG | -14.275 | 167.548 | 2012/01/21 | 3,009,804 | yes |
| VAG12008 | *A. megastoma* | VAG | -14.275 | 167.548 | 2012/01/21 | 2,672,694 | yes |
| VAG12009 | *A. megastoma* | VAG | -14.275 | 167.548 | 2012/01/21 | 132,204[1] | yes[2] |
| VAG12010 | *A. megastoma* | VAG | -14.275 | 167.548 | 2012/01/21 | 1,859,440 | yes |
| VAG12011 | *A. megastoma* | VAG | -14.275 | 167.548 | 2012/01/21 | 2,859,824 | yes |
| VAG12012 | *A. marmorata* | VAG | -14.275 | 167.548 | 2012/01/21 | 885,610 | yes |
| VAG12013 | *A. megastoma* | VAG | -14.275 | 167.548 | 2012/01/21 | 4,839,628 | yes |
| VAG12014 | *A. megastoma* | VAG | -14.275 | 167.548 | 2012/01/22 | 3,150,160 | yes |
| VAG12015 | *A. megastoma* | VAG | -14.275 | 167.548 | 2012/01/22 | 2,285,658 | yes |
| VAG12016 | *A. marmorata* | VAG | -14.275 | 167.548 | 2012/01/22 | 2,609,394 | yes |
| VAG12018 | *A. marmorata* | VAG | -14.275 | 167.548 | 2012/01/22 | 2,795,482 | yes |
| VAG12019 | *A. marmorata* | VAG | -14.275 | 167.548 | 2012/01/22 | 3,403,952 | yes |
| VAG12020 | *A. megastoma* | VAG | -14.275 | 167.548 | 2012/01/22 | 8,602,088 | yes |
| VAG12021 | *A. megastoma* | VAG | -14.275 | 167.548 | 2012/01/22 | 254,536[1] | yes[2] |
| VAG12022 | *A. marmorata* | VAG | -14.275 | 167.548 | 2012/01/22 | 1,420,164 | yes |
| VAG12023 | *A. interioris* | VAG | -14.275 | 167.548 | 2012/01/23 | 3,310,828 | yes |
| VAG12024 | *A. marmorata* | VAG | -14.275 | 167.548 | 2012/01/22 | 2,999,122 | yes |
| VAG12025 | *A. marmorata* | VAG | -14.275 | 167.548 | 2012/01/22 | 5,478,366 | yes |
| VAG12026 | *A. marmorata* | VAG | -14.275 | 167.548 | 2012/01/22 | 219,894[1] | yes[2] |
| VAG12027 | *A. megastoma* | VAG | -14.275 | 167.548 | 2012/01/22 | 6,337,898 | yes |
| VAG12028 | *A. marmorata* | VAG | -14.275 | 167.548 | 2012/01/22 | 260,506[1] | yes[2] |
| VAG12029 | *A. marmorata* | VAG | -14.275 | 167.548 | 2012/01/24 | 3,218,850 | yes |
| VAG12030 | *A. marmorata* | VAG | -14.275 | 167.548 | 2012/01/24 | 818,028 | yes |
| VAG12031 | *A. marmorata* | VAG | -14.275 | 167.548 | 2012/01/24 | 6,862,982 | yes |

#### Supplementary Table 1 (continued)

| Specimen ID | Species | Site | Lat. | Lon. | Date | # reads | Morphology |
|---|---|---|---|---|---|---|---|
| VAG12032 | *A. megastoma* | VAG | -14.275 | 167.548 | 2012/01/24 | 255,918[1] | yes[2] |
| VAG12033 | *A. marmorata* | VAG | -14.275 | 167.548 | 2012/01/24 | 725,846 | yes[2] |
| VAG12034 | *A. megastoma* | VAG | -14.275 | 167.548 | 2012/01/24 | 8,292,228 | yes |
| VAG12035 | *A. marmorata* | VAG | -14.275 | 167.548 | 2012/01/24 | 528,696[1] | yes[2] |
| VAG12036 | *A. marmorata* | VAG | -14.275 | 167.548 | 2012/01/24 | 11,598,732 | yes |
| VAG12037 | *A. marmorata* | VAG | -14.275 | 167.548 | 2012/01/24 | 2,505,822 | yes[2] |
| VAG12038 | *A. megastoma* | VAG | -14.275 | 167.548 | 2012/01/24 | 4,382,324 | partial |
| VAG12039 | *A. marmorata* | VAG | -14.261 | 167.605 | 2012/01/28 | 18,191,152 | yes |
| VAG12040 | *A. marmorata* | VAG | -14.261 | 167.605 | 2012/01/31 | 7,749,912 | yes |
| VAG12041 | *A. megastoma* | VAG | -14.261 | 167.605 | 2012/01/31 | 3,770,604 | yes |
| VAG12044 | *A. megastoma* | VAG | -14.261 | 167.605 | 2012/02/01 | 4,910,168 | yes |
| VAG12045 | *A. marmorata* | VAG | -14.261 | 167.605 | 2012/02/01 | 18,655,124 | yes |
| VAG12046 | *A. megastoma* | VAG | -14.261 | 167.605 | 2012/02/01 | 2,129,488 | yes[2] |
| VAG12047 | *A. megastoma* | VAG | -14.261 | 167.605 | 2012/02/01 | 1,262,984 | yes[2] |
| VAG12049 | *A. obscura* | VAG | -14.261 | 167.605 | 2012/01/31 | 5,359,114 | yes |
| VAG12050 | *A. obscura* | VAG | -14.261 | 167.605 | 2012/02/02 | 2,618,778 | yes |
| VAG12051 | *A. megastoma* | VAG | -14.261 | 167.605 | 2012/02/02 | 8,358,494 | partial |
| VAG12052 | *A. marmorata* | VAG | -14.261 | 167.605 | 2012/02/02 | 11,983,860 | partial |
| VAG12053 | *A. marmorata* | VAG | -14.261 | 167.605 | 2012/02/02 | 3,660,386 | partial |
| VAG12054 | *A. marmorata* | VAG | -14.261 | 167.605 | 2012/02/01 | 494,650[1] | yes[2] |
| VAG12055 | *A. marmorata* | VAG | -14.261 | 167.605 | 2012/02/01 | 6,158,622 | yes |
| VAG12056 | *A. megastoma* | VAG | -14.261 | 167.605 | 2012/02/01 | 6,352,762 | yes |
| VAG12059 | *A. marmorata* | VAG | -14.261 | 167.605 | 2012/01/31 | 983,610 | yes |
| VAG12060 | *A. obscura* | VAG | -14.261 | 167.605 | 2012/02/01 | 1,435,834 | yes |
| VAG12061 | *A. obscura* | VAG | -14.261 | 167.605 | 2012/02/01 | 3,400,270 | yes |
| VAG12062 | *A. megastoma* | VAG | -14.261 | 167.605 | 2012/02/02 | 4,258,918 | yes |
| VAG12063 | *A. marmorata* | VAG | -14.261 | 167.605 | 2012/02/02 | 5,949,864 | yes |
| VAG12064 | *A. megastoma* | VAG | -14.261 | 167.605 | 2012/02/02 | 3,143,290 | yes |
| VAG12065 | *A. megastoma* | VAG | -14.261 | 167.605 | 2012/02/02 | 3,238,970 | yes |
| VAG12067 | *A. marmorata* | VAG | -14.261 | 167.605 | 2012/02/01 | 5,545,036 | yes |
| VAG12068 | *A. marmorata* | VAG | -14.261 | 167.605 | 2012/02/02 | 7,125,294 | yes |
| VAG13070 | *A. marmorata* | VAG | -14.261 | 167.605 | 2013/03/01 | 3,508,578 | yes |
| VAG13071 | *A. marmorata* | VAG | -14.261 | 167.605 | 2013/03/01 | 3,910,052 | yes |
| VAG13072 | *A. marmorata* | VAG | -14.261 | 167.605 | 2013/03/22 | 5,860,892 | yes[2] |
| VAG13073 | *A. obscura* | VAG | -14.261 | 167.605 | 2013/03/28 | 8,105,260 | yes |
| VAG13074 | *A. obscura* | VAG | -14.261 | 167.605 | 2013/03/28 | 7,979,904 | yes |
| VAG13075 | *A. megastoma* | VAG | -14.261 | 167.605 | 2013/03/27 | 4,794,394 | yes |
| VAG13076 | *A. megastoma* | VAG | -14.261 | 167.605 | 2013/03/30 | 3,882,404 | yes |
| VAG13077 | *A. marmorata* | VAG | -14.261 | 167.605 | 2013/03/27 | 6,948,582 | yes |
| VAG13078 | *A. marmorata* | VAG | -14.261 | 167.605 | 2013/03/26 | 6,147,476 | yes |
| VAG13079 | *A. marmorata* | VAG | -14.261 | 167.605 | 2013/03/26 | 7,965,414 | yes |
| VAG13080 | *A. obscura* | VAG | -14.261 | 167.605 | 2013/04/01 | 6,240,488 | yes |
| VAG13081 | *A. marmorata* | VAG | -14.261 | 167.605 | 2013/04/02 | 5,948,510 | yes |
| VAG13082 | *A. marmorata* | VAG | -14.261 | 167.605 | 2013/04/01 | 4,319,756 | yes |
| VAG13083 | *A. marmorata* | VAG | -14.261 | 167.605 | 2013/04/01 | 25,480[1] | yes[2] |
| VAG13084 | *A. obscura* | VAG | -14.261 | 167.605 | 2013/04/01 | 2,637,702 | yes |
| VAG13085 | *A. marmorata* | VAG | -14.261 | 167.605 | 2013/04/02 | 6,013,544 | yes |

**Supplementary Table 1 (continued)**

| Specimen ID | Species | Site | Lat. | Lon. | Date | # reads | Morphology |
|---|---|---|---|---|---|---|---|
| VAG13086 | *A. megastoma* | VAG | -14.261 | 167.605 | 2013/04/03 | 3,298,274 | yes |
| VAG13087 | *A. marmorata* | VAG | -14.261 | 167.605 | 2013/04/03 | 1,115,480 | partial |

**Supplementary Table 2:** Per-species population-genetic parameters for tropical eel species. Reported parameters are calculated for the dataset for phylogenetic analyses (before reducing it to maximally five individuals per species). Heterozygosity ($h$), nucleotide diversity ($\pi$), and population mutation rate ($\Theta$) were calculated twice; first assuming that all missing data are invariable, and second assuming that missing data mask genotypes that are equally variable as the observed ones. For *A. marmorata*, all parameters were calculated first for all individuals jointly and then for each of four populations: WIO, western Indian Ocean (sampling sites AFC, AFC, MAY, REU); SCS, South China Sea (sampling sites PHP, PHC, TAI); WSP, western South Pacific (sampling sites BOU, SO, VAG, NCA, SAA, SAW). $n$, Number of individuals used in genomic analyses.

| Species | $n$ | Completeness | # variable sites | $h$ ($\times10^{-3}$) | $\pi$ ($\times10^{-3}$) | $\Theta$ ($\times10^{-3}$) |
|---|---|---|---|---|---|---|
| *A. marmorata* | 325 | 0.790 | 373,382 | 0.32/0.40 | 0.50/0.53 | 2.90/4.10 |
| *A. marmorata* (WIO) | 42 | 0.877 | 35,094 | 0.19/0.21 | 0.22/0.23 | 0.38/0.61 |
| *A. marmorata* (Java) | 21 | 0.599 | 25,187 | 0.14/0.24 | 0.26/0.28 | 0.32/0.82 |
| *A. marmorata* (SCS) | 63 | 0.877 | 201,746 | 0.53/0.61 | 0.69/0.70 | 2.04/2.65 |
| *A. marmorata* (WSP) | 199 | 0.765 | 191,935 | 0.30/0.39 | 0.44/0.46 | 1.60/3.23 |
| *A. luzonensis* | 20 | 0.737 | 113,858 | 0.72/0.98 | 1.12/1.19 | 1.47/2.53 |
| *A. bicolor* | 4 | 0.513 | 45,975 | 0.67/1.30 | 1.31/1.49 | 0.97/1.96 |
| *A. obscura* | 36 | 0.723 | 87,297 | 0.34/0.48 | 0.56/0.58 | 0.99/1.88 |
| *A. interioris* | 3 | 0.731 | 28,474 | 0.57/0.78 | 0.82/0.88 | 0.68/0.98 |
| *A. megastoma* | 41 | 0.738 | 127,411 | 0.43/0.58 | 0.71/0.71 | 1.40/2.41 |
| *A. mossambica* | 1 | 0.756 | 19,273 | 1.06/1.40 | 1.06/1.40 | 1.06/1.40 |

**Supplementary Table 3:** Per-species population-genetic parameters for tropical eel species. As Supplementary Table 2, but using the dataset for population-genetic analyses.

| Species | $n$ | Completeness | # variable sites | $h$ ($\times10^{-3}$) | $\pi$ ($\times10^{-3}$) | $\Theta$ ($\times10^{-3}$) |
|---|---|---|---|---|---|---|
| *A. marmorata* | 325 | 0.910 | 146,998 | 0.65/0.72 | 0.86/0.88 | 1.14/1.20 |
| *A. marmorata* (WIO) | 42 | 0.948 | 46,793 | 0.53/0.56 | 0.57/0.57 | 0.51/0.67 |
| *A. marmorata* (Java) | 21 | 0.826 | 44,100 | 0.49/0.59 | 0.60/0.60 | 0.56/0.89 |
| *A. marmorata* (SCS) | 63 | 0.940 | 68,595 | 0.70/0.75 | 0.83/0.83 | 0.69/0.91 |
| *A. marmorata* (WSP) | 199 | 0.901 | 139,822 | 0.68/0.76 | 0.81/0.81 | 1.17/1.28 |
| *A. luzonensis* | 20 | 0.859 | 53,935 | 0.62/0.72 | 0.80/0.84 | 0.69/0.99 |
| *A. bicolor* | 4 | 0.721 | 13,436 | 0.23/0.31 | 0.34/0.37 | 0.28/0.43 |
| *A. obscura* | 36 | 0.844 | 60,983 | 0.40/0.48 | 0.48/0.54 | 0.69/0.98 |
| *A. interioris* | 3 | 0.877 | 12,487 | 0.27/0.30 | 0.32/0.34 | 0.30/0.35 |
| *A. megastoma* | 41 | 0.818 | 72,415 | 0.56/0.69 | 0.74/0.80 | 0.80/1.18 |
| *A. mossambica* | 1 | 0.777 | 2,595 | 0.14/0.18 | 0.14/0.18 | 0.14/0.18 |

**Supplementary Table 4:** Mean pairwise genetic distance between tropical eel species. Genetic distances were calculated as uncorrected p-distances (the proportion of sites at which two sequences are different), based on a concatenated alignment of 20,637 RAD loci. These loci were selected to have no fully missing sequences for the maximally five individuals per species with the overall lowest proportion of missing data; only these individuals are included in the alignment. As p-distances are symmetric, species 1 and 2 are exchangeable. Rows are sorted by p-distance.

| Species 1 | Species 2 | # pairs | Mean p-distance |
|---|---|---|---|
| *A. luzonensis* | *A. marmorata* | 25 | 0.0053 |
| *A. interioris* | *A. luzonensis* | 15 | 0.0060 |
| *A. bicolor* | *A. obscura* | 20 | 0.0064 |
| *A. interioris* | *A. marmorata* | 15 | 0.0065 |
| *A. bicolor* | *A. interioris* | 12 | 0.0067 |
| *A. interioris* | *A. obscura* | 15 | 0.0067 |
| *A. luzonensis* | *A. obscura* | 25 | 0.0072 |
| *A. bicolor* | *A. luzonensis* | 20 | 0.0073 |
| *A. bicolor* | *A. marmorata* | 20 | 0.0076 |
| *A. marmorata* | *A. obscura* | 25 | 0.0076 |
| *A. interioris* | *A. megastoma* | 15 | 0.0079 |
| *A. luzonensis* | *A. megastoma* | 25 | 0.0081 |
| *A. marmorata* | *A. megastoma* | 25 | 0.0082 |
| *A. megastoma* | *A. obscura* | 25 | 0.0088 |
| *A. bicolor* | *A. megastoma* | 20 | 0.0090 |
| *A. megastoma* | *A. mossambica* | 5 | 0.0103 |
| *A. interioris* | *A. mossambica* | 3 | 0.0105 |
| *A. luzonensis* | *A. mossambica* | 5 | 0.0106 |
| *A. marmorata* | *A. mossambica* | 5 | 0.0107 |
| *A. mossambica* | *A. obscura* | 5 | 0.0113 |
| *A. bicolor* | *A. mossambica* | 4 | 0.0116 |

**Supplementary Table 5:** Genome assemblies. The results of the BUSCO analysis are given in the order complete (c), complete and single copy (c+s), complete and duplicated (c+d), fragmented (f), missing (m). A total of 2,586 BUSCOs were searched for each assembly. scf., scaffold; ctg., contig.

| Species | Assembly version | Assembly size | N50 scf. | N50 ctg. | —BUSCOs (c/c+s/c+d/f/m)— | | | | |
|---|---|---|---|---|---|---|---|---|---|
| *A. marmorata* | Uncorrected | 880,647,635 | 64,770 | 15,086 | 2,217 | 2,028 | 189 | 254 | 115 |
| | Pilon-corrected | 882,006,954 | 64,942 | 16,529 | 2,215 | 2,020 | 195 | 254 | 117 |
| *A. megastoma* | Uncorrected | 877,063,880 | 61,871 | 13,547 | 2,203 | 2,013 | 190 | 264 | 119 |
| | Pilon-corrected | 877,765,645 | 61,910 | 14,572 | 2,207 | 2,015 | 192 | 260 | 119 |
| *A. obscura* | Uncorrected | 881,549,187 | 54,844 | 12,017 | 2,161 | 1,977 | 184 | 307 | 118 |
| | Pilon-corrected | 882,390,062 | 54,849 | 12,681 | 2,168 | 1,982 | 186 | 304 | 114 |

**Supplementary Table 6:** Cross-validation (CV) error values of maximum-likelihood ancestry inference with ADMIXTURE.
CV errors were inferred for the models $K = 1$ to $K = 8$ based on 117,638 variable sites for five replicates (R1 to R5) per model.

| Model | R1 | R2 | R3 | R4 | R5 |
|---|---|---|---|---|---|
| $K = 1$ | 0.49028 | 0.49037 | 0.49046 | 0.49041 | 0.49040 |
| $K = 2$ | 0.29901 | 0.29916 | 0.29903 | 0.29902 | 0.29913 |
| $K = 3$ | 0.22883 | 0.22879 | 0.22882 | 0.26443 | 0.22880 |
| $K = 4$ | 0.24704 | 0.19584 | 0.19579 | 0.19590 | 0.24694 |
| $K = 5$ | 0.17819 | 0.17819 | 0.17816 | 0.17822 | 0.17820 |
| $K = 6$ | 0.17491 | 0.17496 | 0.17500 | 0.17500 | 0.17488 |
| $K = 7$ | 0.17614 | 0.17613 | 0.17610 | 0.17614 | 0.17380 |
| $K = 8$ | 0.17508 | 0.17421 | 0.17849 | 0.17435 | 0.17487 |

**Supplementary Table 7:** Hybrid individuals.

Genomic and morphological characteristics of hybrid individuals identified via ancestry painting. Note that for backcrossed hybrids, the identity of maternal and paternal species are ambiguous because either the mother or the father is a hybrid itself; in this case, the name listed as maternal species indicates the species identity of the mitochondrial genome. $h_{\text{fixed}}$, heterozygosity at sites fixed between parental species; $f_{\text{m,genome}}$, proportion of genome derived from the maternal species; $f_{\text{m,morphology}}$, similarity to morphology of maternal species, relative to morphology of paternal species.

| Specimen ID | Maternal species | Paternal species | Date | $h_{\text{fixed}}$ | $f_{\text{m,genome}}$ | $f_{\text{m,morphology}}$ | Interpretation |
|---|---|---|---|---|---|---|---|
| BOU15031 | *A. marmorata* | *A. megastoma* | 04/2015 | 0.479 | 0.250 | 0.257 | Backcross[1] |
| SAA16011 | *A. marmorata* | *A. megastoma* | 08/2016 | 0.974 | 0.506 | NA | F1 |
| SAA16012 | *A. marmorata* | *A. megastoma* | 08/2016 | 0.966 | 0.500 | NA | F1 |
| SAA16013 | *A. marmorata* | *A. megastoma* | 08/2016 | 0.981 | 0.510 | NA | F1 |
| SAA16024 | *A. marmorata* | *A. megastoma* | 08/2016 | 0.550 | 0.716 | NA | Backcross[2] |
| SAA16027 | *A. marmorata* | *A. megastoma* | 08/2016 | 0.956 | 0.500 | NA | F1 |
| SAW17B27 | *A. marmorata* | *A. megastoma* | 02/2017 | 0.972 | 0.506 | NA | F1 |
| SAW17B49 | *A. marmorata* | *A. megastoma* | 03/2017 | 0.969 | 0.512 | NA | F1 |
| VAG12012 | *A. marmorata* | *A. megastoma* | 01/2012 | 0.971 | 0.515 | 0.524 | F1 |
| VAG12018 | *A. marmorata* | *A. megastoma* | 01/2012 | 0.977 | 0.500 | 0.685 | F1 |
| VAG12019 | *A. marmorata* | *A. megastoma* | 01/2012 | 0.975 | 0.508 | 0.572 | F1 |
| VAG12024 | *A. marmorata* | *A. megastoma* | 01/2012 | 0.510 | 0.263 | 0.504 | Backcross[1] |
| VAG12029 | *A. marmorata* | *A. megastoma* | 01/2012 | 0.459 | 0.767 | 0.746 | Backcross[2] |
| VAG12037 | *A. marmorata* | *A. megastoma* | 01/2012 | 0.817 | 0.575 | NA | F1 |
| VAG12044 | *A. megastoma* | *A. marmorata* | 02/2012 | 0.973 | 0.514 | 0.537 | F1 |
| VAG12053 | *A. marmorata* | *A. megastoma* | 02/2012 | 0.542 | 0.705 | NA | Backcross[2] |
| VAG12055 | *A. marmorata* | *A. megastoma* | 02/2012 | 0.460 | 0.250 | 0.545 | Backcross[1] |
| VAG13071 | *A. marmorata* | *A. megastoma* | 03/2013 | 0.954 | 0.512 | 0.580 | F1 |
| VAG13078 | *A. marmorata* | *A. megastoma* | 03/2013 | 0.972 | 0.500 | 0.470 | F1 |
| VAG13087 | *A. marmorata* | *A. megastoma* | 04/2013 | 0.584 | 0.324 | NA | Backcross[1] |
| VAG12040 | *A. marmorata* | *A. obscura* | 02/2012 | 0.951 | 0.507 | 0.540 | F1 |
| VAG12045 | *A. marmorata* | *A. obscura* | 02/2012 | 0.899 | 0.510 | 0.618 | F1 |
| VAG13077 | *A. marmorata* | *A. obscura* | 03/2013 | 0.957 | 0.500 | 0.537 | F1 |
| VAG12049 | *A. obscura* | *A. megastoma* | 01/2012 | 0.977 | 0.506 | 0.512 | F1 |
| BOU15017 | *A. marmorata* | *A. interioris* | 04/2015 | 0.939 | 0.520 | 0.666 | F1 |

[1]The mitochondrial genome of *A. marmorata* and the $f_{\text{m,genome}}$ around 0.25 indicate that the mother of the mother of this individual was an *A. marmorata* but all other grantparents were *A. megastoma*.

[2]The mitochondrial genome of *A. marmorata* and the $f_{\text{m,genome}}$ around 0.75 indicate that one of the grandparents of this individual, but not the mother of the mother, was an *A. megastoma* and all other grandparents were *A. marmorata*.

**Supplementary Table 8:** Frequency of hybrids at sampling sites.
$n_t$, number of sampled specimens; $n_g$, number of individuals used in genomic analyses, excluding those with low sequence quality.

| Site | Location | Country | $n_t$ | $n_g$ | # hybrids | Frequency (%) |
|---|---|---|---|---|---|---|
| AFC | Eastern Cape | South Africa | 15 | 14 | 0 | 0 |
| AFS | Lubombo | Swaziland | 1 | 1 | 0 | 0 |
| MAY | Mayotte | France | 18 | 18 | 0 | 0 |
| REU | Réunion | France | 10 | 10 | 0 | 0 |
| JAV | Java | Indonesia | 30 | 27 | 0 | 0 |
| PHP | Pagadian | Philippines | 27 | 27 | 0 | 0 |
| PHC | Cagayan | Philippines | 31 | 26 | 0 | 0 |
| TAI | Yilan County | Taiwan | 30 | 30 | 0 | 0 |
| BOU | Bougainville | Papua New Guinea | 30 | 30 | 2 | 6.7 |
| SOK | Kolombangara | Solomon Islands | 1 | 1 | 0 | 0 |
| SOL | Vella Lavella | Solomon Islands | 11 | 11 | 0 | 0 |
| SON | Nggatokae | Solomon Islands | 5 | 3 | 0 | 0 |
| SOR | Ranongga | Solomon Islands | 12 | 12 | 0 | 0 |
| SOV | Vangunu | Solomon Islands | 2 | 2 | 0 | 0 |
| VAG | Gaua | Vanuatu | 79 | 71 | 16 | 22.5 |
| NCA | New Caledonia | France | 45 | 45 | 0 | 0 |
| SAW | Upolu | Samoa | 71 | 67 | 2 | 3.0 |
| SAA | Tutuila | American Samoa | 38 | 35 | 5 | 14.3 |
| Total | | | 456 | 430 | 25 | 5.8 |

**Supplementary Table 9:** Introgression statistics for species quartets.
All species quartets compatible with the inferred species tree were tested. Quartet comparisons are sorted by $D$ values. mar, *A. marmorata*; luz, *A. luzonensis*; int, *A. interioris*; obs, *A. obscura*; bic, *A. bicolor*; meg, *A. megastoma*; mos, *A. mossambica*; ang, *A. anguilla*; $n$, number of informative sites; $C_{\mathrm{BBAA}}$, number of "BBAA" sites; $C_{\mathrm{ABBA}}$, number of "ABBA" sites; $C_{\mathrm{BABA}}$, number of "BABA" sites; $D$, Patterson's $D$ statistic (Green *et al.* 2010; Durand *et al.* 2011); $f_4$, the $f_4$ statistic (Reich *et al.* 2009); $p$, $p$ value for $f_4 = 0$ assessed through simulations with the F4 program (Meyer *et al.* 2017). The comparison reported in the last table row is based on WGS reads of a single individual of *A. obscura*, *A. marmorata*, and *A. megastoma*, aligned to the available reference-genome assembly of *A. anguilla* (Jansen *et al.* 2017).

| P1 | P2 | P3 | Outgroup | $n$ | $C_{\mathrm{BBAA}}$ | $C_{\mathrm{ABBA}}$ | $C_{\mathrm{BABA}}$ | $D$ | $f_4$ | $p$ |
|---|---|---|---|---|---|---|---|---|---|---|
| mar | luz | int | mos | 10,290 | 273.1 | 182.7 | 77.1 | 0.406 | -0.0070 | 0.000 |
| mar | luz | int | meg | 13,396 | 335.9 | 207.6 | 98.2 | 0.358 | -0.0054 | 0.000 |
| mar | luz | obs | meg | 15,689 | 412.1 | 186.6 | 93.0 | 0.334 | -0.0043 | 0.000 |
| mar | luz | obs | mos | 12,054 | 358.2 | 162.7 | 83.7 | 0.321 | -0.0048 | 0.000 |
| mar | bic | int | mos | 7,772 | 100.9 | 266.3 | 138.4 | 0.316 | -0.0109 | 0.000 |
| mar | luz | bic | mos | 11,542 | 311.9 | 158.1 | 82.8 | 0.313 | -0.0052 | 0.000 |
| mar | bic | int | meg | 9,680 | 136.6 | 295.3 | 168.6 | 0.273 | -0.0077 | 0.000 |
| mar | luz | bic | meg | 14,793 | 360.6 | 168.2 | 103.4 | 0.239 | -0.0035 | 0.000 |
| mar | obs | int | meg | 10,208 | 137.3 | 307.9 | 197.8 | 0.218 | -0.0051 | 0.000 |
| mar | obs | int | mos | 8,068 | 106.3 | 268.6 | 173.1 | 0.216 | -0.0086 | 0.000 |
| obs | bic | mar | meg | 11,372 | 653.6 | 104.7 | 71.2 | 0.191 | -0.0025 | 0.002 |
| obs | bic | int | mos | 8,304 | 373.7 | 123.8 | 84.1 | 0.191 | -0.0030 | 0.005 |
| obs | bic | int | meg | 10,444 | 471.0 | 125.7 | 86.6 | 0.184 | -0.0026 | 0.003 |
| obs | bic | mar | mos | 9,068 | 555.9 | 98.8 | 68.3 | 0.182 | -0.0016 | 0.078 |
| obs | bic | luz | mos | 12,557 | 487.2 | 113.4 | 80.0 | 0.173 | -0.0022 | 0.008 |
| mar | int | meg | mos | 9,951 | 482.4 | 96.4 | 72.7 | 0.140 | -0.0023 | 0.026 |
| obs | bic | luz | meg | 16,064 | 582.5 | 108.7 | 82.5 | 0.137 | -0.0015 | 0.017 |
| mar | luz | meg | mos | 13,129 | 677.0 | 69.0 | 52.9 | 0.133 | -0.0008 | 0.201 |
| luz | mar | bic | int | 14,675 | 392.7 | 105.4 | 84.5 | 0.110 | -0.0011 | 0.106 |
| luz | bic | int | meg | 14,246 | 133.6 | 228.4 | 191.0 | 0.089 | -0.0015 | 0.062 |
| luz | int | meg | mos | 13,632 | 550.4 | 82.4 | 70.2 | 0.080 | -0.0007 | 0.192 |
| luz | bic | int | mos | 11,133 | 111.4 | 197.5 | 168.3 | 0.080 | -0.0022 | 0.042 |
| mar | bic | meg | mos | 11,134 | 441.3 | 110.9 | 95.0 | 0.077 | -0.0003 | 0.430 |
| luz | mar | obs | int | 15,500 | 417.8 | 111.7 | 96.5 | 0.073 | -0.0003 | 0.406 |
| mar | obs | meg | mos | 11,647 | 458.9 | 126.1 | 110.0 | 0.068 | -0.0009 | 0.241 |
| bic | obs | mar | int | 11,303 | 520.4 | 80.0 | 73.0 | 0.046 | -0.0007 | 0.261 |
| obs | bic | meg | mos | 11,761 | 813.0 | 64.7 | 59.5 | 0.042 | -0.0002 | 0.447 |
| bic | obs | luz | int | 15,856 | 526.5 | 78.1 | 72.1 | 0.040 | -0.0010 | 0.141 |
| obs | int | meg | mos | 11,017 | 557.5 | 96.2 | 90.8 | 0.029 | -0.0011 | 0.137 |
| luz | bic | meg | mos | 14,602 | 480.2 | 97.0 | 93.1 | 0.020 | 0.0002 | 0.416 |
| luz | obs | int | meg | 15,143 | 144.2 | 227.1 | 221.7 | 0.012 | 0.0005 | 0.300 |
| bic | int | meg | mos | 10,451 | 535.3 | 84.0 | 82.0 | 0.012 | -0.0007 | 0.213 |
| luz | obs | meg | mos | 15,405 | 507.8 | 107.9 | 106.2 | 0.008 | -0.0001 | 0.461 |
| luz | obs | int | mos | 11,638 | 107.7 | 197.6 | 198.7 | -0.003 | -0.0007 | 0.303 |
| obs | luz | meg | mos | 15,405 | 507.8 | 106.2 | 107.9 | -0.008 | 0.0001 | 0.463 |
| int | bic | meg | mos | 10,451 | 535.3 | 82.0 | 84.0 | -0.012 | 0.0007 | 0.227 |

#### Supplementary Table 9 (continued)

| P1 | P2 | P3 | Outgroup | $n$ | $C_{\mathrm{BBAA}}$ | $C_{\mathrm{ABBA}}$ | $C_{\mathrm{BABA}}$ | $D$ | $f_4$ | $p$ |
|---|---|---|---|---:|---:|---:|---:|---:|---:|---:|
| bic | luz | meg | mos | 14,602 | 480.2 | 93.1 | 97.0 | -0.020 | -0.0002 | 0.420 |
| int | obs | meg | mos | 11,017 | 557.5 | 90.8 | 96.2 | -0.029 | 0.0011 | 0.164 |
| obs | bic | luz | int | 15,856 | 526.5 | 72.1 | 78.1 | -0.040 | 0.0010 | 0.141 |
| bic | obs | meg | mos | 11,761 | 813.0 | 59.5 | 64.7 | -0.042 | 0.0002 | 0.420 |
| obs | bic | mar | int | 11,303 | 520.4 | 73.0 | 80.0 | -0.046 | 0.0007 | 0.263 |
| obs | mar | meg | mos | 11,647 | 458.9 | 110.0 | 126.1 | -0.068 | 0.0009 | 0.232 |
| mar | luz | obs | int | 15,500 | 417.8 | 96.5 | 111.7 | -0.073 | 0.0003 | 0.384 |
| bic | mar | meg | mos | 11,134 | 441.3 | 95.0 | 110.9 | -0.077 | 0.0003 | 0.403 |
| int | luz | meg | mos | 13,632 | 550.4 | 70.2 | 82.4 | -0.080 | 0.0007 | 0.193 |
| mar | luz | bic | int | 14,675 | 392.7 | 84.5 | 105.4 | -0.110 | 0.0011 | 0.119 |
| luz | mar | meg | mos | 14,059 | 690.1 | 53.8 | 69.0 | -0.124 | 0.0008 | 0.183 |
| bic | obs | luz | meg | 16,064 | 582.5 | 82.5 | 108.7 | -0.137 | 0.0015 | 0.013 |
| int | mar | meg | mos | 9,951 | 482.4 | 72.7 | 96.4 | -0.140 | 0.0023 | 0.025 |
| bic | obs | luz | mos | 12,557 | 487.2 | 80.0 | 113.4 | -0.173 | 0.0022 | 0.009 |
| bic | obs | mar | mos | 9,068 | 555.9 | 68.3 | 98.8 | -0.182 | 0.0016 | 0.074 |
| bic | obs | int | meg | 10,444 | 471.0 | 86.6 | 125.7 | -0.184 | 0.0026 | 0.006 |
| bic | obs | mar | meg | 11,372 | 653.6 | 71.2 | 104.7 | -0.191 | 0.0025 | 0.002 |
| bic | obs | int | mos | 8,304 | 373.7 | 84.1 | 123.8 | -0.191 | 0.0030 | 0.006 |
| luz | mar | bic | meg | 14,793 | 360.6 | 103.4 | 168.2 | -0.239 | 0.0035 | 0.000 |
| luz | mar | bic | mos | 11,542 | 311.9 | 82.8 | 158.1 | -0.313 | 0.0052 | 0.000 |
| luz | mar | obs | mos | 12,054 | 358.2 | 83.7 | 162.7 | -0.321 | 0.0048 | 0.000 |
| luz | mar | obs | meg | 15,689 | 412.1 | 93.0 | 186.6 | -0.334 | 0.0043 | 0.000 |
| luz | mar | int | meg | 13,396 | 335.9 | 98.2 | 207.6 | -0.358 | 0.0054 | 0.000 |
| luz | mar | int | mos | 10,290 | 273.1 | 77.1 | 182.7 | -0.406 | 0.0070 | 0.000 |
| obs | mar | meg | ang | 23,165,451 | 1638567.0 | 596786.0 | 587910.0 | 0.007 | — | — |

**Supplementary Table 10:** Introgression statistics for quartets of species and *A. marmorata* populations.
As Supplementary Table 9, but for comparisons involving individual *A. marmorata* populations. Only a single quartet involving *A. marmorata* appeared significant in Supplementary Table 9 and was tested further with separate populations. In addition, the possibility of different degrees of introgression between *A. luzonensis* and the four *A. marmorata* populations was explored because *A. luzonensis* appeared to share more coancestry with the South China Sea population of *A. marmorata* than with other populations in the fineRADstructure analysis (Supplementary Figure 8). O, Outgroup; WIO, western Indian Ocean (sampling sites AFC, AFC, MAY, REU); SCS, South China Sea (sampling sites PHP, PHC, TAI); WSP, western South Pacific (sampling sites BOU, SO, VAG, NCA, SAA, SAW).

| P1 | P2 | P3 | O | $n$ | $C_{\mathrm{BBAA}}$ | $C_{\mathrm{ABBA}}$ | $C_{\mathrm{BABA}}$ | $D$ | $f_4$ | $p$ |
|---|---|---|---|---|---|---|---|---|---|---|
| obs | bic | mar (WIO) | meg | 11,104 | 653.0 | 105.5 | 71.5 | 0.192 | -0.0016 | 0.042 |
| obs | bic | mar (WIO) | mos | 8,869 | 555.0 | 98.8 | 68.8 | 0.179 | -0.0016 | 0.099 |
| obs | bic | mar (Java) | meg | 13,543 | 681.5 | 103.2 | 70.2 | 0.191 | -0.0031 | 0.000 |
| obs | bic | mar (Java) | mos | 8,297 | 508.0 | 89.2 | 62.1 | 0.179 | -0.0025 | 0.008 |
| obs | bic | mar (SCS) | meg | 10,794 | 632.5 | 97.2 | 68.1 | 0.176 | -0.0029 | 0.000 |
| obs | bic | mar (SCS) | mos | 13,340 | 557.4 | 98.6 | 68.7 | 0.179 | -0.0014 | 0.079 |
| obs | bic | mar (WSP) | meg | 16,834 | 656.1 | 100.0 | 70.8 | 0.171 | -0.0015 | 0.025 |
| obs | bic | mar (WSP) | mos | 11,181 | 559.9 | 99.8 | 68.6 | 0.186 | -0.0020 | 0.048 |
| mar (WIO) | mar (Java) | luz | meg | 12,097 | 864.9 | 23.7 | 28.0 | -0.083 | 0.0002 | 0.319 |
| mar (WIO) | mar (Java) | luz | mos | 9,190 | 627.4 | 21.8 | 24.7 | -0.063 | 0.0004 | 0.350 |
| mar (WIO) | mar (SCS) | luz | meg | 18,097 | 735.4 | 48.7 | 63.9 | -0.135 | 0.0006 | 0.256 |
| mar (WIO) | mar (SCS) | luz | mos | 13,755 | 538.9 | 43.0 | 49.9 | -0.074 | 0.0005 | 0.329 |
| mar (WIO) | mar (WSP) | luz | meg | 15,210 | 842.0 | 27.2 | 78.1 | -0.483 | 0.0033 | 0.025 |
| mar (WIO) | mar (WSP) | luz | mos | 12,015 | 612.6 | 26.2 | 41.9 | -0.232 | 0.0012 | 0.238 |
| mar (WSP) | mar (WIO) | luz | meg | 15,210 | 842.0 | 78.1 | 27.2 | 0.483 | -0.0033 | 0.023[1] |
| mar (WSP) | mar (WIO) | luz | mos | 12,015 | 612.6 | 41.9 | 26.2 | 0.232 | -0.0012 | 0.218 |
| mar (WSP) | mar (Java) | luz | meg | 15,119 | 831.1 | 76.9 | 30.4 | 0.434 | -0.0034 | 0.021[1] |
| mar (WSP) | mar (Java) | luz | mos | 11,976 | 605.8 | 41.6 | 28.7 | 0.183 | -0.0010 | 0.253 |
| mar (WSP) | mar (SCS) | luz | meg | 20,938 | 704.2 | 101.3 | 65.7 | 0.213 | -0.0017 | 0.098 |
| mar (WSP) | mar (SCS) | luz | mos | 16,383 | 517.6 | 62.6 | 53.7 | 0.076 | -0.0003 | 0.361 |

[1]Note that the support for introgression between *A. luzonensis* and the *A. marmorata* populations from the western Indian Ocean and Java is not robust to outgroup choice and no longer significant after correcting for multiple tests.

#### References

Alexander DH, Novembre J, Lange K (2009) Fast model-based estimation of ancestry in unrelated individuals. *Genome Res.*, **19**, 1655–1664.

Benton MJ, Donoghue MJ, Asher RJ *et al.* (2015) Constraints on the timescale of animal evolutionary history. *Palaeontologia Electronica*, **18.1.1FC**, 1–106.

Carnevale G, Bannikov AF, Marramà G, Tyler JC, Zorzin R (2014) 5. The Pesciara-Monte Postale Fossil-Lagerstätte: 2. Fishes and other vertebrates. In: *The Bolca Fossil-Lagerstätten: A window into the Eocene World* (eds. Papazzoni CA, Giusberti L, Carnevale G *et al.*), pp. 37–63. Società Paleontologica Italiana.

Dela Pierre F, Bernardi E, Cavagna S *et al.* (2011) The record of the Messinian salinity crisis in the Tertiary Piedmont Basin (NW Italy): The Alba section revisited. *Palaeogeogr. Palaeoclimatol. Palaeoecol.*, **310**, 238–255.

Durand EY, Patterson N, Reich D, Slatkin M (2011) Testing for ancient admixture between closely related populations. *Mol. Biol. Evol.*, **28**, 2239–2252.

Gagnaire PA, Minegishi Y, Zenboudji S *et al.* (2011) Within-population structure highlighted by differential introgression across semipermeable barriers to gene flow in *Anguilla marmorata*. *Evolution*, **65**, 3413–3427.

Green RE, Krause J, Briggs AW *et al.* (2010) A draft sequence of the Neandertal genome. *Science*, **328**, 710–722.

Hasegawa M, Kishino H, Yano T (1985) Dating of the human-ape splitting by a molecular clock of mitochondrial DNA. *Journal of Molecular Evolution*, **22**, 160–174.

Henkel CV, Burgerhout E, de Wijze DL *et al.* (2012a) Primitive duplicate Hox clusters in the European eel's genome. *PLOS ONE*, **7**, e32231.

Henkel CV, Dirks RP, de Wijze DL *et al.* (2012b) First draft genome sequence of the Japanese eel, *Anguilla japonica*. *Gene*, **511**, 195–201.

Ishikawa S, Tsukamoto K, Nishida M (2004) Genetic evidence for multiple geographic populations of the giant mottled eel *Anguilla marmorata* in the Pacific and Indian oceans. *Ichthyol. Res.*, **51**, 343–353.

Jacobsen MW, Pujolar JM, Gilbert MTP *et al.* (2014) Speciation and demographic history of Atlantic eels (*Anguilla anguilla* and *A. rostrata*) revealed by mitogenome sequencing. *Heredity*, **113**, 432–442.

Jansen HJ, Liem M, Jong-Raadsen SA *et al.* (2017) Rapid de novo assembly of the European eel genome from nanopore sequencing reads. *Sci. Rep.*, **7**, 7213.

Kircher M, Sawyer S, Meyer M (2012) Double indexing overcomes inaccuracies in multiplex sequencing on the Illumina platform. *Nucleic Acids Res.*, **40**, e3–e3.

Lawson DJ, Hellenthal G, Myers S, Falush D (2012) Inference of population structure using dense haplotype data. *PLoS Genet.*, **8**, e1002453.

Malinsky M, Trucchi E, Lawson DJ, Falush D (2018) RADpainter and fineRADstructure: Population Inference from RADseq Data. *Mol. Biol. Evol.*, **35**, 1284–1290.

Matschiner M (2016) Fitchi: haplotype genealogy graphs based on the Fitch algorithm. *Bioinformatics*, **32**, 1250–1252.

Matschiner M, Musilová Z, Barth JMI *et al.* (2017) Bayesian phylogenetic estimation of clade ages supports trans-Atlantic dispersal of cichlid fishes. *Syst. Biol.*, **66**, 3–22.

Meyer BS, Matschiner M, Salzburger W (2017) Disentangling incomplete lineage sorting and introgression to refine species-tree estimates for Lake Tanganyika cichlid fishes. *Syst. Biol.*, **66**, 531–550.

Minegishi Y, Aoyama J, Tsukamoto K (2008) Multiple population structure of the giant mottled eel, Anguilla marmorata. *Mol. Ecol.*, **17**, 3109–3122.

Minh BQ, Hahn MW, Lanfear R (2018) New methods to calculate concordance factors for phylogenomic datasets. *bioRxiv*. Doi:10.1101/487801.

Musilova Z, Cortesi F, Matschiner M *et al.* (2019) Vision using multiple distinct rod opsins in deep-sea fishes. *Science*, **364**, 588–592.

Nguyen LT, Schmidt HA, Von Haeseler A, Minh BQ (2015) IQ-TREE: A fast and effective stochastic algorithm for estimating maximum-likelihood phylogenies. *Mol. Biol. Evol.*, **32**, 268–274.

Nielsen R, Beaumont MA (2009) Statistical inferences in phylogeography. *Mol. Ecol.*, **18**, 1034–1047.

Patterson C (1993) Osteichthyes: Teleostei. In: *The fossil record 2*, pp. 621–656. Chapman & Hall, London, UK.

Patterson N, Price AL, Reich D (2006) Population structure and eigenanalysis. *PLoS Genet.*, **2**, e190.

Peterson BK, Weber JN, Kay EH, Fisher HS, Hoekstra HE (2012) Double digest RADseq: An inexpensive method for de novo SNP discovery and genotyping in model and non-model species. *PLoS ONE*, **7**, e37135.

Rabosky DL, Chang J, Title PO *et al.* (2018) An inverse latitudinal gradient in speciation rate for marine fishes. *Nature*, **559**, 392–395.

Reich D, Thangaraj K, Patterson N, Price AL, Singh L (2009) Reconstructing Indian population history. *Nature*, **461**, 489–494.

Rohland N, Reich D (2012) Cost-effective, high-throughput DNA sequencing libraries for multiplexed target capture. *Genome Res.*, **22**, 939–946.

Schabetsberger R, Økland F, Kalfatak D *et al.* (2015) Genetic and migratory evidence for sympatric spawning of tropical Pacific eels from Vanuatu. *Mar. Ecol. Prog. Ser.*, **521**, 171–187.

Schabetsberger R, Miller MJ, Dall'Olmo G *et al.* (2016) Hydrographic features of anguillid spawning areas: potential signposts for migrating eels. *Mar. Ecol. Prog. Ser.*, **554**, 141–155.

Sinha R, Stanley G, Gulati GS *et al.* (2018) Index switching causes "spreading-of-signal" among multiplexed samples in Illumina HiSeq 4000 DNA sequencing. *bioRxiv.* Doi:10.1101/125724.

Stamatakis A (2014) RAxML version 8: a tool for phylogenetic analysis and post-analysis of large phylogenies. *Bioinformatics*, **30**, 1312–1313.

Tavaré S (1986) Some probabilistic and statistical problems in the analysis of DNA sequences. *Lectures on Mathematics in the Life Sciences*, **17**, 57–86.

Watanabe S, Aoyama J, Miller MJ *et al.* (2008) Evidence of population structure in the giant mottled eel, *Anguilla marmorata*, using total number of vertebrae. *Copeia*, **2008**, 680–688.

Watanabe S, Miller MJ, Aoyama J, Tsukamoto K (2009) Morphological and meristic evaluation of the population structure of *Anguilla marmorata* across its range. *J. Fish Biol.*, **74**, 2069–2093.

Yule GU (1925) A mathematical theory of evolution, based on the conclusions of Dr. J. C. Willis, F.R.S. *Phil. Trans. R. Soc. B*, **213**, 21–87.
